# Large scale genomic rearrangements in selected *Arabidopsis thaliana* T-DNA lines are caused by T-DNA insertion mutagenesis

**DOI:** 10.1101/2021.03.03.433755

**Authors:** Boas Pucker, Nils Kleinbölting, Bernd Weisshaar

## Abstract

**Background:** Experimental proof of gene function assignments in plants is based on mutant analyses. T-DNA insertion lines provided an invaluable resource of mutants and enabled systematic reverse genetics-based investigation of the functions of *Arabidopsis thaliana* genes during the last decades.

**Results:** We sequenced the genomes of 14 *A. thaliana* GABI-Kat T-DNA insertion lines, which eluded flanking sequence tag-based attempts to characterize their insertion loci, with Oxford Nanopore Technologies (ONT) long reads. Complex T-DNA insertions were resolved and 11 previously unknown T-DNA loci identified, resulting in about 2 T-DNA insertions per line and suggesting that this number was previously underestimated. T-DNA mutagenesis caused fusions of chromosomes along with compensating translocations to keep the gene set complete throughout meiosis. Also, an inverted duplication of 800 kbp was detected. About 10% of GABI-Kat lines might be affected by chromosomal rearrangements, some of which do not involve T-DNA. Local assembly of selected reads was shown to be a computationally effective method to resolve the structure of T-DNA insertion loci. We developed an automated workflow to support investigation of long read data from T-DNA insertion lines. All steps from DNA extraction to assembly of T-DNA loci can be completed within days.

**Conclusion:** Long read sequencing was demonstrated to be an effective way to resolve complex T-DNA insertions and chromosome fusions. Many T-DNA insertions comprise not just a single T-DNA, but complex arrays of multiple T-DNAs. It is becoming obvious that T-DNA insertion alleles must be characterized by exact identification of both T-DNA::genome junctions to generate clear genotype-to-phenotype relations.

## Background

T-DNA insertion lines contributed substantially to the high-value knowledge about the functions of plant genes. This knowledge has been produced by the plant research community on the basis of gene structures predicted from genome sequences. T-DNA insertional mutagenesis emerged as an effective mechanism for the generation of knock-out alleles for use in reverse genetics and targeted gene function search [1, 2]. Since targeted integration of DNA into plant genomes via homologous recombination was at least technically challenging [3], large collections of sequence-indexed T-DNA integration lines with random insertion sites were used to provide knock-out alleles for the majority of genes [4]. Knowledge about the inserted sequences is an advantage over other mutagenesis methods, because localization of the insertion within the mutagenized genome based on the generation of flanking sequence tags (FSTs) is possible [5, 6]. CRISPR/Cas technology now offers technically feasible alternatives for access to mutant alleles for reverse genetics [7]. However, thousands of T-DNA insertion mutants have been characterized and represent today the main or reference mutant allele for (lack of) a given gene function.

*Agrobacterium tumefaciens* is a Gram-negative soil bacterium with the ability to transfer DNA into plant cells and to integrate this T-DNA stably and at random positions into the nuclear genome [8, 9]. The tumor inducing (Ti) plasmid, which is naturally occurring in *Agrobacteria* and enables them to induce the formation of crown galls in plants, contains the T-DNA that is transferred into plant cells [10]. The T-DNA is enclosed by 25 bp long imperfect repeats that were designated left (LB) and right border (RB) [9]. The T-DNA sequence between LB and RB can be modified to contain resistance genes for selection of successfully transformed plants [11]. T-DNAs from optimized binary plasmids are transformed into *A. thaliana* plants via floral dip to generate stable lines [12]. T-DNA transfer into the nucleus of a plant cell is supported by several VIR proteins which are, in the biotechnologically optimized system, encoded on a separate helper plasmid. It is assumed that host proteins are responsible for integration of the T-DNA into the genome, most likely as a DNA double strand into a double strand break (DSB) using host DNA repair pathways and DNA polymerase theta [9, 13, 14]. T-DNA integration resembles DNA break repair through non-homologous end-joining (NHEJ) or microhomology-mediated end-joining (MMEJ) and is often accompanied by the presence of filler DNA or microhomology at both T-DNA::genome junctions [9, 13]. Chromosomal inversions and translocations are commonly associated with T-DNA insertions in numerous plant species, not only in *A. thaliana* [15–21], suggesting that often more than just one DSB is associated with T-DNA integration [9].

The most important collections of T-DNA lines for the model plant *Arabidopsis thaliana* are SALK (150,000 lines) [6], GABI-Kat (92,000 lines) [22, 23], SAIL (54,000 lines) [24], and WISC (60,000 lines) [25]. In total, over 700,000 insertion lines have been constructed [4]. GABI-Kat lines were generated through the integration of a T-DNA harboring a sulfadiazine resistance gene for selection of transformed lines [22]. Additionally, the T-DNA contains a 35S promoter at RB causing transcriptional up-regulation of plant genes next to the integration site if the right part of the T-DNA next to RB stays intact during integration [1]. Integration sites were predicted based on FSTs and allowed access to knock-out alleles of numerous genes. At GABI-Kat, T-DNA insertion alleles were confirmed by an additional “confirmation PCR” using DNA from the T2 generation [26] prior to the release of a mutant line and donation of the line to the Nottingham Arabidopsis Stock Centre (NASC). Researchers could identify available T-DNA insertion lines via SimpleSearch on the GABI-Kat website [27]. Since 2017, SimpleSearch uses Araport11 annotation data [28]. Araport11 is based on the *A. thaliana* Col-0 reference genome sequence from TAIR9 which includes about 96 annotated gaps filled with Ns as a sign for unknown sequences [29], among them the centromers and several gaps in the pericentromeric regions.

The prediction of integration sites based on bioinformatic evaluations using FST data does often not reveal the complete picture. Insertions might be masked from FST predictions due to truncated borders [13], because of repetitive sequences or paralogous regions in the genome [30], or even lack of the true insertion site in the reference sequence used for FST mapping [31, 32]. Also, confirmation by sequencing an amplicon that spans the predicted insertion site at one of the two expected T-DNA::genome junctions is not fully informative. Deletions and target site duplications at the integration site can occur and can only be detected by examining both borders of the inserted T-DNA [13]. In addition, several studies have reported more complex T-DNA insertions in association with large deletions, insertions, inversions or even chromosomal translocations [13, 18, 19, 21, 33, 34]. A relation of T-DNA insertion mutagenesis to chromosomal translocations has not only been reported for *A. thaliana*. Also for transgenic rice (*Oryza sativa*) [35] and transgenic birch (*Betula platyphylla* x *B. pendula*) [36] changes at the genome level have been reported. Binary vector backbone (BVB) sequences have been detected at insertion sites [19, 37] as well as fragments of *A. tumefaciens* chromosomal DNA [38]. Recombination between two T-DNA loci was described as a mechanism for deletion of an enclosed genomic fragment [39].

Chromosomal rearrangements can occur during the repair of DSBs, e.g. via microhomology-mediated end joining or non-allelic homologous recombination [reviewed by 40, 41]. Evidently, regions with high sequence similarity like duplications are especially prone to chromosomal rearrangements [40].

While usually one T-DNA locus per line was identified by FSTs, the number of T-DNA insertion loci per line is usually higher. For GABI-Kat, it was estimated that about 50% of all lines (12,018 of 21,049 tested, according to numbers from the end of 2019) display a single insertion locus. This estimation is based on segregation analyses using sulfadiazine resistance as a selection marker [22]. Other insertion mutant collections report similar numbers [4]. The average number of T-DNA insertions per line was reported to be about 1.5, but this is probably an underestimation since the kanamycin and BASTA selection marker genes applied to determine the numbers are known to be silenced quite often [4]. For these reasons, it is required that insertion mutants (similar to mutants created by e.g. chemical mutagenesis) are backcrossed to wild type prior to phenotyping a homozygous line.

The FSTs produced for the different mutant populations by individual PCR and Sanger-sequencing allowed usually access to a single T-DNA insertion locus per line, although for GABI-Kat there are several examples with up to three confirmed insertion loci based on FST data (e.g. line GK-011F01, see [27]). This leaves a significant potential of undiscovered T-DNA insertions in lines already available at the stock centers, which has been exploited by the group of Joe Ecker by applying TDNA-Seq (Illumina technology) to the SALK and a part of the GABI-Kat mutant populations (unpublished, all data available from GenBank/ENA). Essentially the same technology has subsequently been used to set up a sequence indexed insertion mutant library of *Chlamydomonas reinhardtii* [42]. With the fast development of new DNA sequencing technologies, the comprehensive characterization of T-DNA insertion lines comes into reach.

Several studies already harnessed high throughput sequencing technologies to investigate T-DNA insertion and other mutant lines [43, 44]. Oxford Nanopore Technologies (ONT) provides a cost-effective and fast approach to study *A. thaliana* genomes, since a single MinION/GridION Flow Cell delivers sufficient data to assemble one genotype [45]. Here, we present a method to fully characterize T-DNA insertion loci and additional genomic changes of T-DNA insertion lines through ONT long read sequencing. We selected 14 lines that contain confirmed T-DNA insertion alleles. The first border or first T-DNA::genome junction has been confirmed by sequencing an amplicon across one junction, but the insertion eluded characterization of the second T-DNA::genome junction. We refer to the additional T-DNA::genome junction that is expected to exist after confirmation of one T-DNA::genome junction as “2^nd^ border”. Within this set of lines that is enriched for unusual relations between the two expected T-DNA::genome junctions, we detected several chromosome fragment or chromosome arm translocations, a duplication of 800 kbp and also an insertion of DNA from the chloroplast (plastome), all related to T-DNA insertion events. The results clearly demonstrate the importance of characterizing both T-DNA::genome junctions for reliable selection of suitable alleles for setting up genotype/phenotype relations for gene function search. In parallel to data evaluation, we created an automated workflow to support long-read-based analyses of T-DNA insertion lines and alleles.

## Results

In total, 14 GABI-Kat T-DNA insertion lines (Table 1, Additional file 1) were selected for genomic analysis via ONT long read sequencing. This set of lines was selected based on prior knowledge which indicated that the insertion locus addressed in the respective line was potentially unusual. The specific feature used for selection was the (negative) observation that creation of confirmation amplicons which span the T-DNA::genome junction failed for one of the two junctions, operationally that means that the 2^nd^ border could not be confirmed. T-DNA insertion loci in the selected lines were assessed by *de novo* assembly of the 14 individual genome sequences, and by a computationally more effective local assembly of selected reads.

**Table 1:** Key findings summary of ONT-sequenced GABI-Kat T-DNA insertion lines.

| Line ID<sup>a</sup> | number of insertions |  |  |  | Summary of observation |
| --- | --- | --- | --- | --- | --- |
|  | FST | PCR- | total | ONT |  |
|  | pred. | conf. | found | found |  |
| GK-038B07 | 2 | 0 (1)<sup>b</sup> | 4<sup>c</sup> | 3 | FST-predicted insertion in Chr5 is part of a fusion of Chr3 and Chr5, 2 T-DNA arrays detected at translocation fusion points of which one contains in addition an inversion of ~2 Mbp fused with another T-DNA array, additional insertion of a complex T-DNA array in Chr1. |
| GK-089D12 | 1 | 1 | 2 | 1 | Fusion of Chr3 and Chr5, FSTs are derived from the single DNA::genome junction that contains LB, both translocation fusions contain mostly canonical T-DNAs, failure to confirm 2<sup>nd</sup> T-DNA::genome junction explained by shortened T-DNA. |
| GK-430F05 | 1 | 1 | 3 | 2 | Three T-DNA insertion sites, two complex T-DNA arrays with one containing more than 5 kbp BVB<sup>d</sup>, the confirmed T-DNA insertion in Chr3 is complex, no theoretical explanation for failure of confirmation PCR at 2<sup>nd</sup> DNA::genome junction, one insertion in the pericentromeric region of (probably) Chr4 containing a rearranged RB region. |
| GK-654A12<sup>d</sup> | 1 | 1 | 2<sup>c</sup> | 1 | Fusion of Chr1 and Chr4 with a complex T-DNA array that also contains BVB, Chr1-part of the fusion predicted by FST, the compensating fusion of Chr4 and Chr1 does not contain T-DNA at the translocation fusion point, translocation explains failure to confirm 2<sup>nd</sup> T-DNA::genome junction, additional insertion of a T-DNA array in Chr2 with a 162 bp duplicated inversion at the integration site. |
| GK-767D12 | 1 | 1 | 2 | 1 | Large segmental duplication and inversion at the predicted insertion site, a long T-DNA array also containing BVB present at the southern end of the segmental duplication, inversion explains failure to confirm 2<sup>nd</sup> T-DNA::genome junction, the northern fusion point of the inverted segmental duplication does not contain T-DNA, another canonical T-DNA insertion in Chr2 but with 4 Mbp distance. |
| GK-909H04 | 1 | 1 | 3 | 2 | The predicted T-DNA insertion contains an inverted duplication of 20 kbp at the integration site explains why the 2<sup>nd</sup> T-DNA::genome junction could not be confirmed, integration of 652 bp derived from the plastome at the northern fusion point of the duplication but no T-DNA, two additional canonical T-DNA insertions in Chr1 and Chr2. |
| GK-947B06 | 1 | 1 | 2 | 1 | Complex T-DNA array insertion of 3 T-DNAs at the predicted insertion site in Chr1, no theoretical explanation for failure of the confirmation PCR at the 2<sup>nd</sup> DNA::genome junction, additional complex insertion containing 2 T-DNAs and BVB on Chr2. |

<sup>a</sup> Lines with newly detected insertions, all lines are listed in Additional file 2.
<sup>b</sup> The T-DNA insertion used for line selection was a false positive case not detected by ONT seq.
<sup>c</sup> One locus in a line with a chromosome translocation causes the FSTs from this single locus to map to two places in the reference sequence. These two places might also be the source of potentially existing FST from the compensating chromosome fusion. If the compensating chromosome fusion does not contain a T-DNA, the number of insertions is lower than expected.
<sup>d</sup> BVB, binary vector backbone

A tool designated “loreta” (<u>lo</u>ng <u>re</u>ad-based t-DNA <u>a</u>nalysis) has been developed during the analyses and might be helpful for similar studies (see methods for details). The results of both approaches demonstrate that a full *de novo* assembly is not always required if only certain regions in the genome are of interest. The 14 GABI-Kat lines harbor a total of 26 T-DNA insertions resulting in an average of 1.86 insertions per line. A total of 11 insertion loci detected in seven of 14 lines were not revealed by previous FST based attempts to detect T-DNA insertions (Table 1, Additional file 2 and 3). In case of GK-038B07, the lack of re-detection of the expected insertion allele of At4g19510 was explained by a PCR template contamination during the initial confirmation, the line that contains the real insertion (source of the contamination) is most probably GK-159D11. A DNA template contamination is also likely in case of GK-040A12 where the insertion allele of At1g52720 expected from the confirmation was also not found in the ONT data. At least, the error detected fits to the selection criteria, because the 2^nd^ T-DNA::genome junction can obviously not be detected if the T-DNA insertion allele as such is not present in the line.

Six of the 14 lines studied by ONT whole genome sequencing displayed chromosomal rearrangements. To visualize these results, we created ideograms of the five *A. thaliana* chromosomes with a color code for each chromosome. The colors allow to visually perceive information on chromosome arm translocations, and the changeover points indicate presence or absence of T-DNA sequences.

### Chromosome fusions

In four lines, fusions of different chromosomes were detected. These fusions result from chromosome arm translocations which were, in all four cases, compensated within the line by reciprocal translocations.

The T-DNA insertion on chromosome 5 (Chr5) of GK-038B07 is part of a complex chromosome arm translocation (Fig. 1). A part of Chr5 is fused to Chr3, the replaced part of Chr3 is fused to an inversion of 2 Mbp on Chr5. This inversion contains T-DNAs at both ends, one of which is the insertion predicted by FSTs.

**Fig. 1:**
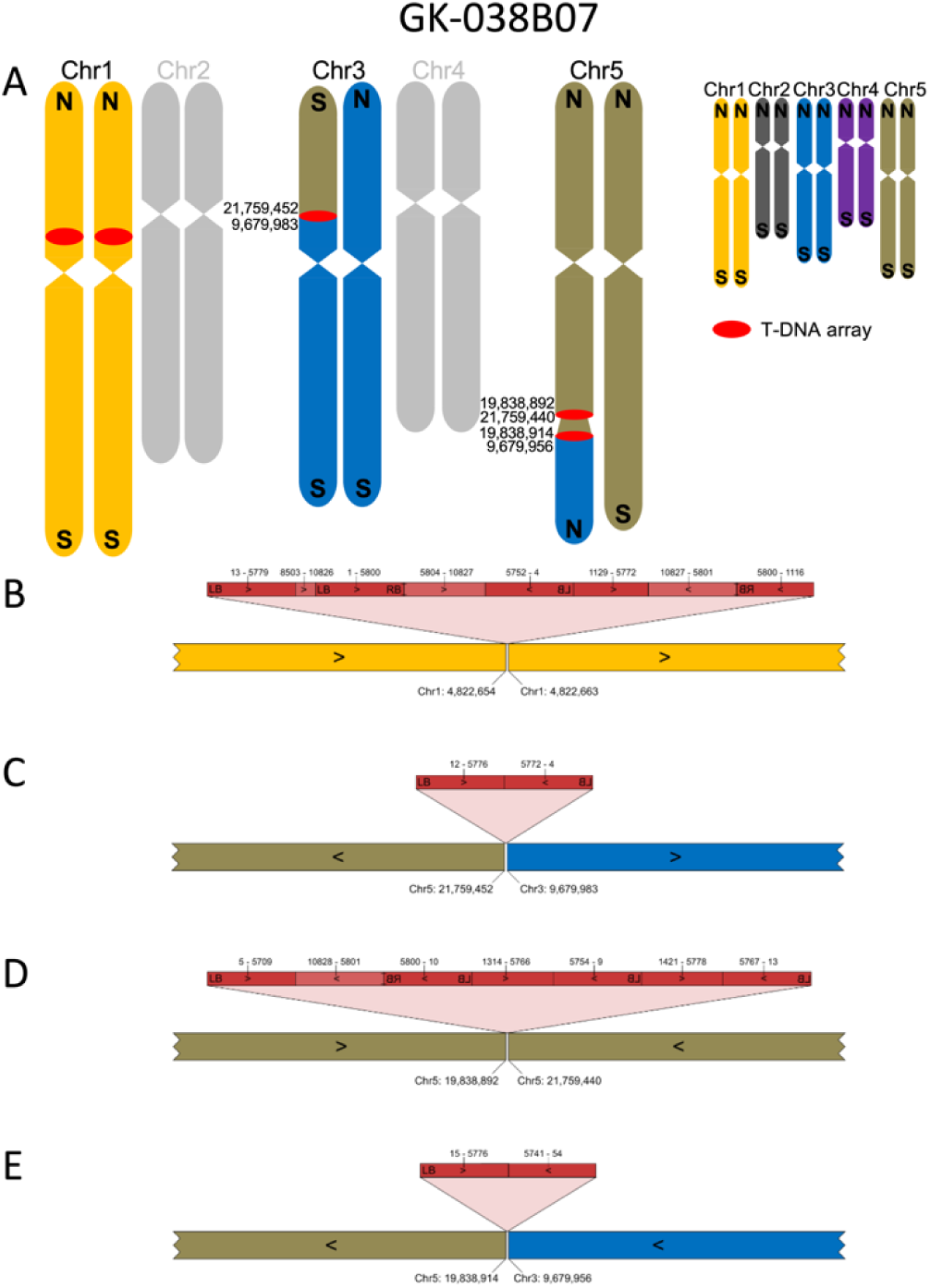
Organization of the nuclear genome of GK-038B07 with a focus on translocations, inversions and T-DNA structures. A: Ideograms of the chromosomes display the reciprocal fusion of Chr3 and Chr5 as well as a 2 Mbp inversion between two T-DNA arrays at the fusion sites; numbers indicate end points of pseudochromosome fragments according to TAIR9. Chromosomes depicted in grey match the Col-0 sequence. The insert displays the color codes used for all five chromosomes; N, northern end of chromosome; S, southern end of chromosome. B-E: Visualization of the four T-DNA insertion loci of GK-038B07 resolved by local assembly. LB and RB, T-DNA left and right border; dark red, *bona fide* T-DNA sequences located between the borders; light red, sequence parts from the binary vector backbone (BVB); numbers above the red bar indicate nucleotide positions with position 1 placed at the left end of LB in the binary vector which makes position 4 the start of the transferred DNA [13]; numbers below the colored bars indicate pseudochromosome positions according to TAIR9.

For line GK-082G09, two FST predictions had been generated and one FST lead to the prediction of an insertion at Chr3:6,597,745 which was confirmed by PCR. Confirmation of the expected corresponding 2^nd^ border failed. Another FST-based prediction at Chr5:23,076,864 was not addressed by PCR. ONT sequencing confirmed both predictions (Fig. 2). There is only one complex insertion consisting of multiple T-DNA copies and BVB in GK-082G09 that fuses the south of Chr5 to an about 6.6 Mbp long fragment from the north of Chr3 (Fig. 2C). This translocation is compensated by a fusion of the corresponding parts of both chromosomes without a T-DNA at the fusion point. The second fusion point of Chr3 and Chr5, that was detected in the *de novo* assembly of the genome sequence of GK-082G09, was validated by generating and sequencing a PCR amplicon spanning the translocation fusion point (Fig. 2B, see Additional file 1 for sequences/accession numbers and Additional file 4 for the primer sequences).

**Fig. 2:**
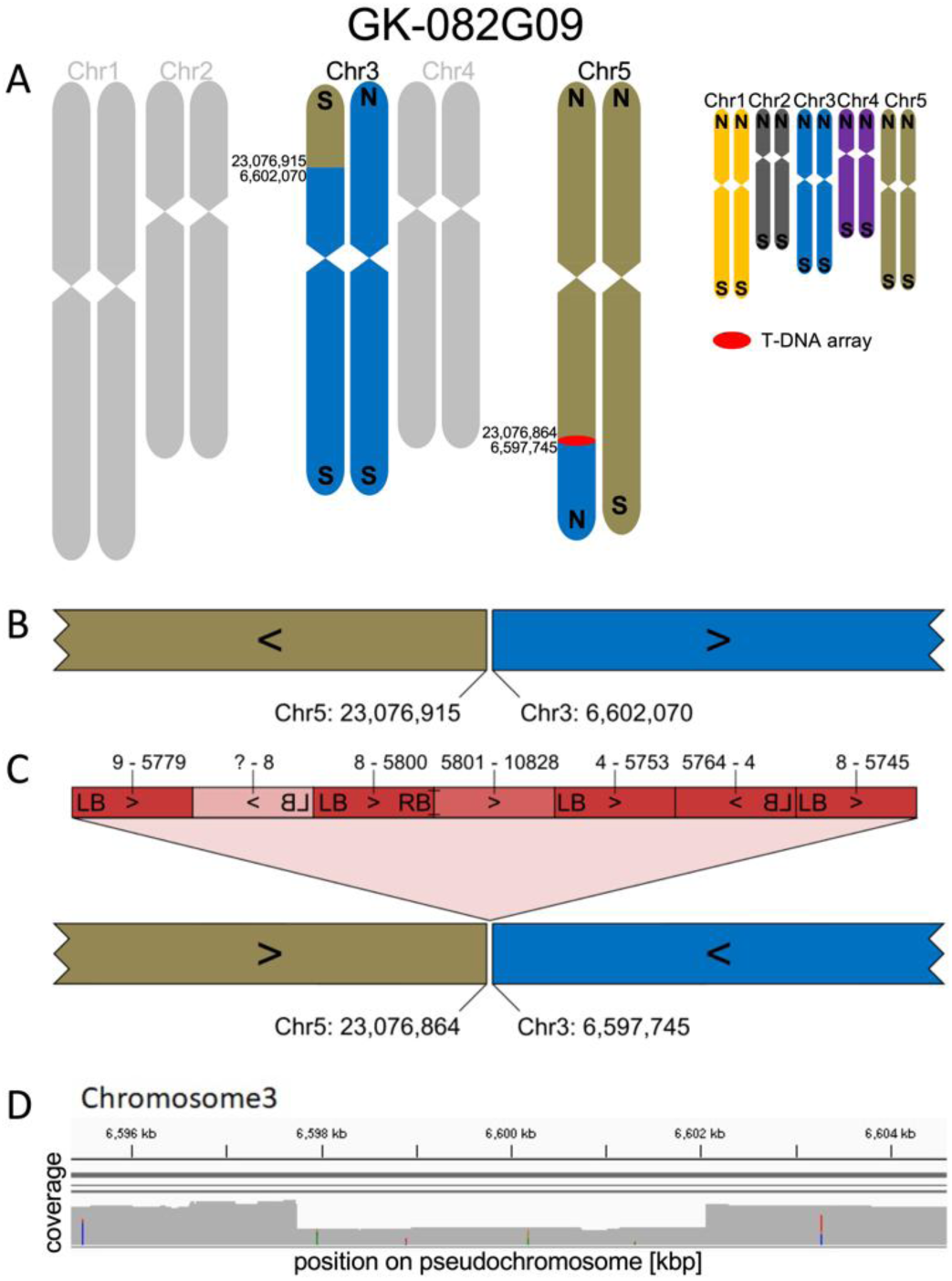
Organization of the nuclear genome of GK-082G09 with a focus on translocations, inversions and T-DNA structures. A-C: For a description of the figure elements see legend to Fig. 1. D: Read coverage depth analyses of the region of Chr3 that is involved in the fusions which confirms a deletion of about 4 kbp from Chr3. The reads that cover the deleted part were derived from the wild type allele present in the segregating population (see methods).

Line GK-089D12 harbors two T-DNA insertions (Fig. 3) and both were predicted by FSTs, one in Chr3 and one in Chr5. Since fragments of Chr3 and Chr5 are exchanged in a reciprocal way with no change in sequence direction (southern telomeres stay at the southern ends of the chromosomes), PCR confirmation would have usually resulted in “fully confirmed” insertion alleles. Only long read sequencing allowed to determine the involvement of translocations. The line was studied because the shortened T-DNA at 089D12-At5g51660-At3g63490 (Fig. 3B, see Additional file 3 for designations of insertions) caused failure of formation of the confirmation amplicon.

**Fig. 3:**
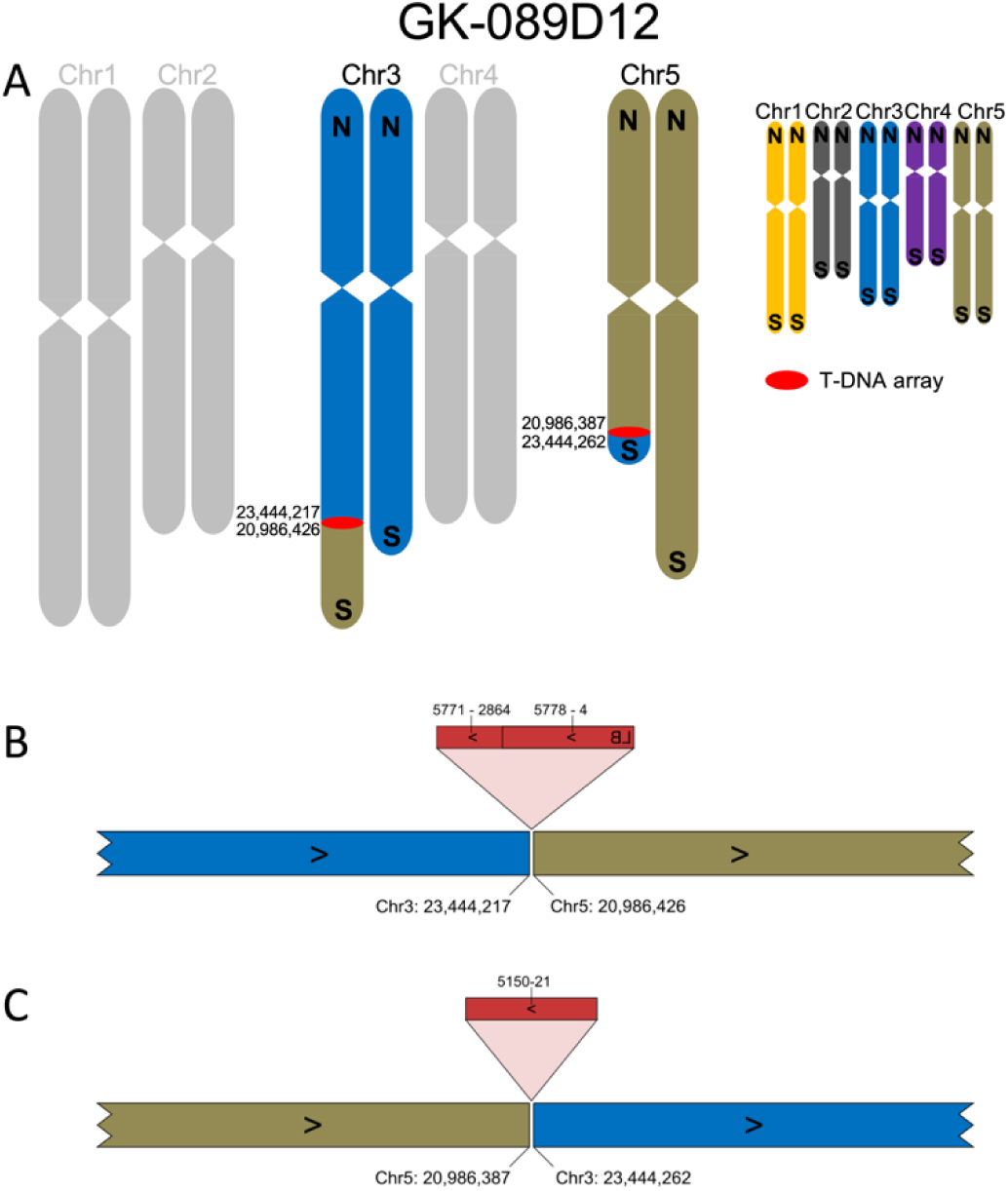
Organization of the nuclear genome of GK-089D12 with a focus on translocations, inversions and T-DNA structures. A-C: For a description of the figure elements see legend to Fig. 1.

FSTs from line GK-654A12 indicated a T-DNA insertion on Chr1. ONT sequencing revealed a translocation between Chr1 and Chr4 that explained failure to generate the confirmation amplicon at the 2^nd^ border (Fig. 4). The southern arms of Chr1 and Chr4 are exchanged, with a T-DNA array inserted at the fusion point of the new chromosome that contains CEN1 (centromere of Chr1). The fusion point of the new chromosome that contains CEN4 does not contain T-DNA sequences. Also this T-DNA-free fusion point (designated 654A12-FCAALL-0-At1g45688) was validated by generating and sequencing a PCR amplicon which spanned the fusion site (Fig. 4C, Additional files 1 and 4). The T-DNA array at 654A12-At1g45688-FCAALL contains BVB sequences, interestingly as an independent fragment and not in an arrangement that is similar to the binary plasmid construction which provided the T-DNA (Fig. 4B).

**Fig. 4:**
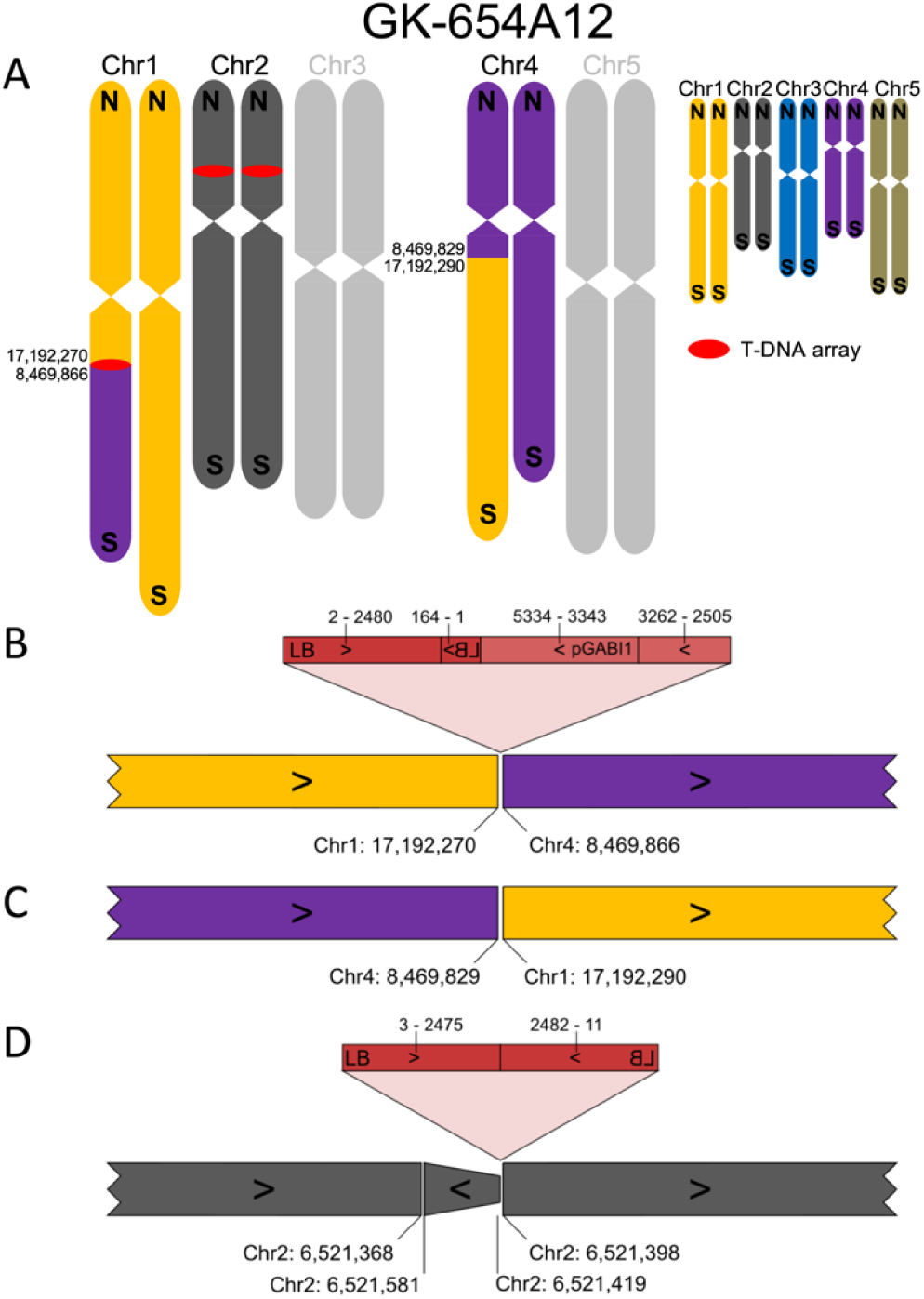
Organization of the nuclear genome of GK-654A12 with a focus on translocations, inversions and T-DNA structures. A-C: For a description of the figure elements see legend to Fig. 1. See Additional file 2 for an explanation of pGABI1. D: The T-DNA insertion in Chr2 is associated with a small duplicated inversion of about 160 bp as already described for a fraction of all T-DNA::genome junctions [13].

### Intrachromosomal rearrangements and a large duplication

For line GK-433E06 the FST data indicated four insertions, one T-DNA insertion (433E06-At1g73770-F9L1) at Chr1:27,742,275 has been confirmed by amplicon sequencing. ONT sequencing revealed an intrachromosomal translocation that exchanged the two telomeres of Chr1 together with about 5 Mbp DNA. The FSTs that indicated two T-DNA insertions in chromosome 1 were derived from one T-DNA array (Fig. 5). Once more, the compensating fusion point, designated 433E06-F9L1-0-At1g73770, does not contain T-DNA sequences which was validated by amplicon sequencing (Fig. 5C, Additional files 1 and 4).

**Fig. 5:**
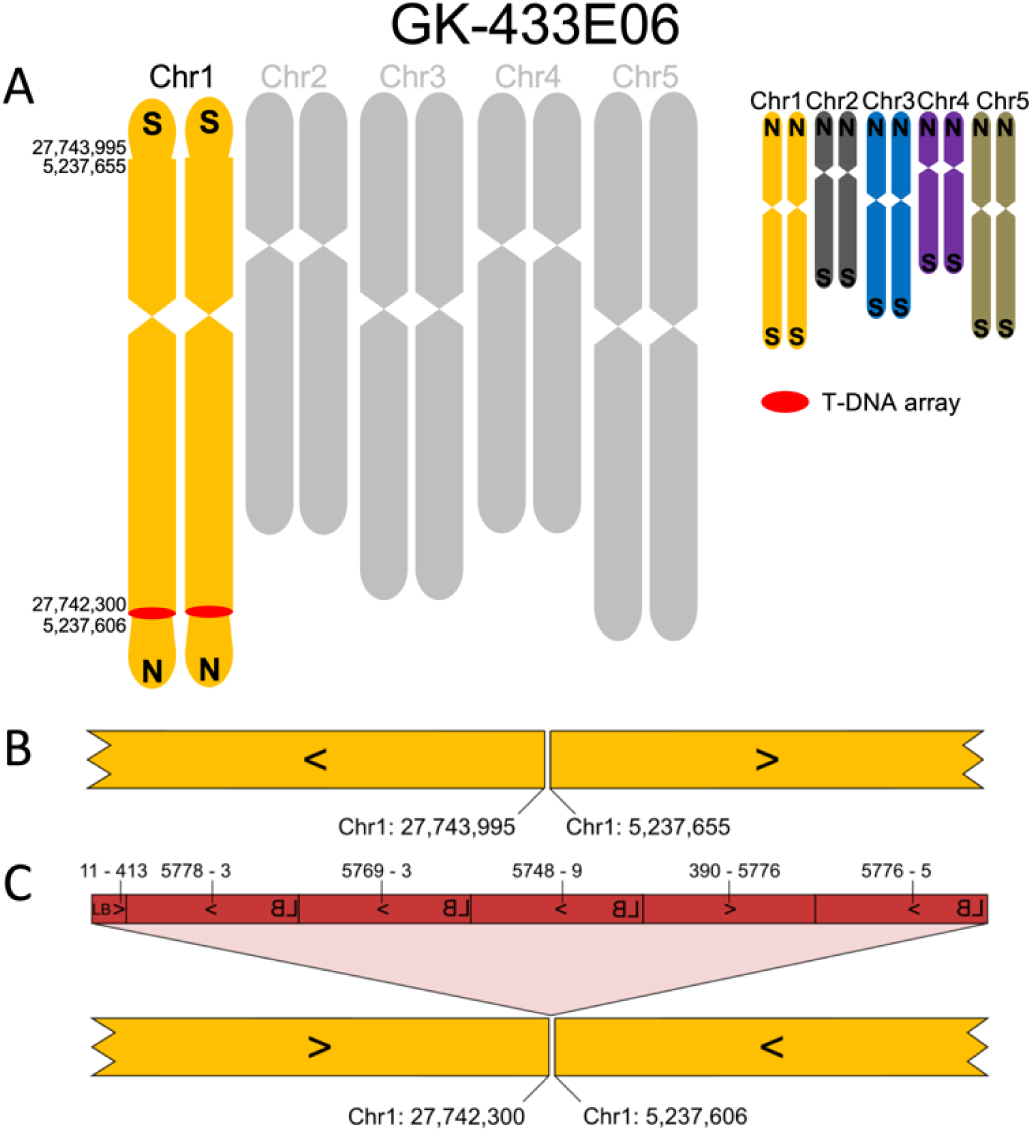
Organization of the nuclear genome of GK-433E06 with a focus on translocations, inversions and T-DNA structures. A-C: For a description of the figure elements see legend to Fig. 1.

In line GK-767D12, a large duplication of a part of Chr2 that covers about 800 kbp was detected (Fig. 6). The duplication is apparent from read coverage analyses (Fig. 6B) based on read mapping against the TAIR9 reference genome sequence (Col-0) which was performed for all lines studied (Additional file 5). The duplicated region is inserted in reverted orientation (inversion) next to the T-DNA insertion 767D12-At2g19210-At2g21385. This insertion was predicted by an FST at Chr2:8,338,072 and has been confirmed by PCR, the 2^nd^ border confirmation for the T-DNA insertion failed because of reversed orientation. The other end of the duplicated inversion of Chr2 is fused to Chr2:8,338,361 (designated 767D12-At2g21385-0-At2g19210) without T-DNA sequences (Fig. 6D). Also this T-DNA-free fusion point was validated by sequencing a PCR amplicon which spanned the fusion site (Additional files 1 and 4). The T-DNA array at 767D12-At2g19210-At2g21385 is the largest we detected in this study. It consists of 8 almost complete T-DNA copies arranged in diversified configurations and includes a large BVB fragment (Fig. 6C).

**Fig. 6:**
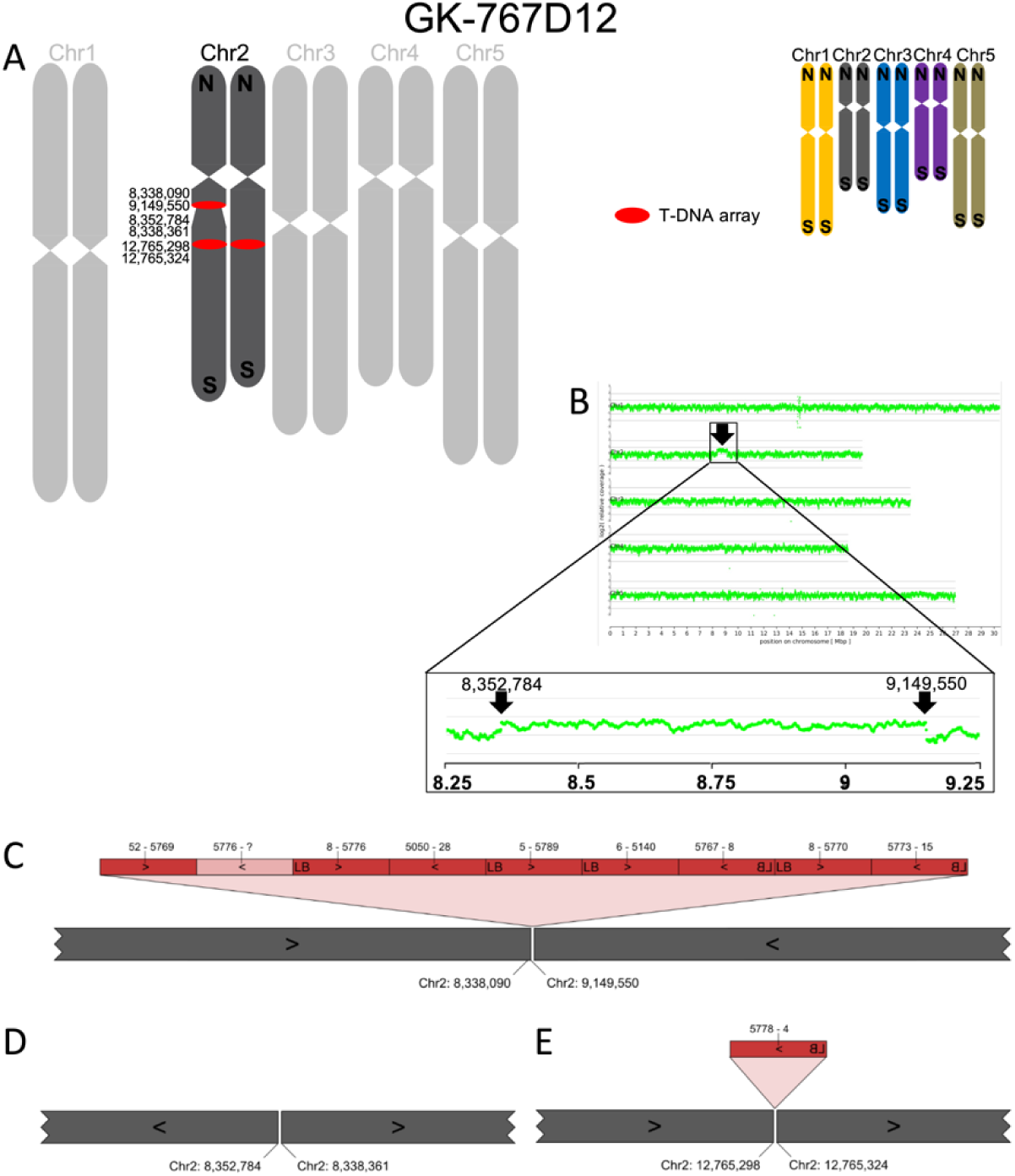
Organization of the nuclear genome of GK-767D12 with a focus on translocations, inversions and T-DNA structures. A-E: For a description of the figure elements see legend to Fig. 1. B: Results from a read coverage depth analysis are depicted that revealed a large duplication compared to the TAIR9 Col-0 reference sequence. We used read coverage depth data to decide for the selection of the zygosity of the insertions and rearrangements displayed for Chr2 in the ideograms in panel A.

FST analyses detected only one T-DNA insertion in line GK-909H04. This insertion, designated 909H04-At1g54390, had been confirmed by PCR but failed for the 2^nd^ border. ONT sequencing revealed an inverted duplication of about 20 kbp next to the T-DNA insertion site (Fig. 7). The fusion between this inverted duplication and the remaining part of Chr1 does not contain T-DNA sequences, but a 652 bp fragment derived from the plastome (Fig. 7C). The cpDNA insertion was validated by generating and sequencing a PCR amplicon spanning the insertion and both junctions to the genome (Additional files 1 and 4). ONT sequencing also revealed an additional insertion of a truncated T-DNA (909H04-At1g38212 at about 14.3 Mbp of Chr1) which is in the pericentromeric region not far from CEN1 (CEN1 is located at 15,086,046 to 15,087,045 and marked in the TAIR9 reference sequence by a gap of 1,000 Ns). Initial analyses indicated that this insertion might be associated with a deletion of about 45 kbp. However, the predicted deletion was not indicated by the read coverage depth analyses, and the region is rich in TEs (also At1g38212 is annotated as “transposable element gene”). We assembled a new genome sequence of the Col-0 wild type used at GABI-Kat (assembly designated Col-0_GKat-wt, see below) and studied the structure of 909H04-At1g38212 on the basis of this assembly. The results indicated that the deletion predicted on the basis of the TAIR9 assembly is a tandemly repeated sequence in TAIR9 which is differently represented in Col-0_GKat-wt (Additional file 6). The 3’-end of an example read from line GK-909H04 maps continuously to Col-0_GKat-wt and also to a sequence further downstream in TAIR9. The evidence collected clearly shows that there are only 13 bp deleted at the T-DNA insertion at 14.3 Mbp of Chr1 (Fig. 7D), and that the initially predicted deletion is caused by errors in the TAIR9 assembly in this pericentromeric region.

**Fig. 7:**
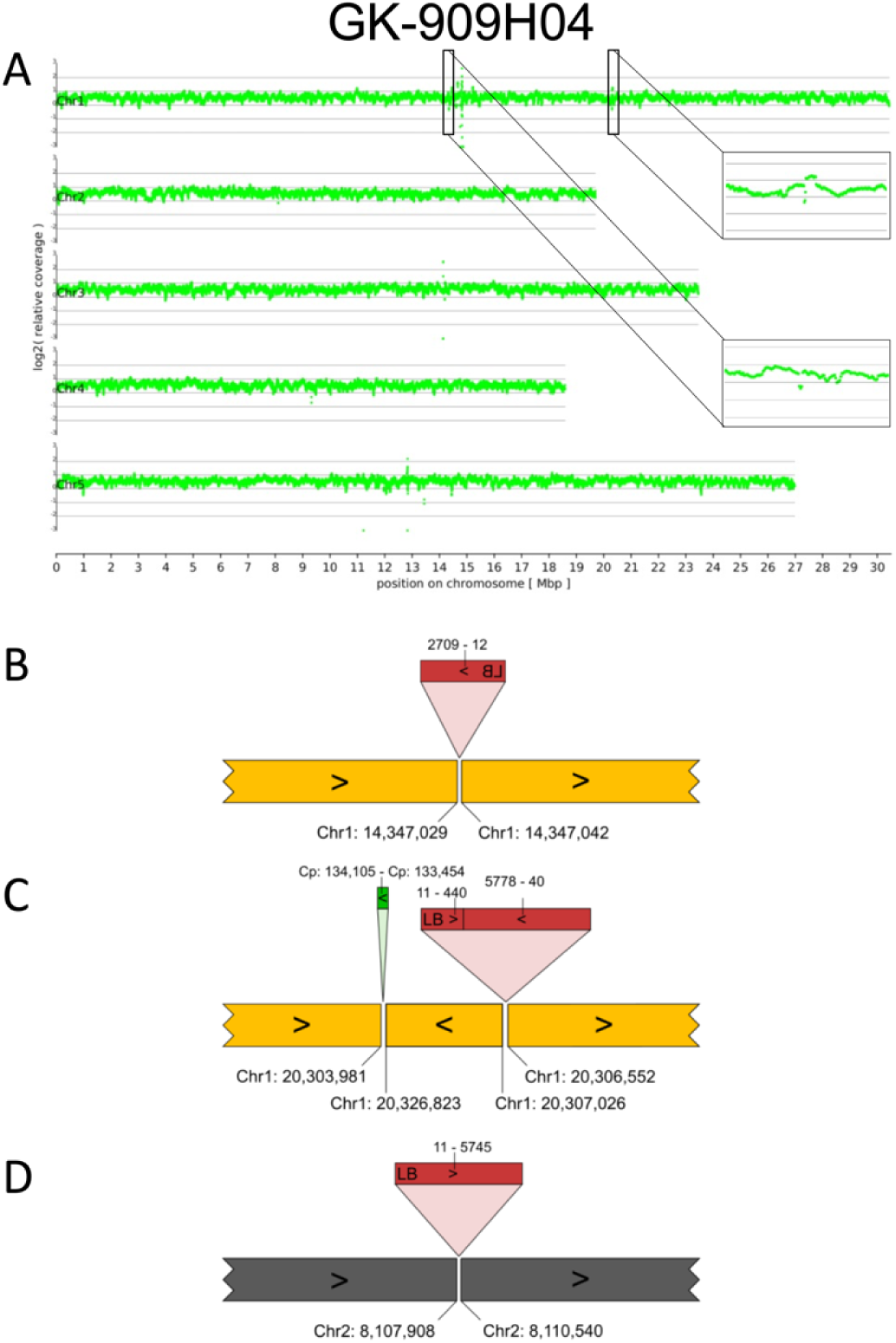
Organization of the nuclear genome of GK-909H04 with a focus on insertions and T-DNA structures. A-D: For a description of the figure elements see legend to Fig. 6. A: The read coverage depth plot includes two zoom-in enlargements of the regions at 14.3 and 20.3 Mbp of Chr1. These two display variable coverage in the region of the truncated T-DNA insertion 909H04-At1g38212 (see text), and increased coverage next to the T-DNA insertion 909H04-At1g54390. C: Duplicated inversion detected in the local assembly of GK-909H04 which is also supported by read coverage depicted in Panel A. Green block, sequence part from the plastome (cpDNA).

The six junctions (five sequences) that contained no T-DNA, three from compensating chromosome fusions, one from the 800 kbp inversion and two at both ends of the cpDNA insertion (see Additional file 3), were analyzed for specific features at the junctions. The observations made were fully in line with what has already been described for T-DNA insertion junctions: some short filler DNA and microhomology was found (Additional file 7). A visual overview over the T-DNA insertion structures of all 14 lines, including those not displaying chromosomal rearrangements, is presented in Additional file 8.

### Detection of novel T-DNA insertions and T-DNA array structures

As mentioned above, 11 T-DNA insertion loci were newly detected in 7 of 14 lines studied, indicating that these were missed by FST-based studies (Table 1, Additional file 2 and 3). The primer annealing sites for FST generation at LB seem to be present in all 11 T-DNA insertions only found by ONT sequencing. Analysis of the data on T-DNA::genome junctions summarized in Additional file 3 revealed that a majority of the T-DNA structures have LB sequences at both T-DNA::genome junctions (14 of 26). The bias for the T-DNA::genome junctions involving LB is increased by the fact that several of the RB junctions were truncated, and also by some other junctions which involve BVB sequences. True T-DNA::genome junctions involving intact RB were not in the dataset, and in 14 out of 26 cases an internal RB::RB fusion was detected.

While about 40% (8 to 10 of 26, depending on judgement of small discontinuous parts) of the insertions contain a single T-DNA copy (referred to here as “canonical” insertions), some of which even further truncated and shortened, there are often cases of complex arrays of T-DNA copies inserted as T-DNA arrays. We observed a wide variety of configurations of the individual T-DNA copies within the complex arrays. In six out the 26 cases BVB sequences were detected, in the case of 038B07-At1g14080, 082G09-At5g57020-At3g19080, 430F05-At4g23850 and 947B06-T7M7 even almost complete vector sequences.

### Sequence read quality decreased in T-DNA arrays

During the analyses of T-DNA insertion sequences, we frequently faced regions without sequence similarity to any sequence in the *A. thaliana* genome sequence, the sequence of the Ti-plasmid (T-DNA and BVB), or the *A. tumefaciens* genome sequence. Analysis of the read quality (Phred score) in these regions compared to other regions on the same read revealed a quality drop (Fig. 8).

**Fig. 8:**
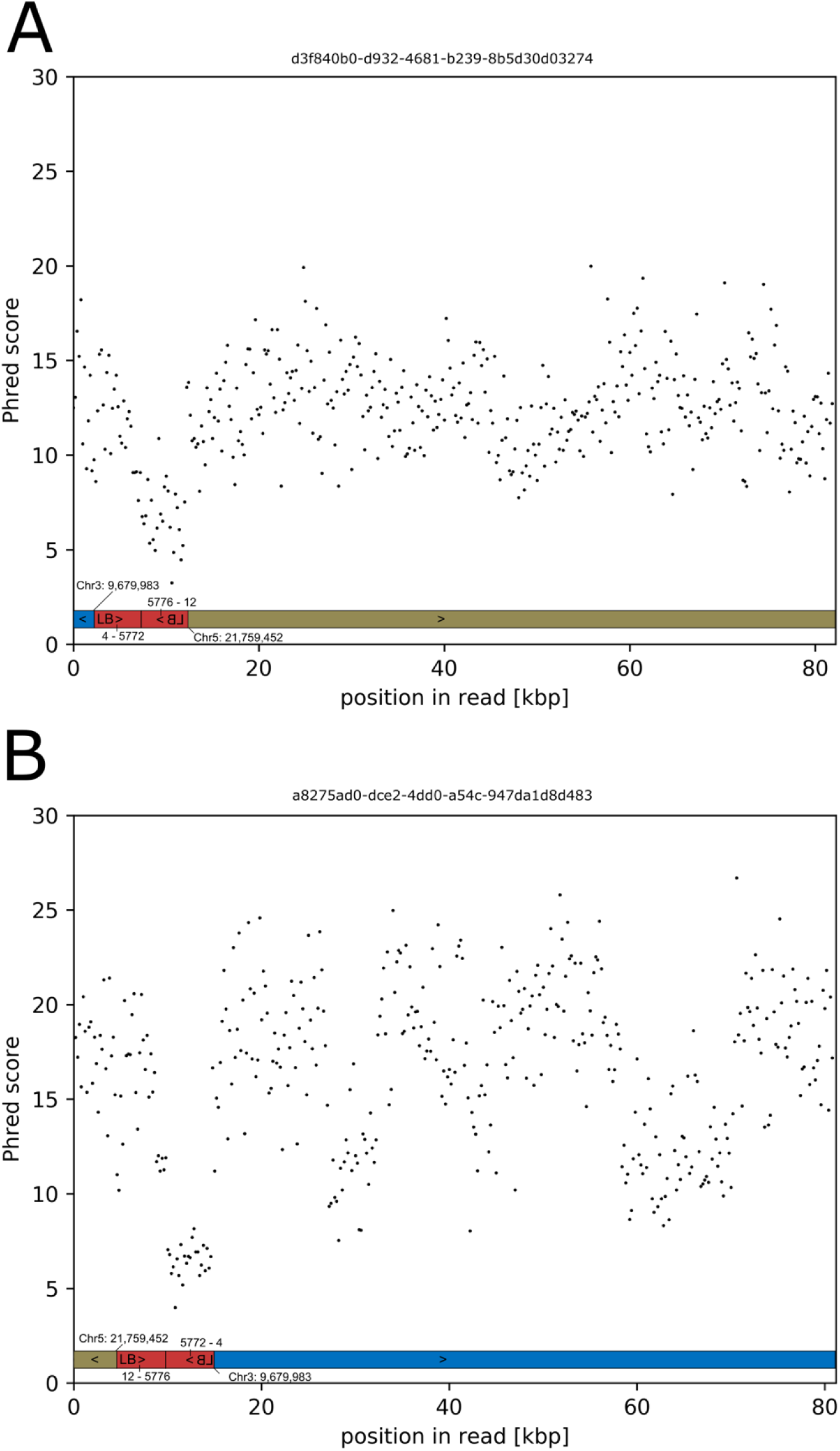
Decrease of Phred score in ONT reads when running from genomic sequence into a T-DNA array. The insertion used as example is from line GK-038B07 and is shown in Fig. 1C. A/B: Read quality data from two ONT reads with opposite direction. The reads are displayed in 5’ to 3’ orientation. Green and blue bars indicate the read parts matching Chr3 and Chr5, respectively. The red bar indicates the position of the T-DNA array which is arranged as inverted repeat. Read IDs of the two examples are shown above each panel. The read quality drop to a Phred score of about 7 occurs when the second part of the inverted repeat is sequenced.

Consequently, the number of miscalled bases in these regions is especially high. These miscalls prevent matches in BLAST searches where a perfect match of several consecutive bases is required as seed for a larger alignment. In some cases, entire reads displayed an extremely low quality. Reads that display such locally increased error rates were found in the context of T-DNA array structures which involve head-to-head or tail-to-tail configurations that have the ability to form foldback structures. It is important to note that the second T-DNA unit was sequenced with lower quality independent of the read direction (Fig. 8).

### Independent Col-0 assembly resolves misassemblies

As mentioned above for the insertion allele 909H04-At1g38212, the detection of rearrangements in the insertion lines is not only dependent on the quality of the reads from or assemblies of the genomes of the lines studied. The results are also dependent on the correctness of the reference sequence. While the quality of the Col-0 reference sequence (the sequence from TAIR9 is still the most recent, see Introduction) is generally of high quality, there are some sequence regions that are not fully resolved. We used a subset of our ONT data, namely ultra-long (> 100 kbp, see Methods) T-DNA free reads, to *de novo* assemble the Col-0 genome sequence. The assembly, designated Col-0_GKat-wt, comprises 35 contigs after polishing and displays an N50 of 14.3 Mbp (GCA_905067165, see Additional file 9). The Col-0_GKat-wt assembly is about 4 Mbp longer than TAIR9 but still does not reach through any of the centromers. Comparison to the TAIR9 sequence indicated that the main gain in assembly length was reached in the pericentromeric regions.

Our collection of ONT sequencing datasets from the GABI-Kat lines provides a combined coverage of over 500x for the TAIR9 reference genome sequence of Col-0. In addition to using ultra-long reads for generating an assembly, the reads were also used for identification of potentially problematic regions in the reference sequence. We identified conflicting regions by evaluating read alignments to assemblies and obtained a list of 383 candidate regions (Additional file 10). We compared selected regions against our *de novo* genome assembly and focused first on the locus At1g38212 (at about 14.3 Mbp of Chr1, see Fig. 7). The differences in this region of the TAIR9 assembly, which were detected when analyzing the T-DNA insertion allele 909H04-At1g38212 (Additional file 6), did show up again. Together with nine other examples selected across all chromosomes, Additional file 11 displays regional comparisons of TAIR9 to Col-0_GKat-wt. The 96 gaps containing various numbers of Ns which are reported for TAIR9 are frequently detected (Additional files 10 and 11).

## Discussion

By sequencing GABI-Kat T-DNA insertion lines with ONT technology, we demonstrate the power of long read sequencing for the characterization of complex T-DNA insertion lines. The complexity of these lines has, at least, four aspects: (i) the number of different insertion loci present in a given line in different regions of the nuclear genome, (ii) the variance of the structures of one or several T-DNA copies appearing at a given insertion locus, (iii) the changes in the genome sequence in the direct vicinity of the T-DNA, and (iv) the changes of chromosome topology related to T-DNA integration.

### Number of T-DNA insertion loci per *A. thaliana* insertion line

The average number of T-DNA insertions per *A. thaliana* T-DNA insertion line is assumed to be about 1.5 [4]. However, the insertion lines available at the stock centers like NASC or ABRC list in almost all cases only one insertion per line. In our limited dataset of 14 lines, 11 new insertions were detected among a total of 26, indicating that one should expect an average of about 2 insertions per line. The 11 new insertions all contain sufficiently intact LB sequences that should have allowed generation of FSTs. The reason for the lack of detection is probably that the FST data generated at GABI-Kat in total have not reached the saturation level, although several insertions are predicted per line at GABI-Kat [23]. The potential of existing T-DNA insertion lines for finding additional knock-out alleles in existing T-DNA insertion lines is also indicated by the fact that TDNA-Seq revealed additional insertion loci in established and FST-indexed lines (see Introduction).

Clearly, analysis by ONT sequencing can effectively reveal additional insertions and can successfully be used to fully characterize the genomes of T-DNA insertion lines. This approach is faster, less laborious, more comprehensive and compared to the level of reliability also significantly cheaper than PCR- or short-read based methods.

### Structure of the inserted T-DNA or T-DNA array

The variance of the T-DNA structures that were resolved by ONT sequencing spans a really wide range of configurations and lengths. Tandem repeats as well as inverted repeats [46] are occurring. Insertion length starts with 2.7 kbp for 909H04-At1g38212 (Fig. 7B) and reaches up to about 50 kbp for 767D12-At2g19210-At2g21385 (Fig. 6C). The lines were selected to contain a T-DNA by checking for resistance to sulfadiazine [22] which is provided by the T-DNA used at GABI-Kat. However, since there are regularly several insertions per line, also T-DNA fragments with a truncated selection marker gene are to be expected - given that resistance is provided in trans. For SALK, SAIL and WISC insertion lines, T-DNA arrays sizes of up to 236 kbp have been reported [19]. We hypothesize that the complexity of T-DNA arrays might correlate with the tendency of selection marker silencing, which could mechanistically be implemented via siRNA [19]. The comparably reduced complexity of T-DNA arrays derived from pAC161 (the binary vectors mostly used at GABI-Kat) could thus explain why the sulfadiazine selection marker stays active for many generations.

Inclusion of BVB sequences in T-DNA array structures has been reported repeatedly for various species [37, 47, 48]. For the studied GABI-Kat lines, BVB sequences were structurally resolved as internal components of T-DNA arrays as well as at the junction to genomic sequences. A total of six T-DNA arrays with BVB sequences were detected among 26 cases, indicating that about 20% of all insertions, and an even higher percentage of lines, contain inserted BVB sequences.

We detected only few intact right border sequences in contrast to left border sequences, which fits the empirical observation that FST-generation for characterization of insertion populations is much more productive at LB than at RB [4–6]. In turn, the lines studied here are selected from insertions detected by using LB for FST generation, which introduces a bias. When insertions accessed via FSTs from RB were studied, RB is found to be more precisely cut than LB [13, 49], which is explained by protection of RB by VirD2 [9]. Nevertheless, within the longer T-DNA arrays and also in the insertions newly detected by ONT sequencing, most of the RBs are lost. This does not fit well to current models for the integration mechanism and explanations for the observed internal “right end to right end” (without RB) fusions in T-DNA arrays and requires further investigation.

### Changes in the genome sequence at the insertion site

Changes in the genome sequence in the direct vicinity of the T-DNA insertion site have already been described in detail [13]. However, that study relied on data from PCR amplicon sequences and could, therefore, not detect events that affect distances longer than the length of an average amplimer of about 2 kbp. In addition, amplicons from both T-DNA::genome junctions were required. Here, we addressed insertions that failed to fulfill the “amplicon sequences from both junctions available” criterion. This allowed to focus on a set of GABI-Kat lines that has a higher chance of showing genomic events (Table 1). The T-DNA::genome junctions studied here fall, with one exception, into the range already described for DSB-based integration and repair by NHEJ, with filler sequences and microhomology at the insertion site [9]. The exception is 909H04-At1g54385-cp-At1g54440 (Fig. 7C), an insertion allele that displays an about 20 kbp long duplicated inversion and in addition 652 bp derived from the plastome at the additional breakpoint that links the inversion back to the chromosome. During repair of the initial DSBs and in parallel to T-DNA integration, also cpDNA is used to join broken ends of DNA at insertion loci. Inversions require more than one repaired DSB in the DNA at the insertion site to be explained, and that cpDNA is available in the nucleus has been demonstrated experimentally [50] and also in the context of horizontal gene transfer [51].

### Genome level changes and translocations related to T-DNA integration

Our analyses revealed five lines with chromosome arm translocations, either exchanged within one chromosome (GK-433E06, Fig. 5) or exchanged between two different chromosomes (Figures 1 to 4). In addition, line GK-767D12 displayed a chromosomal rearrangement that resulted in an inverted duplication of 0.8 Mbp. This aligns well with previous reports of interchromosomal structural variations, translocations, and chromosome fusions in T-DNA insertion lines [18, 19, 33, 34]. Because of the bias for complex cases in the criteria we used for selection of the lines investigated, we cannot deduce a reliable value for the frequency of chromosomal rearrangements in the GABI-Kat population. However, an approximation taking into account that the 6 cases are from 14 lines sequenced, and the 14 lines sequenced are a subset of 342 out of 1,818 lines with attempted confirmation of both borders but failure at the 2^nd^ T-DNA::genome junction, ends up with about one of 10 GABI-Kat lines that may display chromosome-level rearrangements (∼10%). It remains to be determined if this estimation holds true, but the approximation fits somehow to the percentage of T-DNA insertion lines that show Mendelian inheritance of mutant phenotypes (88%) while 12% do not [52]. For the SALK T-DNA population, 19% lines with chromosomal translocations have been reported [18], based on genetic markers and lack of linkage between markers from upstream and downstream of an insertion locus.

Although the number of investigated lines with chromosome arm translocations is small, the high proportion of fusions between Chr3 and Chr5 in our dataset is conspicuous (3 out of 5, see Additional file 3). Also for the line SAIL_232 a fusion of Chr3 and Chr5 was reported [19]. This work addressed 4 T-DNA insertion lines (two SALK, one WISC and SAIL_232) by ONT sequencing and Bionano Genomics (BNG) optical genome maps. Translocations involving chromosomes other than Chr3 and Chr5 were observed in our study and have also been reported before [16–18, 33], but it is possible that translocations between Chr3 and Chr5 occur with a higher rate than others. Full sequence characterization of the genomes of (many) more T-DNA insertion lines by long read sequencing have the potential to reveal hot spots of translocations and chromosome fusions, if these exist.

ONT reads are best suited for the identification of large scale structural variants, for this reason we did not address single nucleotide variants (SNV) or small insertions/deletions (InDels). T-DNA-independent structural variants that were identified, except chromosome fusions, are mostly explained by potential errors in the TAIR9 reference sequence as discussed below. One may conclude that the remaining 8 out of 14 lines for which we have not detected large structural variation are identical in genome organization to the Col-0 wild type. However, this conclusion is based on negative results and should be taken with care.

### Compensating translocations

The chromosome arm translocations detected are all “reciprocal” translocations, which involve two breakpoints and exchange parts of chromosomes. Both rearranged chromosomes are equally detected in the sequenced DNA of the line. The combination of both rearranged chromosomes in the offspring is maintained because homozygosity of only one of the two rearranged chromosomes is expected to be lethal due to imbalance of gene dose for large chromosomal regions. However, if both rearranged chromosomes can be transmitted together in one gametophyte, both rearranged chromosomes might exist in offspring in homozygous state [53]. The induction of chromosome mutations, chromosomal rearrangements and chromosome arm translocations by T-DNA insertion mutagenesis has since a long time received attention in *A. thaliana* genetics. One reason is that these types of mutations cause distorted segregation among offspring since similar segregation results are also indicative of genes essential for gametophyte development [53–55]. With regard to the chromosome arm translocations we detected, it is important to note that some of the arms are fused without integrated T-DNA. The case of line GK-082G09 (one T-DNA insertion locus, still reciprocal chromosome arm translocation) is relevant in this context, because the presence of a single T-DNA insertion per line was used as a criterion to select valid candidates for gametophyte development mutants.

Our analyses of the sequences of T-DNA free chromosomal junctions (e.g. 082G09-At5g57020-0-At3g19080) did not result in the detection of specialties that make these junctions different from T-DNA::genome junctions. We cannot fully exclude that the T-DNA free junctions are the result of recombination of two loci that initially both contained T-DNA, and that one T-DNA got lost during recombination at one of two loci. However, it is also possible that the translocations are the direct result of DSB repair, similar to what has been found after targeted introduction of DSBs [56]. We speculate that both, T-DNA containing and T-DNA free junction cases, result from DSB/integration/repair events that involve genome regions which happen to be in close contact, even if different chromosomes are involved. It is evident that several DSB breaks are required, and repair of these DSB can happen with the DNA that is locally available, might it be cpDNA (see above), T-DNA that must have been delivered to DSB repair sites, or different chromosomes that serve as template for fillers [13] or as target for fusion after a DSB happened to occur.

ONT sequencing of mutants offers easy access to data on presence or absence of translocations and other structural variants. We demonstrated that long read sequencing is a suitable method for characterizing lines that failed in previous characterization attempts based on FSTs. For example, the investigation of T-DNA insertion alleles/lines that display deformed pollen phenotypes, which was impacted by chromosome fusions and uncharacterized T-DNA insertions [55], can now be combined with long read sequencing to reveal all insertion and structural variation events with high resolution. Clearly, comprehensive characterization of T-DNA insertion lines, independent from the population from which the mutant originates, as well as other lines used for forward and reverse genetic experiments, can prevent unnecessary work and questionable results. The entire workflow from DNA extraction to the final genome sequence can be completed in less than a week.

Estimated costs per line for ONT sequencing are $200-300 if multiple samples are consecutively analyzed on one GridION flow cell or if barcoding and multiplexing are applied. The application of “loreta” supports the inspection of T-DNA insertions as soon as the read data are generated. Therefore, we recommend to characterize T-DNA insertion alleles by long read sequencing.

### Analyses of inverted duplicated DNA sequences by ONT sequencing

Decreased quality (Phred scores) was previously described for ONT sequence reads as consequence of inverted repeats which might form secondary structures and thus interfere with the DNA translocation through the nanopore [57]. Complex T-DNA arrays are a challenge to ONT sequencing and probably all other current sequencing technologies because of the frequently included inverted repeat structures. We observed that the second part of an inverted repeat has low sequence quality, while the first part (probably not engaging in secondary structure) is of good sequence quality. Evidence has been provided that changed translocation speed is responsible for the low Phred scores in inverted repeats [57]. The quality decrease needs to be considered especially when performing analyses at the single read level. These stretches of sequence with bad quality also pose a challenge for the assembly, especially since the orientation of the read determines which part of the inverted repeat is of good or poor quality. However, we were able to solve the problem to a satisfying level by manual consideration of reads from the opposite direction which contain the other part of the inverted repeat in good sequence quality.

## Conclusions

This study presents a comprehensive characterization of multiple GABI-Kat lines by long read sequencing. The results argue strongly for full characterization of mutant alleles to avoid misinterpretation and errors in gene function assignments. If an insertion mutant and the T-DNA insertion allele in question are not characterized well at the level of the genotype, the phenotype observed for the mutant might be due to a complex integration locus, and not causally related to the gene that is expected to be knocked-out by the insertion. Structural changes at the genome level, including chromosome translocations and other large rearrangements with junctions without T-DNA, may have confounding effects when studying the genotype to phenotype relations with T-DNA lines. This conclusion must also consider that during the last 20 to 30 years, many T-DNA alleles have been used in reverse genetic experiments. Finally, and similar to the ONT sequence data that resulted from the analyses of four SALK and SAIL/WISC T-DNA insertion lines [19], the ONT sequence data from this study allowed to detect and correct many non-centromeric misassemblies in the current reference sequence.

## Methods

### Plant material

The lines subjected to ONT sequencing were chosen from a collection of GABI-Kat lines which were studied initially to collect statistically meaningful data about the structure of T-DNA insertion sites at both ends of the T-DNA insertions [13]. In this context and also after 2015, confirmation amplicon sequence data from both T-DNA::genome junctions of individual T-DNA insertions were created at GABI-Kat, which was successful for 1,481 cases from 1,476 lines by the end of 2019 (1,319 cases were successfully completed for both junctions in the beginning of 2015). To generate this dataset, 1,835 individual T-DNA insertions from 1,818 lines with one T-DNA::genome junction already confirmed were addressed, meaning that there were 354 cases from 342 lines which failed at the 2^nd^ T-DNA::genome junction. From these 354 cases, we randomly selected the 14 insertions (in 14 different lines) that were studied here, with good germination as additional criterion for effective handling (Additional file 1). Since the focus of interest in insertion alleles was always to identify null alleles of genes, all 14 insertions addressed are CDSi insertions (insertions in the coding sequence or enclosed introns, that means between ATG and Stop codons on genomic DNA). A total of 100 T2 seeds of each line were plated with sulfadiazine selection as described [26]. Surviving T2 plantlets contain at least one integrated T-DNA, either in hemizygous or in homozygous state. Sulfadiazine-resistant plantlets were transferred to soil, grown to about 8-leaf stage and pooled for DNA extraction. For a single locus with normal heritability, statistically 66% of the chromosomes in the pool contain the T-DNA. The T-DNA in GK-654A12 is from pGABI1 [38], the other lines contain T-DNA from pAC161 [22].

### DNA extraction, size enrichment, and quality assessment

Genomic DNA was extracted from young plantlets or young leaves through a CTAB-based protocol (Additional file 12) modified from [22, 58]. We observed, like others before [4], that the quality of extracted DNA decreased with the age of the leaf material processed. Young leaves lead to the best results in our hands. The cause might be increasing cell and vacuole size containing more harmful metabolites which may cause reduced quality and yield of DNA. As DNA quality for ONT sequencing also decreases with storage time, we processed the DNA as soon as possible after extraction. DNA quantity and quality was initially assessed based on NanoDrop (Thermo Scientific) measurement, and on an agarose gel for DNA fragment size distribution. Precise DNA quantification was performed via Qubit (Thermo Fisher) measurement using the broad range buffer following the supplier’s instructions. Up to 9 µg of genomic DNA were subjected to an enrichment of long fragments via Short Read Eliminator kit (Circulomics) according to the suppliers’ instructions.

### Library preparation and ONT sequencing

DNA solutions enriched for long fragments were quantified via Qubit again. One µg DNA (R9.4.1 flow cells) or two µg (R10 flow cells) were subjected to library preparation following the LSK109 protocol provided by ONT. Sequencing was performed on R9.4.1 and R10 flow cells on a GridION. Real time base calling was performed using Guppy v3.0 on the GridION (R9.4.1 flow cells) and on graphic cards in the de.NBI cloud [59] (R10 flow cells), respectively.

### *De novo* genome sequence assemblies

Reads of each GK line were assembled separately to allow validation of other analysis methods (see below). Canu v1.8 [60] was deployed with previously optimized parameters [61]. Assembly quality was assessed based on a previously developed Python script (Table 2). No polishing was performed for assemblies of individual GABI-Kat lines as these assemblies were only used to analyze large structural variants and specifically T-DNA insertions.

**Table 2:** Availability of scripts.

| Description | URLs |
| --- | --- |
| Previously developed scripts for general tasks | https://github.com/bpucker/script_collection |
| Tool for the analysis of ONT datasets ("loreta") | https://github.com/nkleinbo/loreta |
| Scripts for the analyses presented in this study | https://github.com/bpucker/GKseq |

Through removal of all T-DNA reads from the combined ONT read dataset from all insertion lines (Col-0 background [22]) and size filtering, a comprehensive data set of ultra-long reads (longer than 100,000 nt) was generated. This dataset is available from ENA/GenBank with the ID ERS5246674 (SAMEA7490021). The assembly of these ultra-long reads was computed as described above. Polishing was performed with Racon v.1.4.7 [62] and medaka v.0.10.0 as previously described [58]. Potential contamination sequences were removed based on sequence similarity to the genome sequences of other species, and contigs smaller than 100 kbp were discarded as previously described [58, 63]. To ensure accurate representation of the Col-0 wild type genome structure, the assembly was checked for the chromosome fusion events reported for GK-082G09, GK-433E06, and GK-654A12 as well as for the chloroplast DNA integration of GK-909H04. The Sanger reads of the validation amplicons generated for these loci were subjected to a search via BLASTn [64] using default settings. BLASTn was also used to validate the absence of any T-DNA or plasmid sequences in this assembly using pSKI015 (AF187951), pAC161 (AJ537514), and pROK2 [6] as query.

### Analyses of the Col-0_GKat-wt assembly and comparison to TAIR9

To identify differences to the TAIR9 reference genome sequence, the Col-0_GK-wt contigs were sorted and orientated using pseudogenetic markers derived from TAIR9. The TAIR9 sequence was split into 500 bp long sequence chunks which were searched against the Col-0_GK-wt contigs via BLAST. Unique hits with at least 80% of the maximal possible BLAST score were considered as genetic markers. The following analysis with ALLMAPS [65] revealed additional and thus unmatched sequences of Col-0_GK-wt around the centromeres.

ONT reads were aligned to the TAIR9 reference genome sequence via Minimap2 v2.1-r761 [66]. Mappings were converted into BED files with bedtools v2.26 [67]. The alignments were evaluated for the ends of mapping reads, and these ends were quantified in genomic bins of 100 bp using a dedicated tool designated Assembly Error Finder (AEF) v0.12 (Table 2) with default parameters. Neighboring regions with high numbers of alignment ends were grouped if their distance was smaller than 30 kbp. Regions with outstanding high numbers of alignment ends indicate potential errors in the targeted assembly. A selection of these regions from TAIR9 was compared against Col-0_GKat-wt through dot plots [32].

### Analysis of T-DNA insertions

The T-DNA insertions of each line were analyzed in a semi-automatic way. A tool was developed, written in Python and designated “loreta” (Table 2), that needs as input: reads in FASTQ format, T-DNA sequences in FASTA format, a reference file containing sequences for assembly annotation in FASTA format (for this study: sequences of T-DNA and vector backbone, the *A. thaliana* nuclear genome, plastome, chondrome, and the *A. tumefaciens genome*), and – if available – precomputed *de novo* assemblies (as described above). The output are HTML pages with annotated images displaying models of T-DNA insertions and their neighborhood. Partial assemblies of reads containing T-DNA sequences are computed, and the parts of the *de novo* assemblies containing T-DNA sequences are extracted. All resulting sequences as well as all individual reads containing T-DNA sequences are annotated using the reference file. If run on a local machine, a list of tools that need to be installed is given in the github repository (Table 2). For easier access to the tool on a local machine, there is also a Docker file available that can be used to build a Docker image.

Reads containing T-DNA sequences were identified by BLASTn [64, 68] using an identity cutoff of 80% and an e-value cutoff of 1e-50. All identified reads were then assembled using Canu v1.8 with the same parameters as for the *de novo* assemblies (see above) and in addition some parameters to facilitate assemblies with low coverage: correctedErrorRate=0.17, corOutCoverage=200, stopOnLowCoverage=5 and an expected genome size of 10 kbp. From the precomputed *de novo* assemblies, fragments were extracted that contain the T-DNA insertion and 50 kbp up- and downstream sequence. The resulting fragments, contigs from the Canu assembly, the contigs marked as “unassembled” by Canu as well as all individual reads (converted to FASTA using Seqtk-1.3-r106 [69]) were annotated using the reference sequences. For this purpose, a BLASTn search was performed (contig versus reference sequences) with the same parameters as for the identification of T-DNA reads. These BLAST results were mapped to the sequence (contig/read) as follows: BLAST hits were annotated one after another, sorted by decreasing score. If the overlap of a BLAST hit with a previously annotated one exceeds 10 bp, the second BLAST hit was discarded. For further analysis, reads were mapped back to the assembly. All reads were mapped back to the fragments of the *de novo* assembly, reads containing T-DNA were mapped back to (1) the Canu contigs, (2) “unassembled” contigs and (3) individual reads containing T-DNA sequences. Mapping was performed using Minimap2 [66] with the default options for mapping of ONT sequencing data. To further inspect the chromosome(s) sequences prior to T-DNA insertion, the same analysis was performed using the *A. thaliana* sequences neighboring the T-DNA insertion. A FASTA file was generated that contains these flanking sequences using bedtools [67], reads containing this part of *A. thaliana* sequence (and no T-DNA) are again identified using BLAST, assembled and annotated as described above. Infoseq from the EMBOSS package [70] was used to calculate the length of different sequences in the pipeline.

All information is summarized in HTML files containing images; these images display all annotated sequences along with mapped reads and details on the BLAST results. These pages were used for manual inspection and final determination of the insertion structures. For lines with a single canonical insertion, such as 050B11-At5g64610 in GK-050B11 (see Additional file 3), the assembled and annotated contig of the partial assembly was sufficient. In more complex cases like GK-038B07, the exact insertion structure was not clear based on the assembled contigs or based on sections of the *de novo* assembly. If, based on the read mappings shown in the visualization, the partial assembly looked erroneous (many partial mappings), individual reads were used for the determination of the insertion structure. These individual reads were also considered for determination of exact T-DNA positions, if these positions were not clear from contigs / sections. This was often the case for head-to-head or tail-to-tail configurations of T-DNA arrays, where one of the T-DNAs was represented by sequence of low quality and could not be annotated (and led to misassemblies in assembled contigs). These cases were resolved by identification of reads orientated in the other direction, because then the sequence derived from the other T-DNA was of low quality and by combining the annotated results, a clear picture could be derived. If different reads contradicted each other in exact positions of the T-DNA, we chose the “largest possible T-DNA” that could be explained by individual reads.

### Mapping of ONT reads for detection of copy number variation

ONT reads from each line were aligned against the TAIR9 Col-0 reference genome sequence using Minimap v2.10-r761 [71] with the options ‘-ax map-ont --secondary=no’. The resulting mappings were converted into BAM files via samtools v1.8 [72] and used for the construction of coverage files with a previously developed Python script [73]. Coverage plots (see Additional file 5) were constructed as previously described [61] and manually inspected for the identification of copy number variations.

### Sequence read quality assessment

Reads containing T-DNA sequence were annotated based on sequence similarity to other known sequences based on BLASTn [64, 68] usually matching parts of the Ti-plasmid or *A. thaliana* genome sequence. Reads associated with complex T-DNA insertions were considered for downstream analysis if substantial parts (>1 kbp) of the read sequence were not matched to any database sequences via BLASTn. Per base quality (Phred score) of such reads was assessed based on a sliding window of 200 nt with a step size of 100 nt.

### Chromosome fusion and cpDNA insertion validation via PCR and Sanger sequencing

Chromosomal fusions without a connecting T-DNA were analyzed via PCR using manually designed flanking primers (Additional file 4). Amplicons were generated using genomic DNA extracted from plants of the respective line as template with Q5 High-Fidelity DNA polymerase (NEB) following supplier’s instructions. PCR products were separated on a 1% agarose gel and visualized using ethidiumbromide and UV light. Amplicons were purified with Exo-CIP Rapid PCR Cleanup Kit (NEB) following supplier’s instructions. Sanger sequencing was performed at the Sequencing Core Facility of the Center for Biotechnology (Bielefeld University, Bielefeld, Germany) using BigDye terminator v3.1 chemistry (Thermo Fisher) on a 3730XL sequencer. The resulting Sanger sequences were merged using tools from the EMBOSS package [70]. After transferring reverse reads to their reverse complement using revseq, a multiple alignment was generated using MAFFT [74]. One consensus sequence for each amplicon was extracted from the alignments using em_cons with option -plurality 1, and the resulting sequences were submitted to ENA (see Additional file 1 for accession numbers).

### Analyses of T-DNA free chromosome fusion junctions

The five junction sequences were analyzed by BLAST essentially as described [13]. Briefly, searches were performed against all possible target sequences (*A. tumefaciens*; *A. thaliana* nucleome, plastome and chondrome; T-DNA and vector backbone) using BLASTn default parameters. If the complete query was not covered, the unmatched part of the query sequence was classified as filler and extracted. Subsequently, this sequence was used in a BLAST search with an e-value cutoff of 10, a word-size of 5 and the ‘-task “blastn-short” option activated to detect smaller and lower quality hits. If this was not successful (as in GK-909H04), the filler sequence was extended by 10 bases up- and downstream and the procedure described above was repeated.

## Declarations

### Ethics approval and consent to participate

Not applicable

### Consent for publication

Not applicable

### Availability of data and materials

Sequence read datasets generated and analyzed during this study were made available at ENA under the accession PRJEB35658. Individual run IDs are included in Additional file 1. The Col-0 genome sequence assembly of the GABI-Kat Col-0 genetic background (Col-0_GKat-wt) is available at ENA under the accession GCA_905067165.

### Authors’ contribution

BP performed DNA extraction and sequencing. BP and NK performed bioinformatic analyses. BP, NK, and BW interpreted the results and wrote the manuscript.

## Supplementary information

**Additional file 1**: Summary of GABI-Kat insertion line data, segregation data for the F2 families after selection for sulfadiazine resistance, sequencing data including run IDs from submission to ENA/SRA, and accession numbers of T-DNA free junction sequences.

**Additional file 2**: Extended version of Table 1 covering all 14 lines.

**Additional file 3**: Overview of the T-DNA insertions and associated structural variants in the investigated GABI-Kat lines.

**Additional file 4**: Sequences of oligonucleotides used for the validation of fusion points of chromosomal translocations and other large structural variants.

**Additional file 5**: Read coverage of all analyzed lines in relation to the TAIR9 reference genome sequence.

**Additional file 6**: Structure of genomic locus around one insertion in GK-909H04.

**Additional file 7**: Analysis results of genomic fusion junction sequences without T-DNA insertion.

**Additional file 8**: Visual overview over all insertions detected.

**Additional file 9**: Assembly statistics of Col-0_GK-wt.

**Additional file 10**: Potential errors in the TAIR9 reference genome sequence of Col-0.

**Additional file 11**: Dot plots between TAIR9 and Col-0_GK-wt for potential errors in the reference sequence.

**Additional file 12**: Protocol for the extraction of genomic DNA from *A. thaliana* for ONT sequencing.

## Acknowledgements

We are very grateful to the Bioinformatics Resource Facility support team of the CeBiTec and to de.NBI for providing computing infrastructure and excellent technical support. We also thank the Sequencing Core Facility of the CeBiTec for granting access to the ONT infrastructure. Many thanks to Tobias Busche, Christian Rückert, and Jörn Kalinowski for general support related to the ONT sequencing, and to Prisva Viehöver for high quality Sanger sequencing. We thank Andrea Voigt for excellent technical support. The bioinformatic work was supported in part by grants from the German Federal Ministry of Education and Research (BMBF) for the project “Bielefeld-Gießen Center for Microbial Bioinformatics–BiGi” (grant no. 031A533) within the German Network for Bioinformatics Infrastructure (de.NBI).

