## Additional file 2 for "Large scale genomic rearrangements in selected *Arabidopsis thaliana* T-DNA lines are caused by T-DNA insertion mutagenesis"

Additional file 2 (full Table 1): Key findings summary of ONT-sequenced GABI-Kat T-DNA insertion lines.

| **Line ID^a^** | **number of insertions** | | | | **Summary of observation** |
| --- | --- | --- | --- | --- | --- |
|  | **FST pred.** | **PCR- conf.** | **total found** | **ONT found** |  |
| **GK-038B07** | 2 | 0 (1)^b^ | 4^c^ | 3 | FST-predicted insertion in Chr5 is part of a fusion of Chr3 and Chr5, 2 T-DNA arrays detected at translocation fusion points of which one contains in addition an inversion of ~2 Mbp fused with another T-DNA array, additional insertion of a complex T-DNA array in Chr1. |
| GK-040A12 | 3 | 1 (2)^b^ | 1 | 0 | Insertion of a T-DNA array consisting of 3 almost complete copies of the T-DNA, both T-DNA::genome junctions at LB. |
| GK-050B11 | 1 | 1 | 1 | 0 | One shortened canonical T-DNA inserted with intact LB, the other end contains neither RB nor LB regions which explains failure to confirm the 2^nd^ T-DNA::genome junction. |
| **GK-082G09** | 2 | 1 | 1^c^ | 0 | Fusion of Chr3 and Chr5, both FSTs were derived from one fused locus containing a complex array of T-DNA copies that includes part of the binary vector backbone, the compensating fusion of Chr5 and Chr3 does not contain a T-DNA at the translocation fusion point, failure to confirm the 2^nd^ T-DNA::genome junction is explained by the fact that is was expected on the same chromosome. |
| **GK-089D12** | 1 | 1 | 2 | 1 | Fusion of Chr3 and Chr5, FSTs are derived from the single DNA::genome junction that contains LB, both translocation fusion points contain mostly canonical T-DNAs, failure to confirm the 2^nd^ T-DNA::genome junction explained by shortened T-DNA. |
| GK-290G05 | 1 | 1 | 1 | 0 | One shortened canonical T-DNA inserted with intact LB detected by an FST, the other end contains neither RB nor LB regions which explains failure to confirm the 2^nd^ DNA::genome junction. |
| GK-399C06 | 2 | 1 | 2 | 0 | Two canonical T-DNA insertions that match the loci predicted from FSTs, the PCR-confirmed insertion in Chr2 contains a deletion of about 800 bp at insertion site and partially repetitive sequences that explain failure to confirm 2^nd^ DNA::genome junction. |
| GK-410B07 | 1 | 1 | 1 | 0 | One complex T-DNA array insertion consisting of 4 T-DNA copies, both T-DNA::genome junctions at LB, deletion of about 1.8 kbp at insertion site which explains failure to confirm 2^nd^ T-DNA::genome junction, our automatic primer design did not consider the second FST from the same locus that would have allowed to correctly predict the 2^nd^ DNA::genome junction. |
| GK-430F05 | 1 | 1 | 3 | 2 | Three T-DNA insertion sites in total, two complex T-DNA arrays with one of the two containing more than 5 kbp binary vector backbone, the confirmed T-DNA insertion in Chr3 is also complex, no theoretical explanation for failure of the confirmation PCR at the 2^nd^ DNA::genome junction, one insertion in the pericentromeric region of (probably) Chr4 that contains a rearranged RB region. |
| **GK-433E06** | 4 | 1 | 1^c^ | 0 | The terminal parts of Chr1 with telomers are translocated in an intra-chromosomal exchange, only the southern translocation fusion point which connects the northern end of Chr1 to the southern arm contains a complex T-DNA array, the northern translocation fusion point does not contain T-DNA, translocation explains failure to confirm 2^nd^ T-DNA::genome junction. |
| **GK-654A12**^d^ | 1 | 1 | 2^c^ | 1 | Fusion of Chr1 and Chr4 with a complex T-DNA array that also contains binary vector backbone, Chr1-part of the fusion predicted by FST, the compensating fusion of Chr4 and Chr1 does not contain T-DNA at the translocation fusion point, translocation explains failure to confirm 2^nd^ T-DNA::genome junction, additional insertion of a T-DNA array in Chr2 with a 162 bp duplicated inversion at the integration site. |
| **GK-767D12** | 1 | 1 | 2 | 1 | Large segmental duplication and inversion at the predicted insertion site, a very long T-DNA array also containing binary vector backbone present at the southern end of the segmental duplication, inversion explains failure to confirm 2^nd^ T-DNA::genome junction, the northern fusion point of the inverted segmental duplication does not contain T-DNA, another canonical T-DNA insertion in Chr2 but with 4 Mbp distance. |
| **GK-909H04** | 1 | 1 | 3 | 2 | The predicted T-DNA insertion contains an inverted duplication of about 20 kbp at the integration site which explains why the 2^nd^ T-DNA::genome junction could not be confirmed by PCR, integration of 652 bp derived from the plastome at the northern fusion point of the duplication but no T-DNA, two additional canonical T-DNA insertions in Chr1 and Chr2. |
| GK-947B06 | 1 | 1 | 2 | 1 | Complex T-DNA array insertion of 3 T-DNAs at the predicted insertion site in Chr1, no theoretical explanation for failure of the confirmation PCR at the 2^nd^ DNA::genome junction, additional complex insertion containing 2 T-DNAs and binary vector backbone on Chr2. |

^a^ Lines discussed in detail in the text are marked in **bold**. These are the lines with chromosomal rearrangements and GK-909H04. Please refer to Additional file 3 for more detail on individual insertions.

^b^ The T-DNA insertion that was used for line selection turned out to be a false positive case that was not detectable in the ONT read data of this line.

^c^ One locus in a T-DNA line containing a translocation due to chromosome fusion causes the FSTs from this single locus to map to two places in the reference sequence. These two places might also be the source of potentially existing FST from the compensating chromosome fusion. If the compensating chromosome fusion does not contain a T-DNA, the number of insertions is lower than expected.

^d^ The T-DNA in 654A12 is derived from pGABI1 (Ulker et al. 2008, Nature Biotechnology 26:1015-1017) while the other lines contain the T-DNA from pAC161 (Rosso et al. 2003, Plant Molecular Biology 53:247-259).
