## Additional file 5 for "Large scale genomic rearrangements in selected *Arabidopsis thaliana* T-DNA lines are caused by T-DNA insertion mutagenesis"

Additional file 5: Read coverage of all analyzed lines in relation to the TAIR9 reference genome sequence.

This analysis was performed as previously described (Pucker et al. (2019), Genes 10:671). The coverage values were log2 transformed to accommodate different values in the same figure. Outliers around the centromeres and other not completely assembled regions are due to read mapping artifacts and not due to *bona fide* variations.

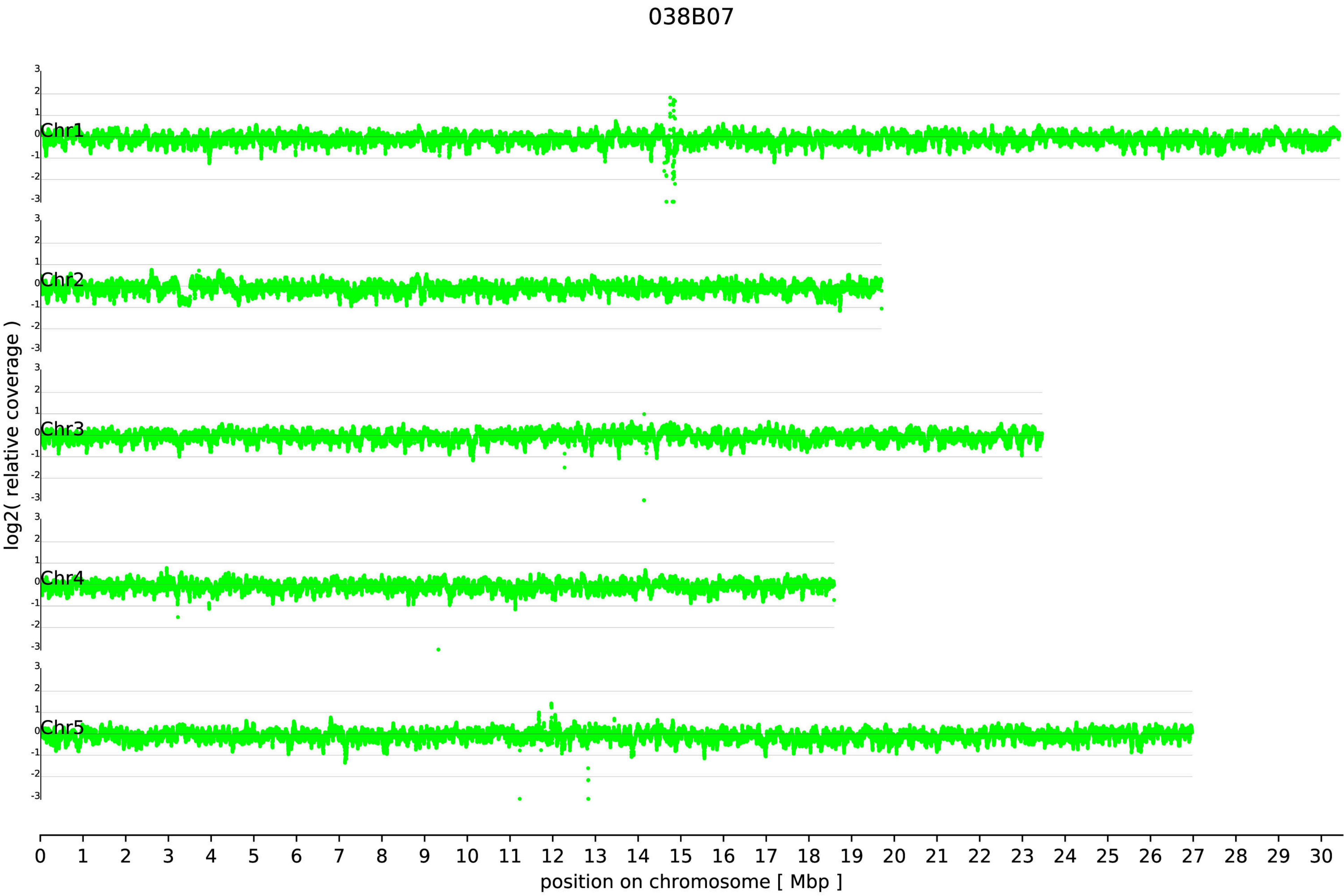

038B07
Chr1
Chr2
Chr3
Chr4
Chr5
log2( relative coverage )
position on chromosome [ Mbp ]

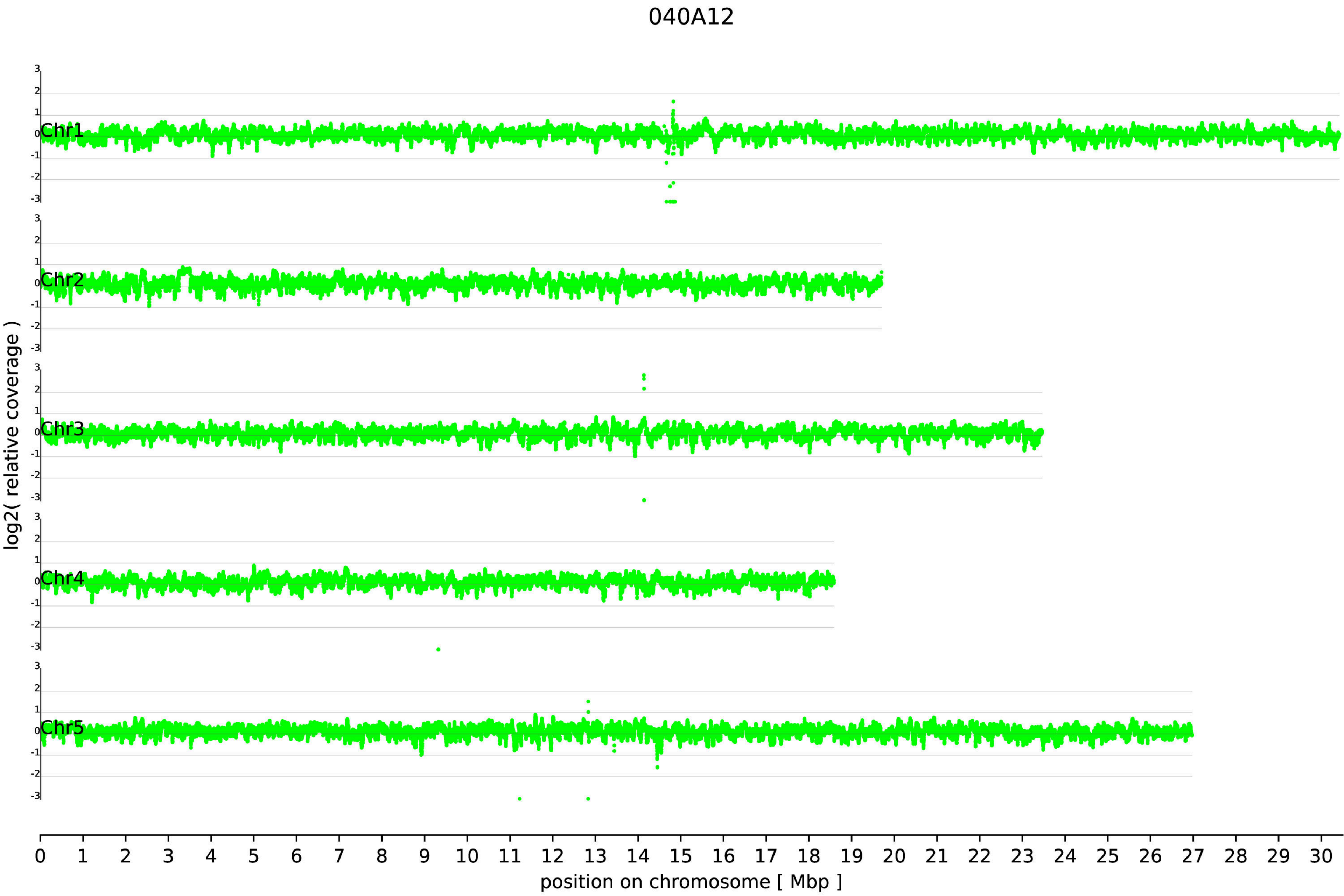
040A12
log2( relative coverage )
Chr1
Chr2
Chr3
Chr4
Chr5
position on chromosome [ Mbp ]

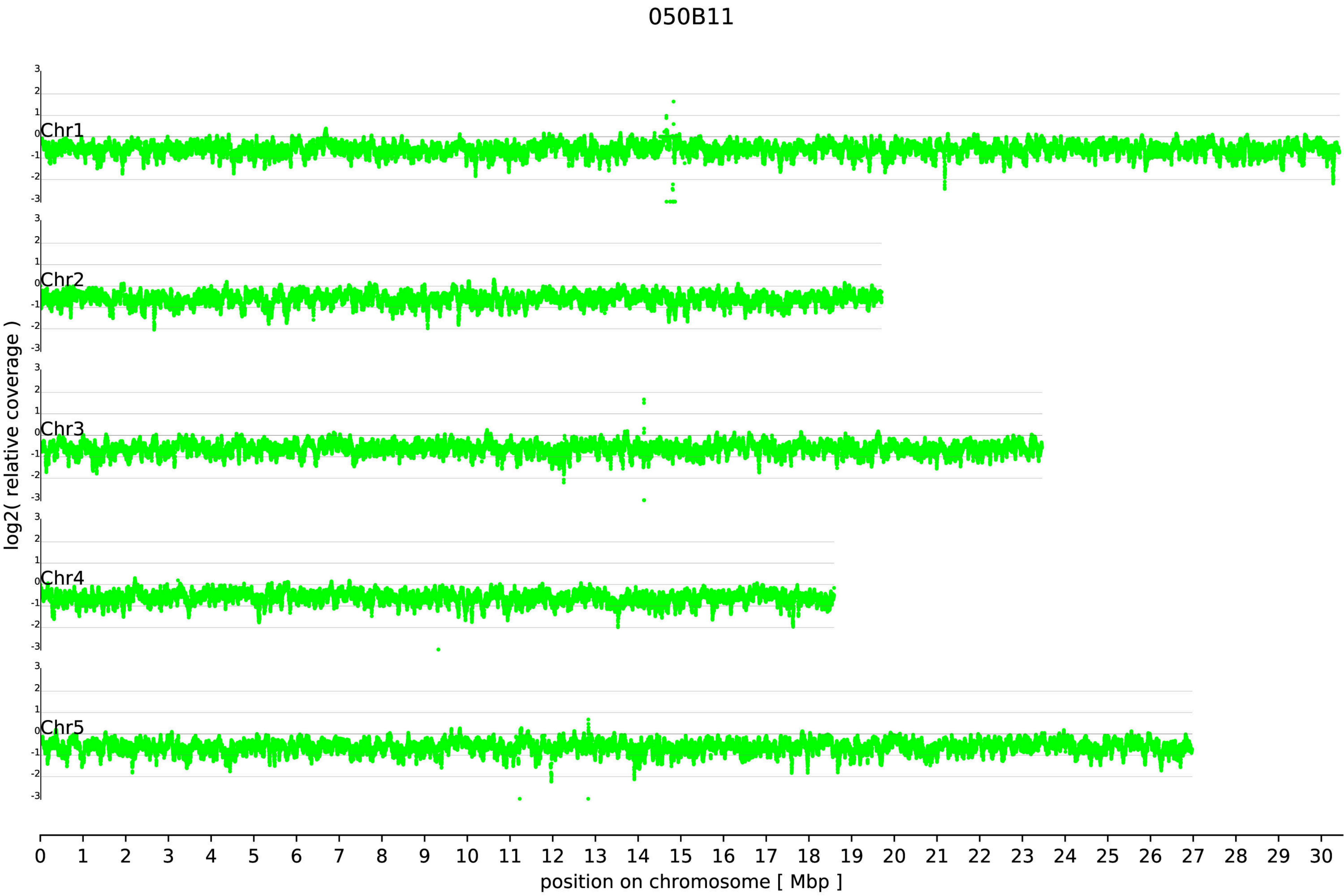

050B11
Chr1
Chr2
Chr3
Chr4
Chr5
3
2
1
0
-1
-2
-3
log2( relative coverage )
0 1 2 3 4 5 6 7 8 9 10 11 12 13 14 15 16 17 18 19 20 21 22 23 24 25 26 27 28 29 30
position on chromosome [ Mbp ]

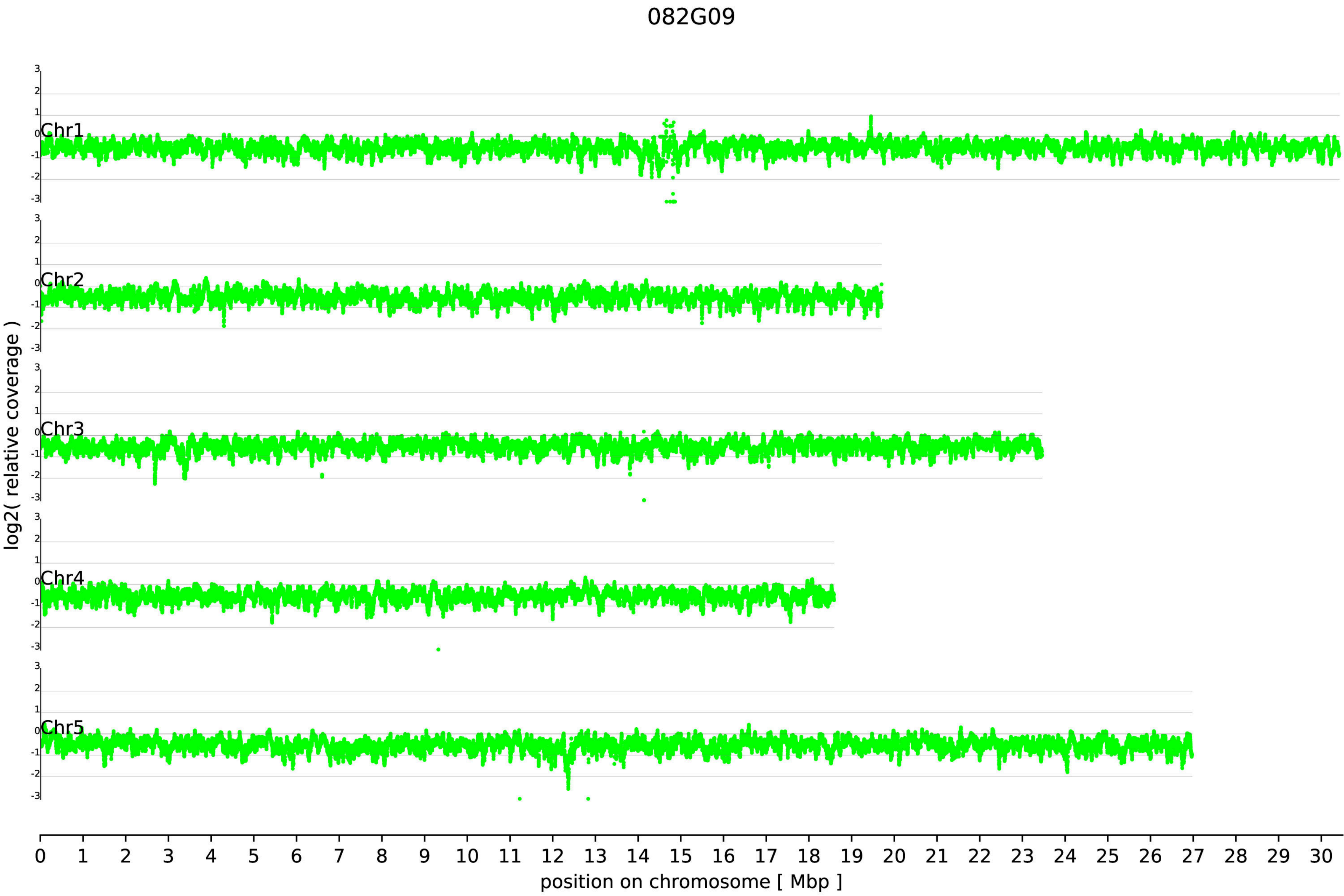
082G09
log2( relative coverage )
Chr1
Chr2
Chr3
Chr4
Chr5
3
2
1
0
-1
-2
-3
0 1 2 3 4 5 6 7 8 9 10 11 12 13 14 15 16 17 18 19 20 21 22 23 24 25 26 27 28 29 30
position on chromosome [ Mbp ]

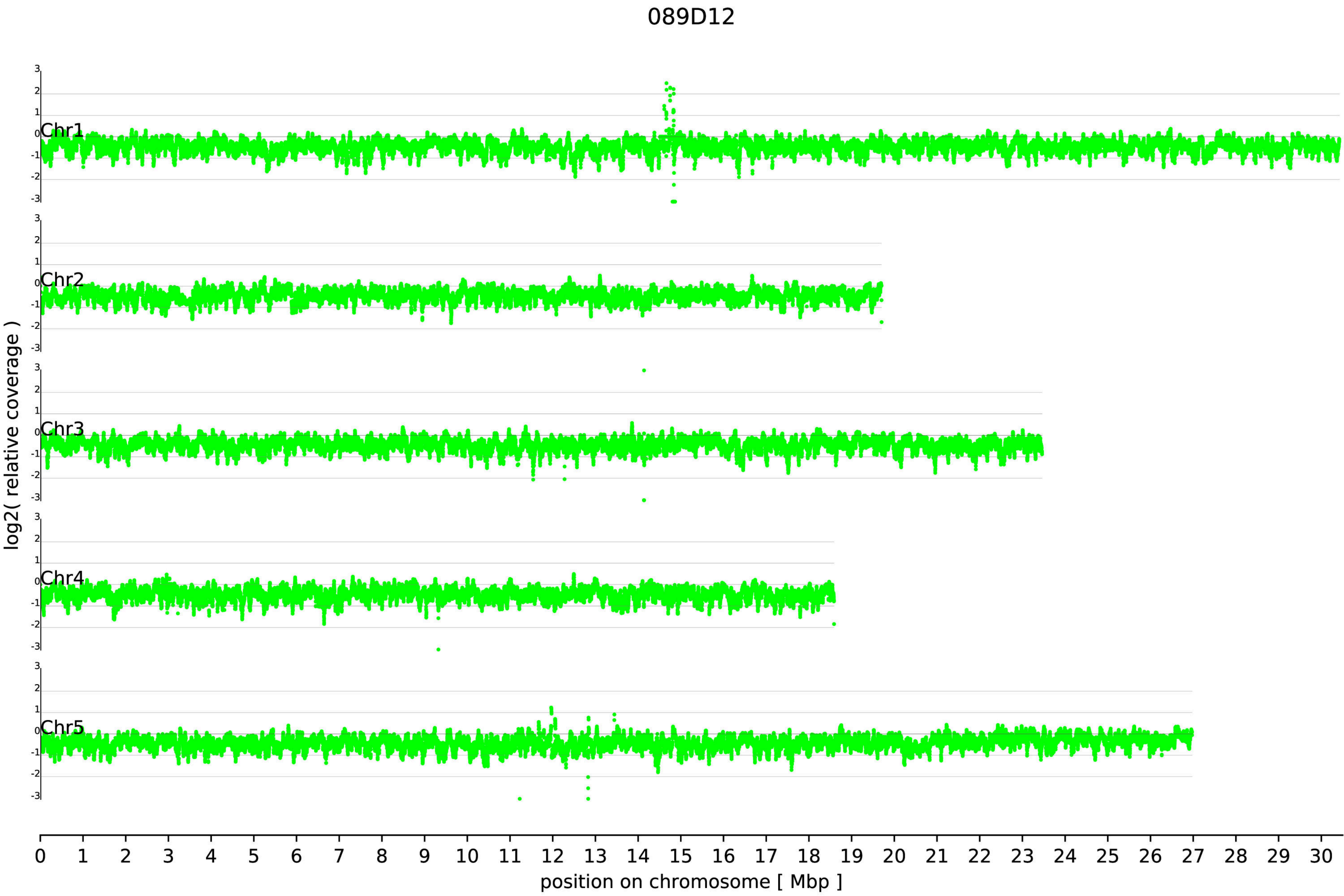

089D12
log2( relative coverage )
Chr1
Chr2
Chr3
Chr4
Chr5
3
2
1
0
-1
-2
-3
position on chromosome [ Mbp ]

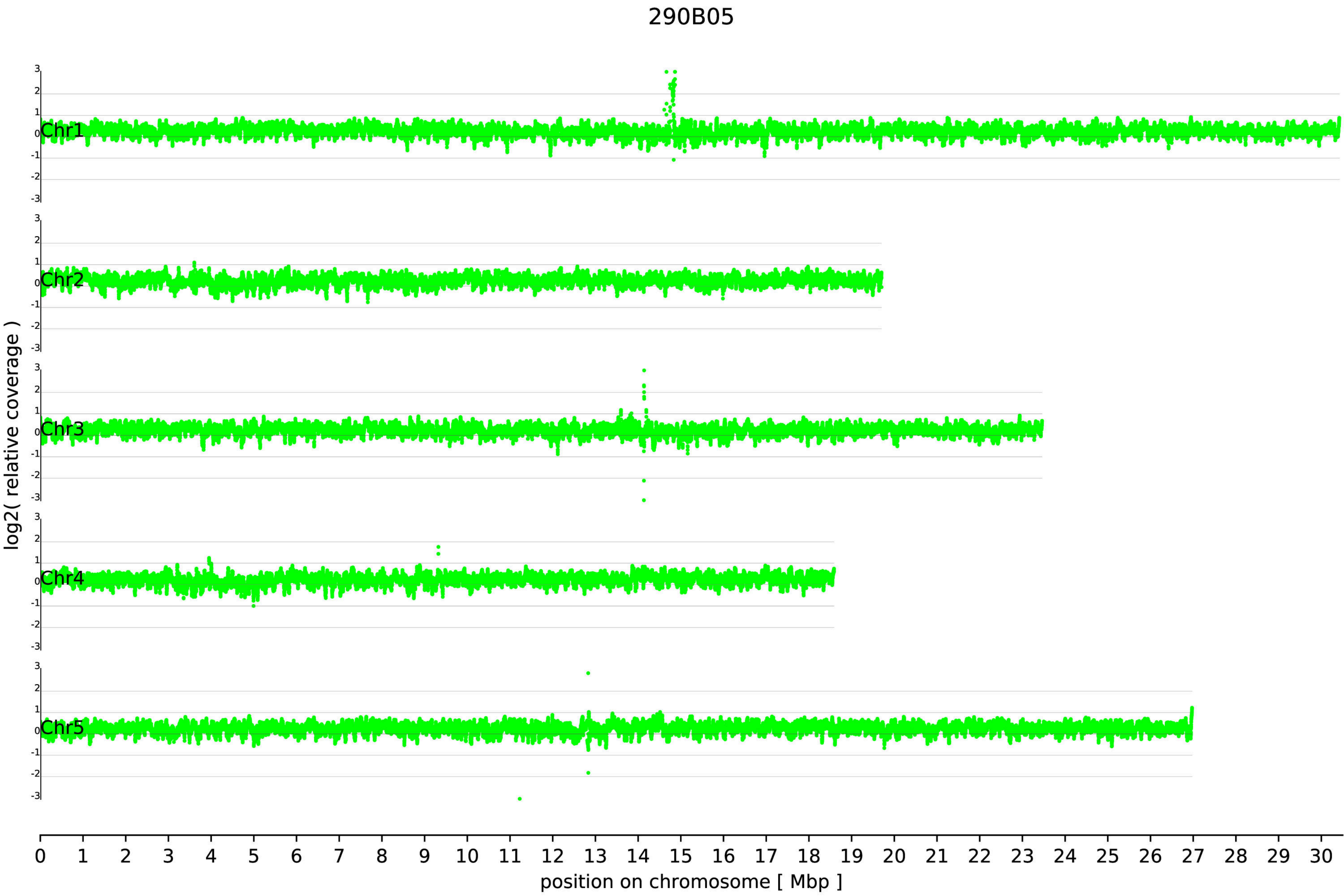
290B05
log2( relative coverage )
Chr1
Chr2
Chr3
Chr4
Chr5
3
2
1
0
-1
-2
-3
0 1 2 3 4 5 6 7 8 9 10 11 12 13 14 15 16 17 18 19 20 21 22 23 24 25 26 27 28 29 30
position on chromosome [ Mbp ]

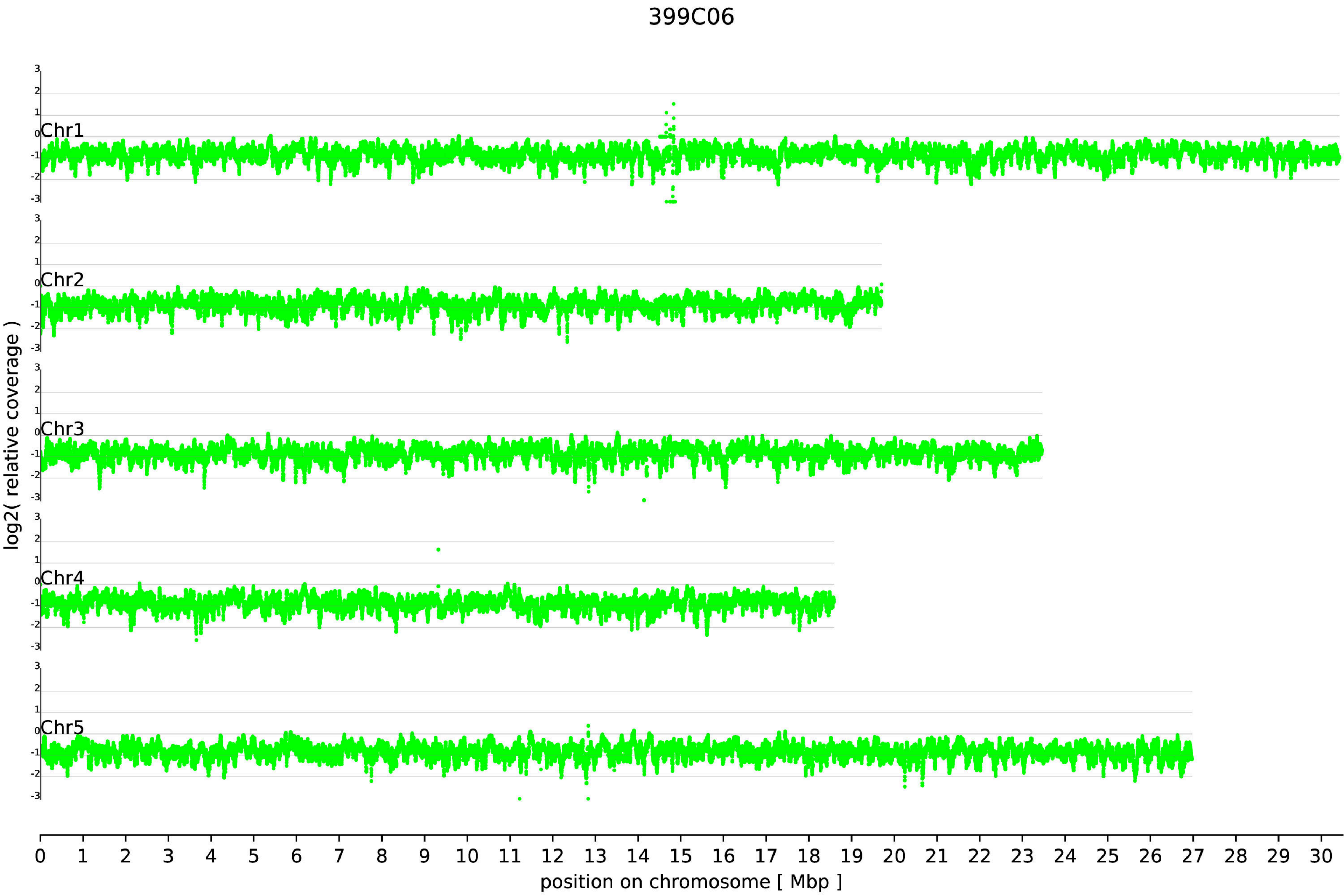

399C06
Chr1
Chr2
Chr3
Chr4
Chr5
log2( relative coverage )
position on chromosome [ Mbp ]

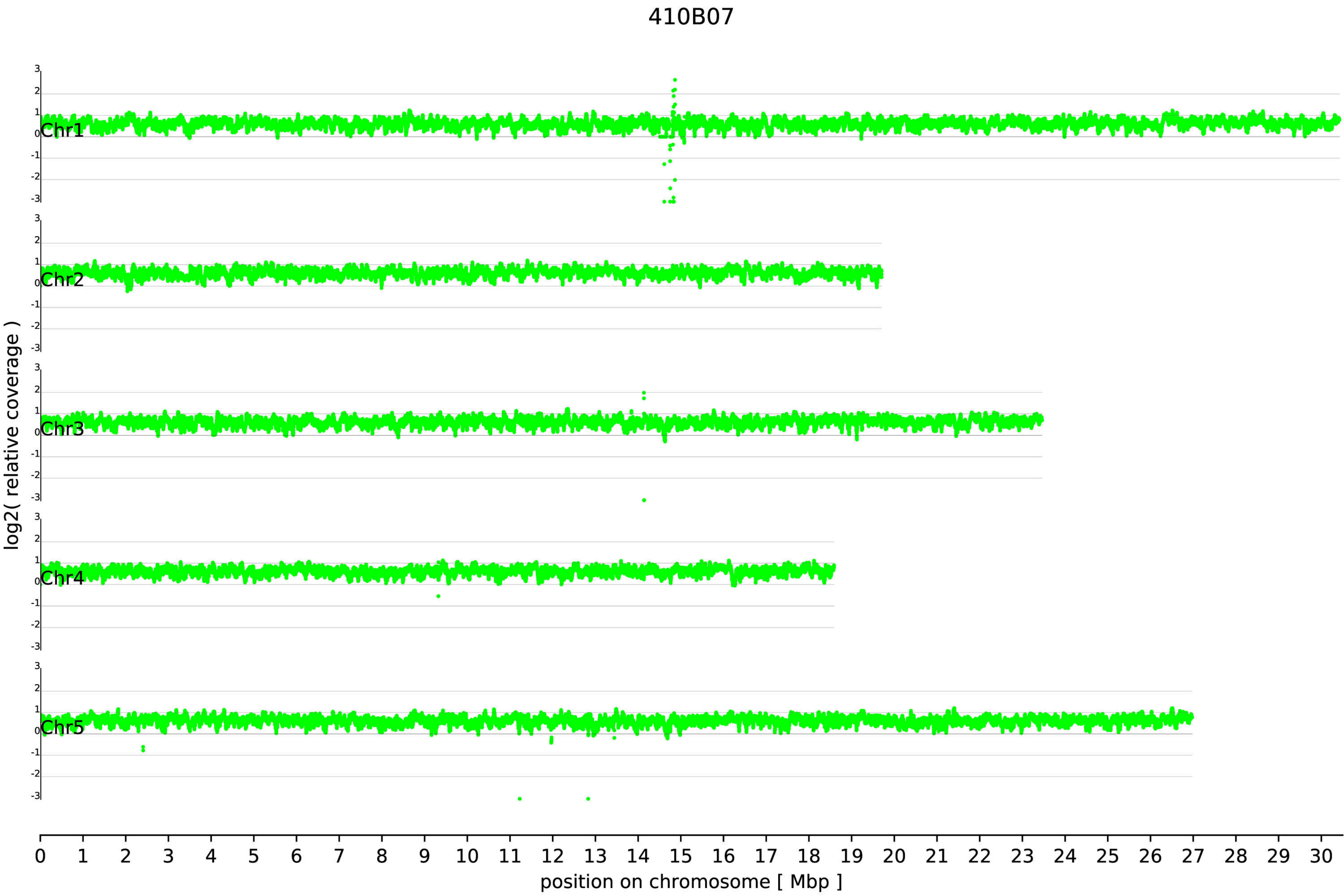

410B07
Chr1
Chr2
Chr3
Chr4
Chr5
log2( relative coverage )
position on chromosome [ Mbp ]

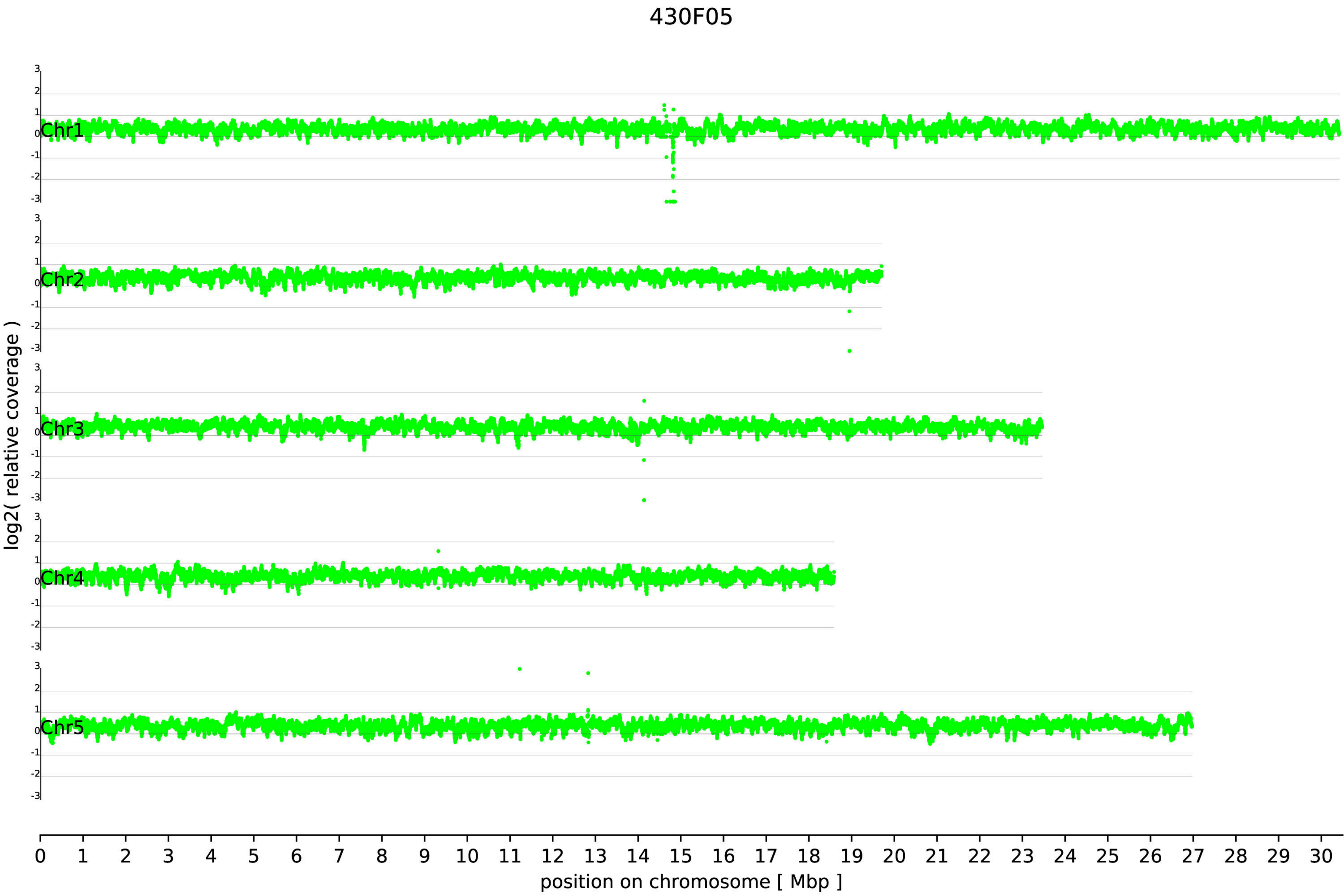
430F05
Chr1
Chr2
Chr3
Chr4
Chr5
log2( relative coverage )
3
2
1
0
-1
-2
-3
0 1 2 3 4 5 6 7 8 9 10 11 12 13 14 15 16 17 18 19 20 21 22 23 24 25 26 27 28 29 30
position on chromosome [ Mbp ]

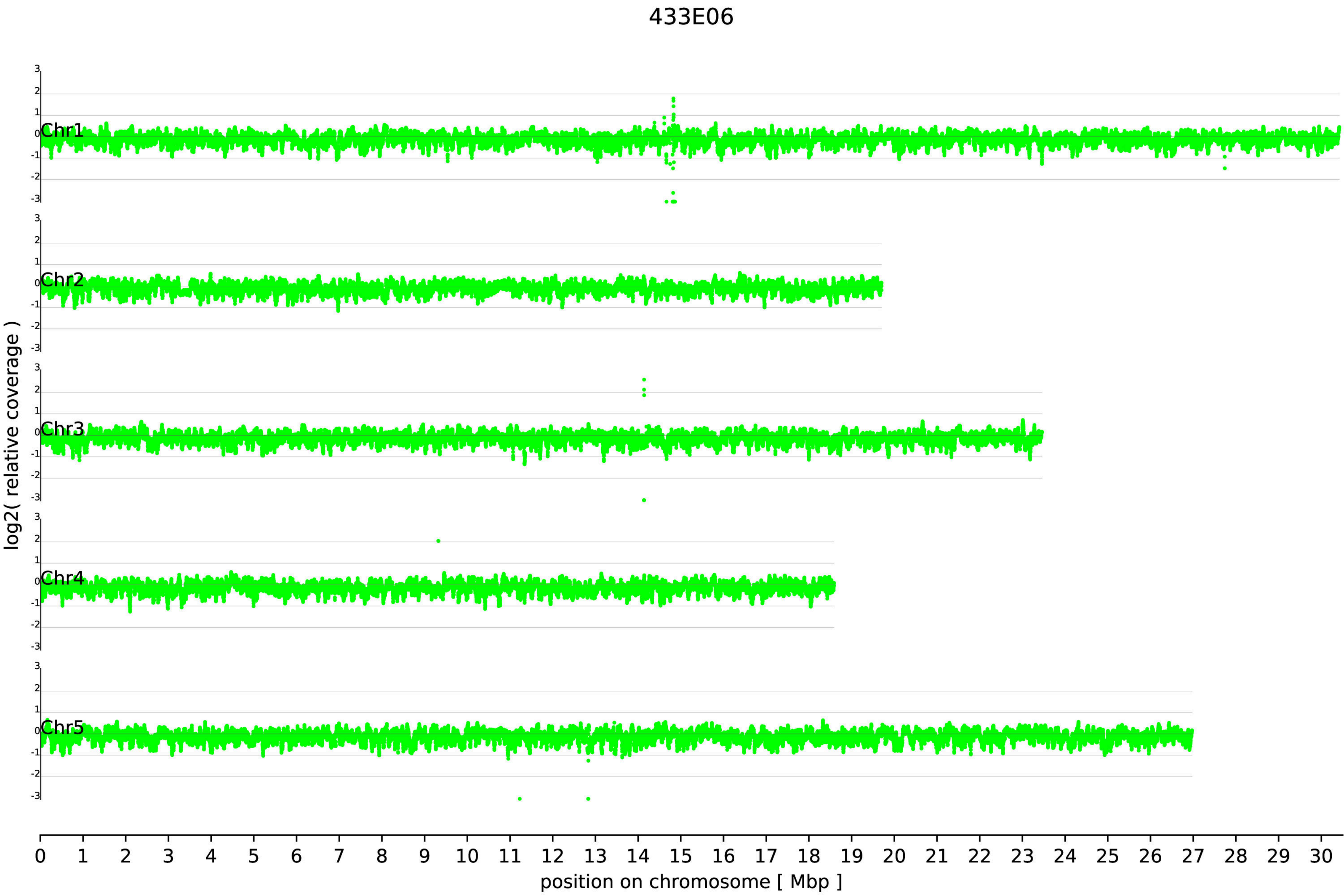
433E06
Chr1
Chr2
Chr3
Chr4
Chr5
log2( relative coverage )
position on chromosome [ Mbp ]

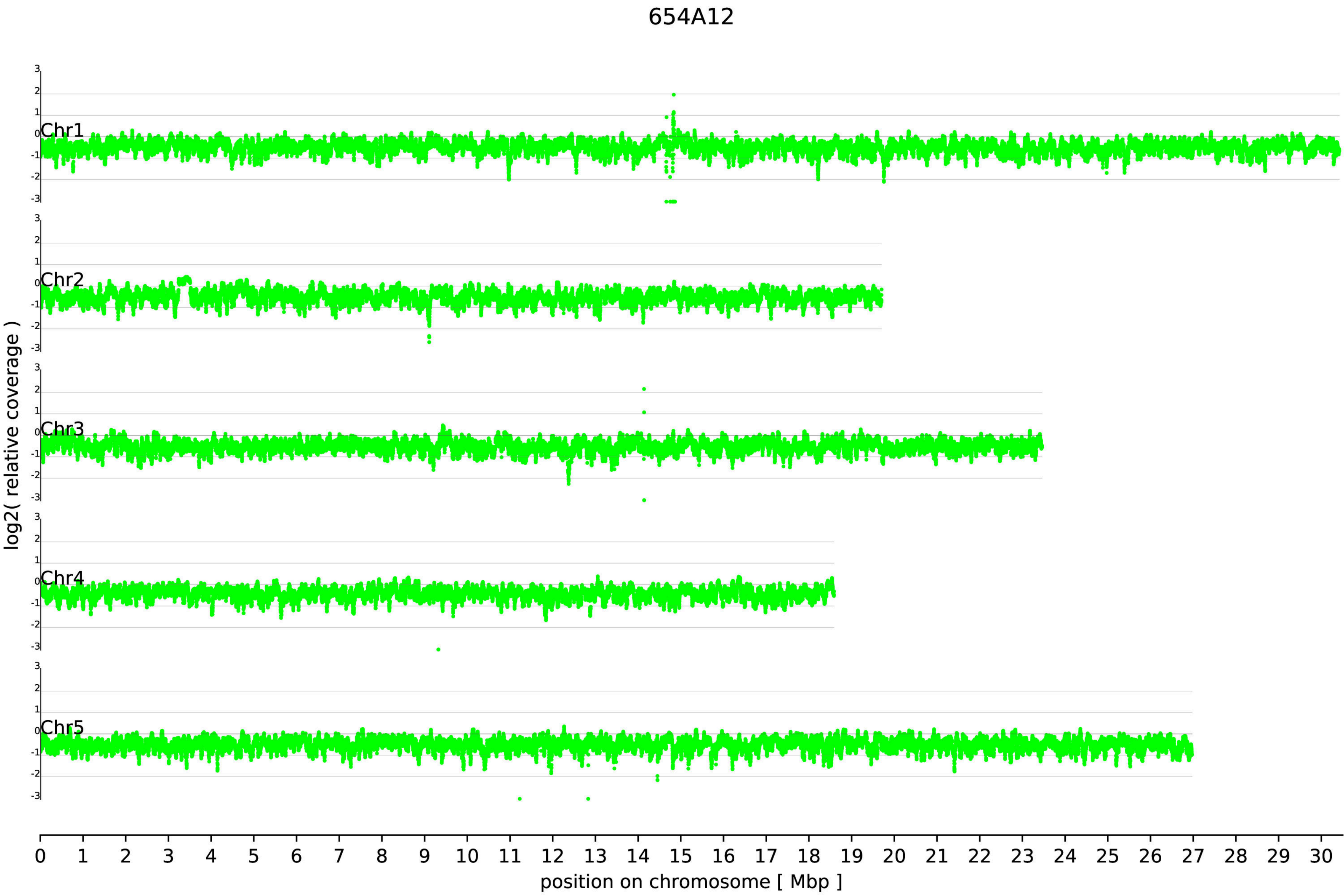
654A12
Chr1
Chr2
Chr3
Chr4
Chr5
log2( relative coverage )
position on chromosome [ Mbp ]

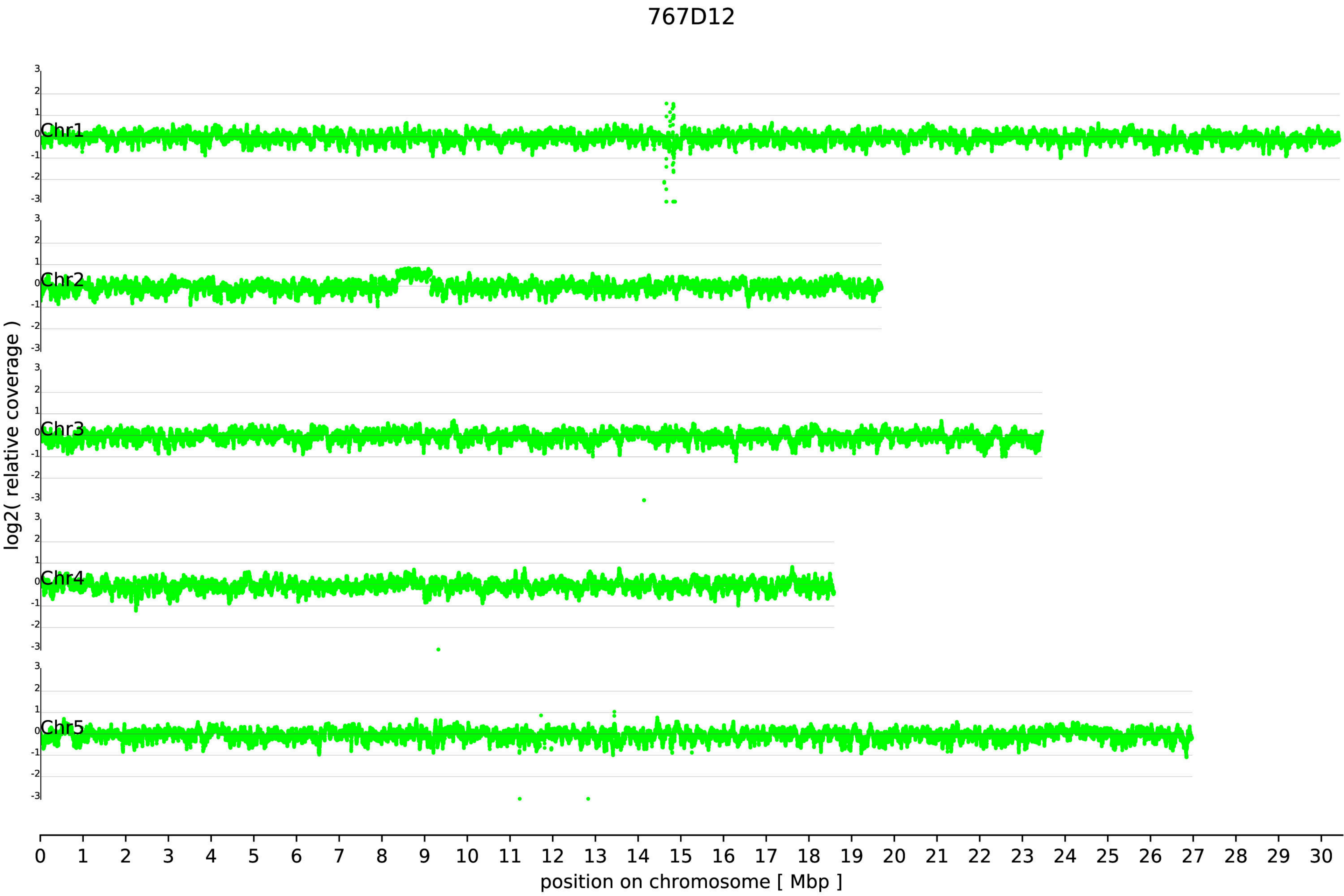

767D12
Chr1
Chr2
Chr3
Chr4
Chr5
3
2
1
0
-1
-2
-3
log2( relative coverage )
0 1 2 3 4 5 6 7 8 9 10 11 12 13 14 15 16 17 18 19 20 21 22 23 24 25 26 27 28 29 30
position on chromosome [ Mbp ]

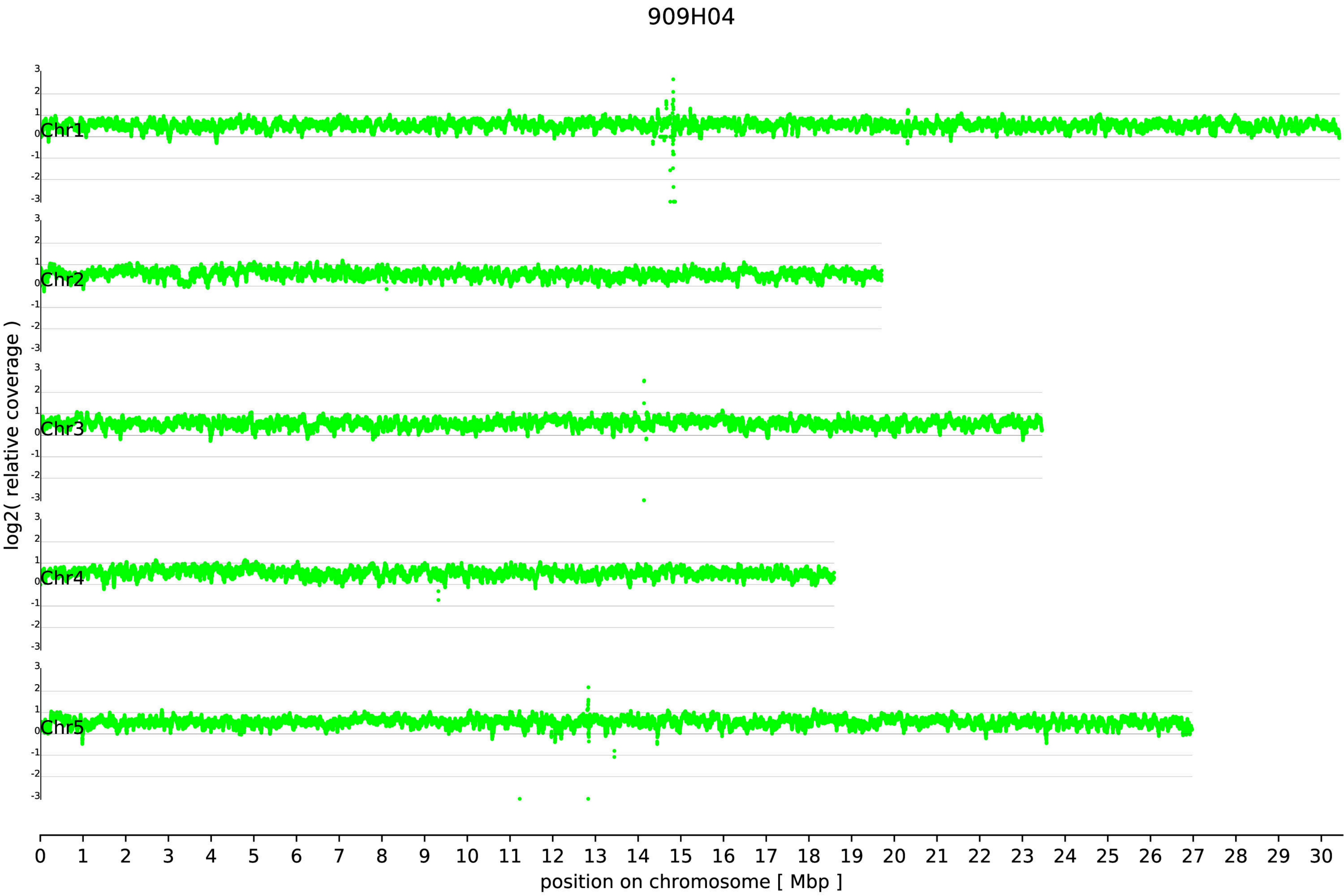

909H04
log2( relative coverage )
Chr1
Chr2
Chr3
Chr4
Chr5
3
2
1
0
-1
-2
-3
position on chromosome [ Mbp ]

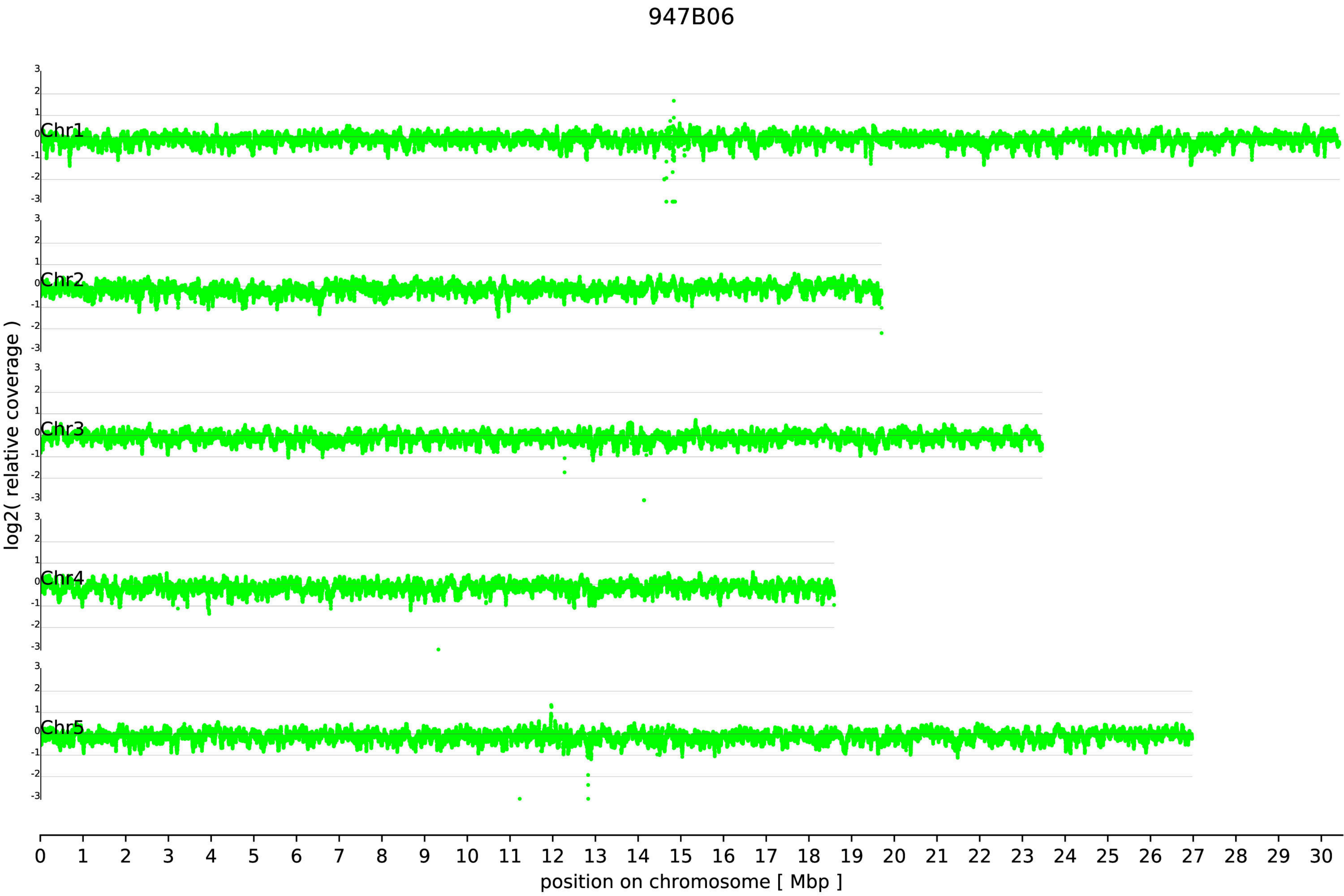
947B06
Chr1
Chr2
Chr3
Chr4
Chr5
3
2
1
0
-1
-2
-3
log2( relative coverage )
0 1 2 3 4 5 6 7 8 9 10 11 12 13 14 15 16 17 18 19 20 21 22 23 24 25 26 27 28 29 30
position on chromosome [ Mbp ]
