## Additional file 6 for "Large scale genomic rearrangements in selected *Arabidopsis thaliana* T-DNA lines are caused by T-DNA insertion mutagenesis"

Additional file 6: Structure of genomic locus around one insertion in GK-909H04.

Black lines indicate the TAIR9 and Col-0_GKat-wt genome sequences, red block represents the T-DNA, orange arrows indicate the positions of mapped ONT reads, blue blocks indicate the position of repeats and contain their length in bp.

### locus 909H04-At1g38212

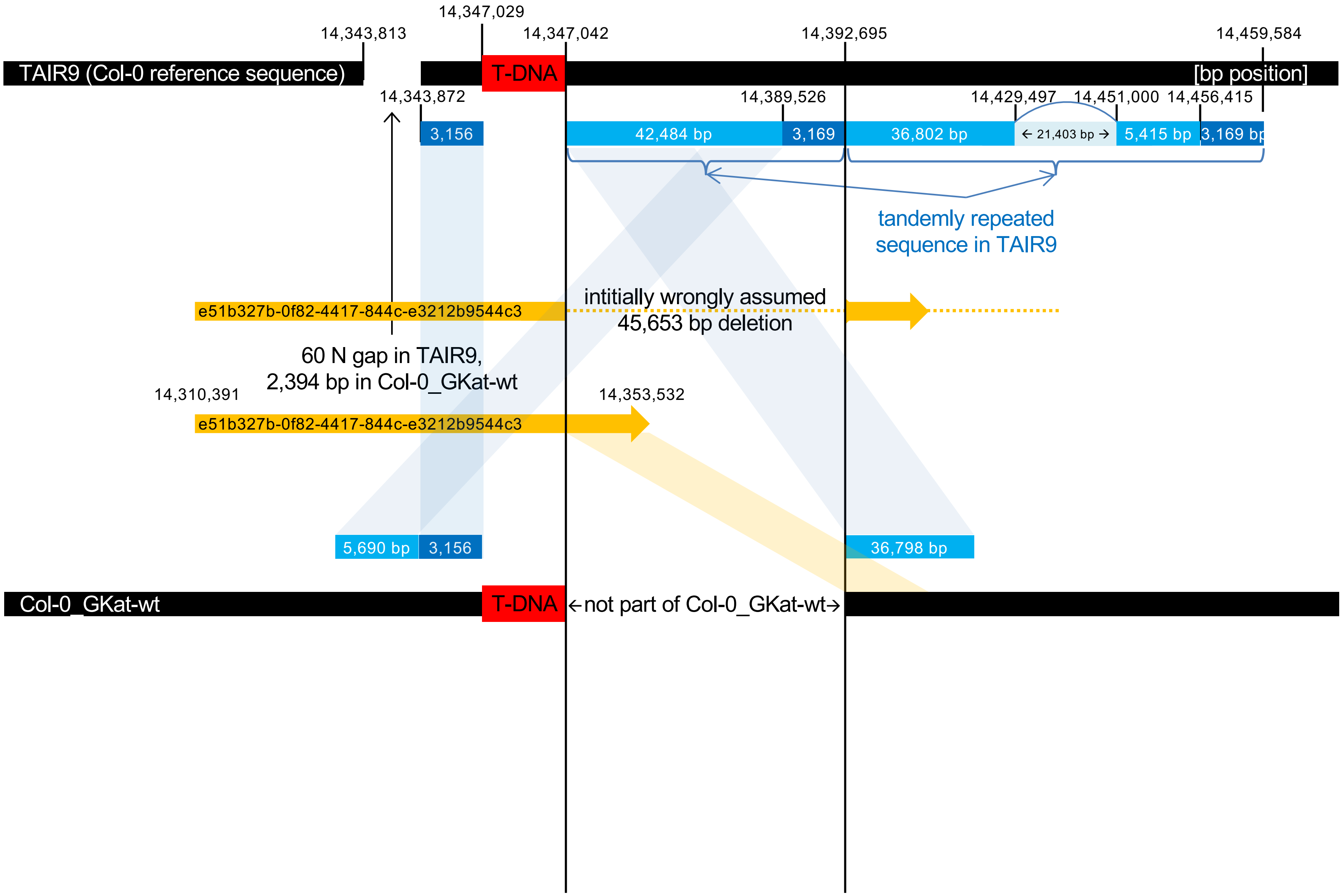
