## Additional file 7 for "Large scale genomic rearrangements in selected *Arabidopsis thaliana* T-DNA lines are caused by T-DNA insertion mutagenesis"

**Additional file 7: Analysis results of genomic fusion junction sequences without T-DNA insertion.**

**GK-082G09**

The junction sequence from 082G09-At5g57020-0-At3g19080 showed the following features:

1-269: Chr5:23077183<-23076915

270-311: Filler

312-538: Chr3:6602070->6602295

The filler has the following sequence:

>GK-082G09:270-311

ACATGTGAATATTCAATTAGTACCAGTTTACATGTGATATCG

… and with additional 10 bp up- and downstream:

>GK-082G09:260-321

TCAACTCTCTACATGTGAATATTCAATTAGTACCAGTTTACATGTGATATCGTTTTATAGGT

Origin of the filler:

1-21:Chr1:28384085->28384104

24-38:Chr1:15512555->15512569

The filler seems to consist of two shorter partial sequences. However, the exact origin of the filler is not clear, since there were several almost identical BLAST hits at different locations in the genome sequence.

**GK-433E06**

The junction sequence from 433E06-F9L1-0-At1g73770 showed the following features:

1-154: Chr1:27744148<-27743995

147-322: Chr1:5237655->5237830

The junction thus shows a microhomology of 8 bp.

**GK-654A12**

The junction sequence from 654A12-FCAALL-0-At1g45688 showed the following features:

1-228: Chr4:8469597->8469824

229-243: Filler

244-517: Chr1:17192289->17192562

The filler has the following sequence:

>GK-654A12:229-243

TGTATGTAGATTTTT

… and with additional 10 bp up- and downstream:

>GK-654A12:219-253

ATCTTTTATTTGTATGTAGATTTTTCTTTGAGAAT

BLAST search of the filler revealed the following origin:

1-15: Chr4:893885<-893871

However, the exact origin of the filler is not clear, since there were several almost identical BLAST hits at different locations in the genome sequence.

**GK-767D12**

The junction sequence from 767D12-At2g21385-0-At2g19210 showed the following features:

1-555: Chr2:8353340<-8352784

555-966: Chr4:8338361->8338772

The junction thus shows a microhomology of 1 bp.

**GK-909H04**

The two junction sequences from 909H04-At1g54385-cp-At1g54440 showed the following features:

1-182: Chr1:20303800->20303979

183: 1bp Filler

184-212: Chr1:20306530<-20306502

213-220: Filler #1

221-261: Chr1:20307057->20307097

304-949: Cp:133454<-134105

950-957: Filler #2

958-1183: Chr1:20326823<-20326598

Filler #1 has the following sequence:

>GK-909H04:213-220

caagcccc

… and with additional 10 bp up- and downstream:

>GK-909H04:203-230

aatgcgtaaacaagccccacaatacatg

Filler #1 had no BLAST hits, but with 10 bp up- and downstream revealed the following origin:

11-26: Chr1:8213672<-8213657

Filler #2 has the following sequence:

>GK-909H04:950-957

tggaaaaa

… and with additional 10 bp up- and downstream:

>GK-909H04:940-967

gctaagttcatggaaaaattaggttcta

Filler #2 had no BLAST hits, but with 10 bp up- and downstream revealed the following origin:

8-27: Chr5:11909972->11909991

However, for both fillers, there are similar good BLAST hits at different locations in the genome sequence.
