## Additional file 8 for "Large scale genomic rearrangements in selected *Arabidopsis thaliana* T-DNA lines are caused by T-DNA insertion mutagenesis"

Additional file 8: Visual overview over all insertions detected.

Color codes in the ideograms were used for the five chromosomes; N, northern end of chromosome; S, southern end of chromosome. T-DNA insertions are indicated in red. Local assemblies of the T-DNA insertion loci as well as chromosomal fusions are displayed to reveal their detailed structure. Read coverage plots are used to show deletion an/or duplication of genomic regions. For additional details about the figure elements see legend to figures 1 and 6 in the main text.

# GK-038B07

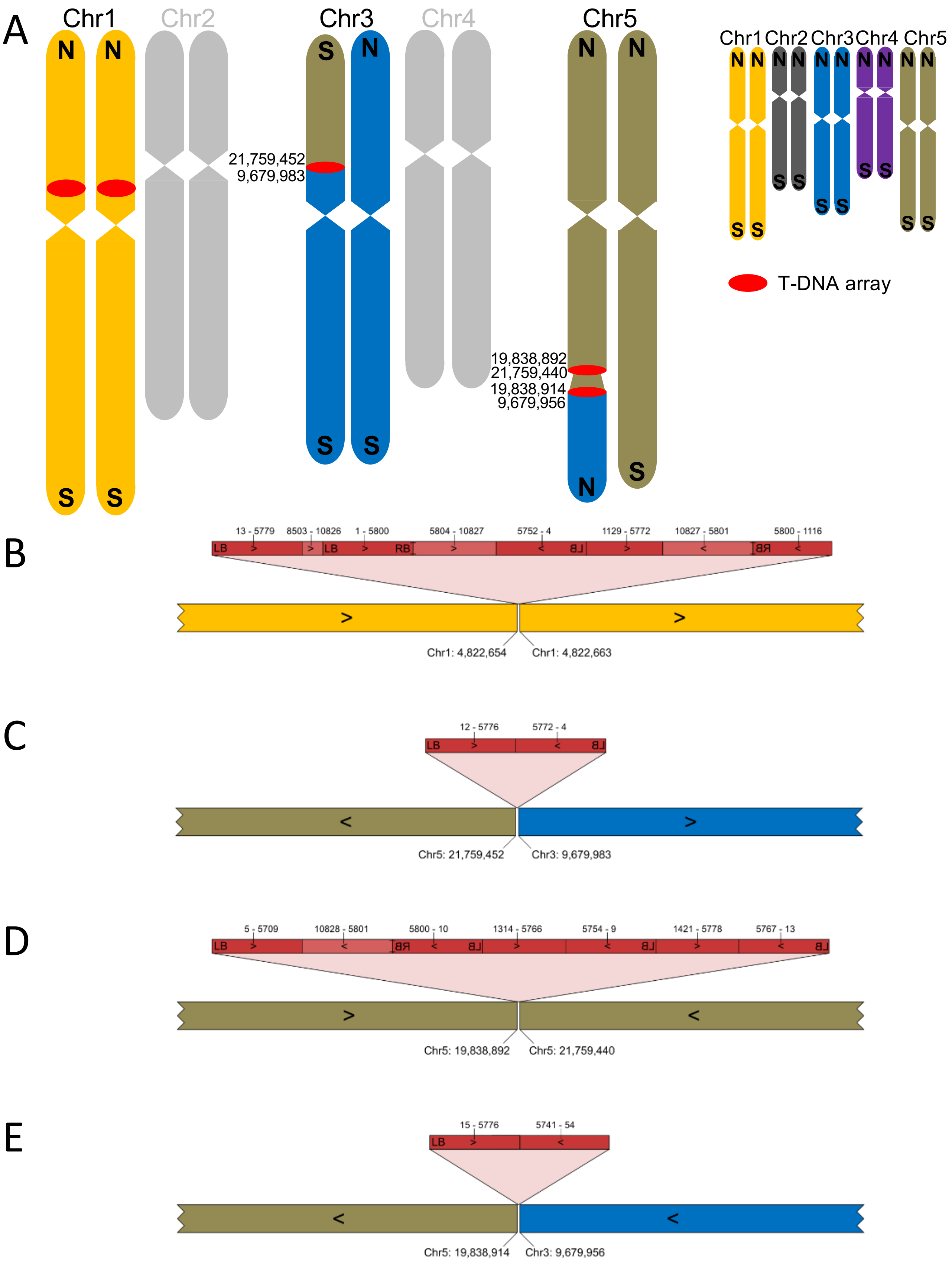
A
Chr1
Chr2
Chr3
Chr4
Chr5
21,759,452
9,679,983
19,838,892
21,759,440
19,838,914
9,679,956
T-DNA array
B
13 - 5779
8503 - 10826
1 - 5800
5804 - 10827
5752 - 4
1129 - 5772
10827 - 5801
5800 - 1116
Chr1: 4,822,654
Chr1: 4,822,663
C
12 - 5776
5772 - 4
Chr5: 21,759,452
Chr3: 9,679,983
D
5 - 5709
10828 - 5801
5800 - 10
1314 - 5766
5754 - 9
1421 - 5778
5767 - 13
Chr5: 19,838,892
Chr5: 21,759,440
E
15 - 5776
5741 - 54
Chr5: 19,838,914
Chr3: 9,679,956

# GK-040A12

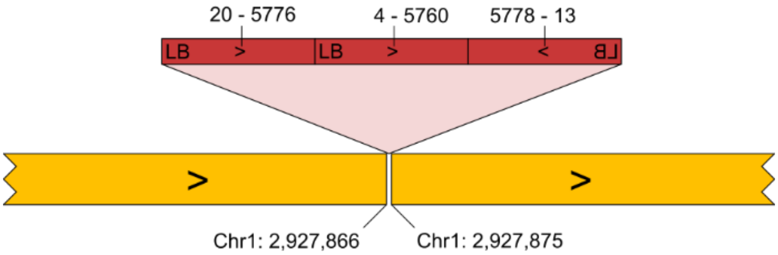

# GK-050B11

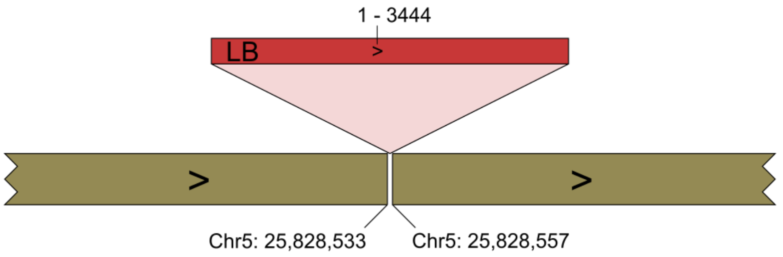

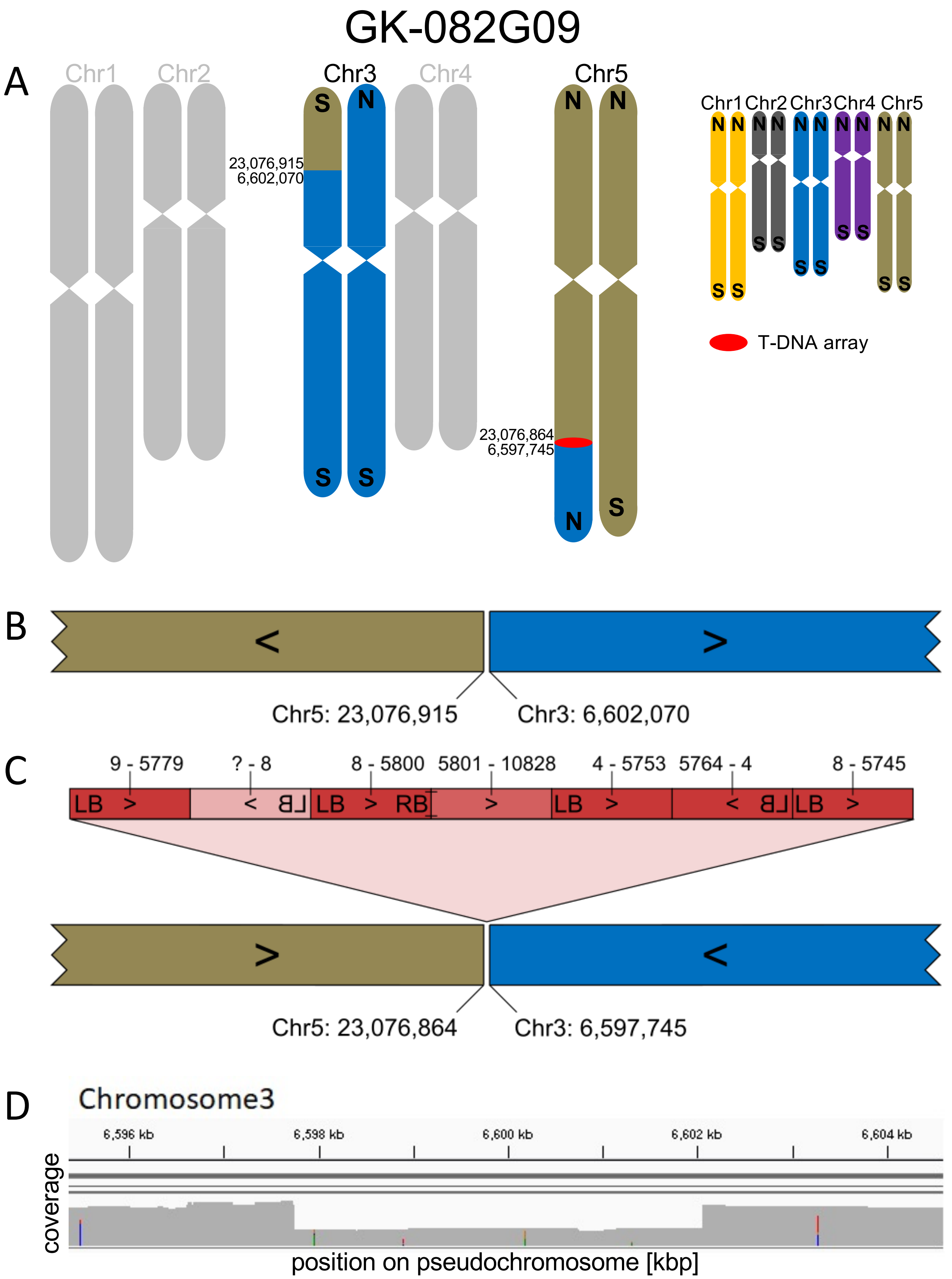

GK-082G09
A
Chr1
Chr2
Chr3
Chr4
Chr5
S
N
23,076,915
6,602,070
23,076,864
6,597,745
T-DNA array
B
Chr5: 23,076,915
Chr3: 6,602,070
C
9 - 5779
? - 8
8 - 5800
5801 - 10828
4 - 5753
5764 - 4
8 - 5745
LB
RB
Chr5: 23,076,864
Chr3: 6,597,745
D
Chromosome3
6,596 kb
6,598 kb
6,600 kb
6,602 kb
6,604 kb
coverage
position on pseudochromosome [kbp]

# GK-089D12

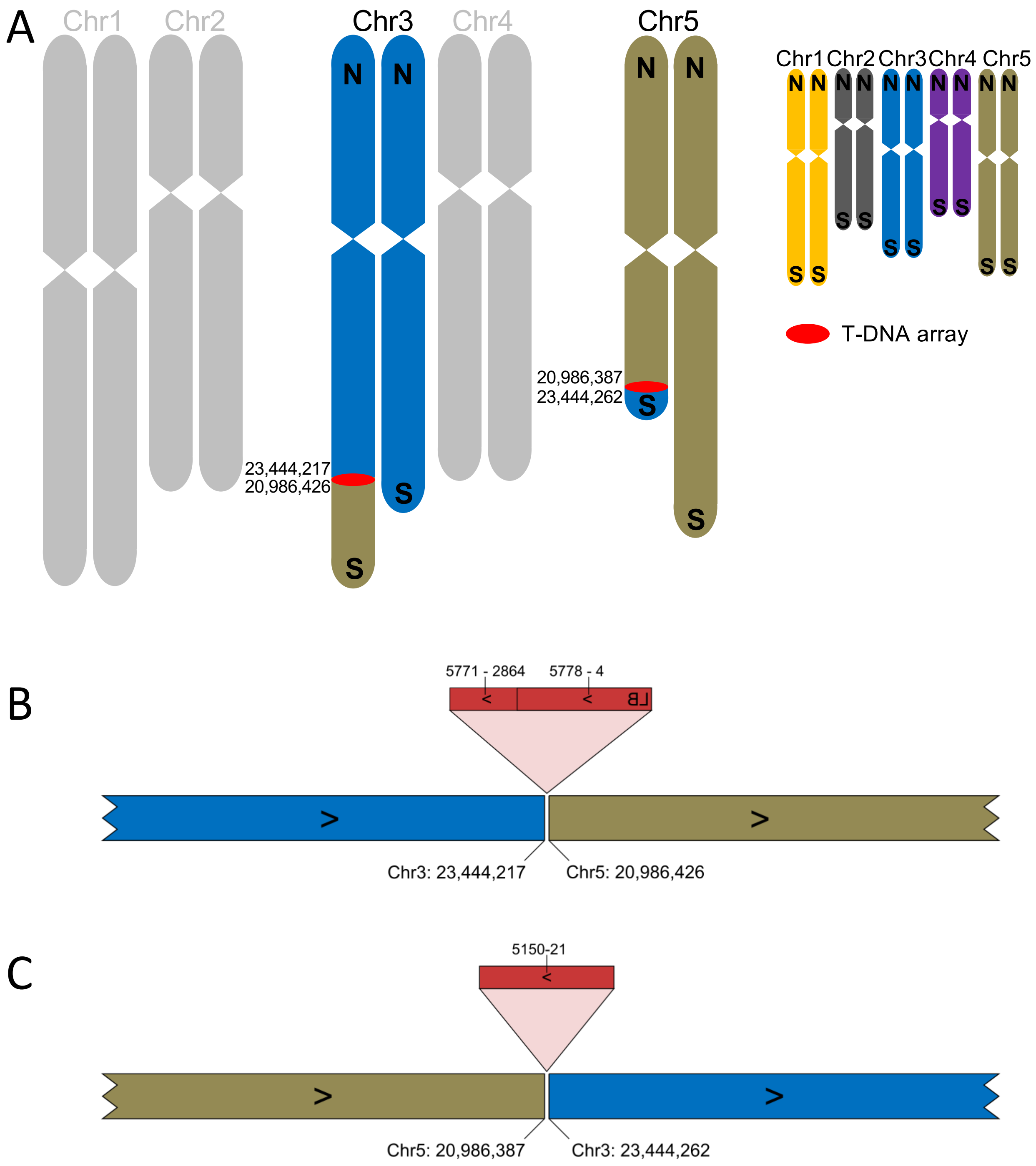
A
Chr1
Chr2
Chr3
Chr4
Chr5
N
S
23,444,217
20,986,426
20,986,387
23,444,262
Chr1 Chr2 Chr3 Chr4 Chr5
T-DNA array
B
5771 - 2864
5778 - 4
LB
Chr3: 23,444,217
Chr5: 20,986,426
C
5150-21
Chr5: 20,986,387
Chr3: 23,444,262

# GK-290G05

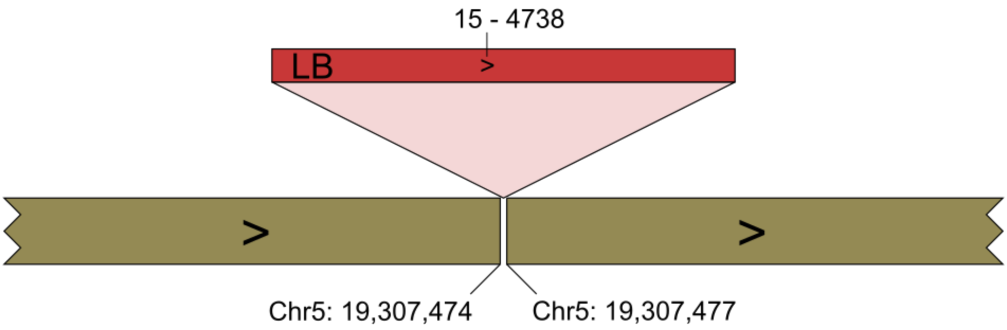

# GK-399C06

A

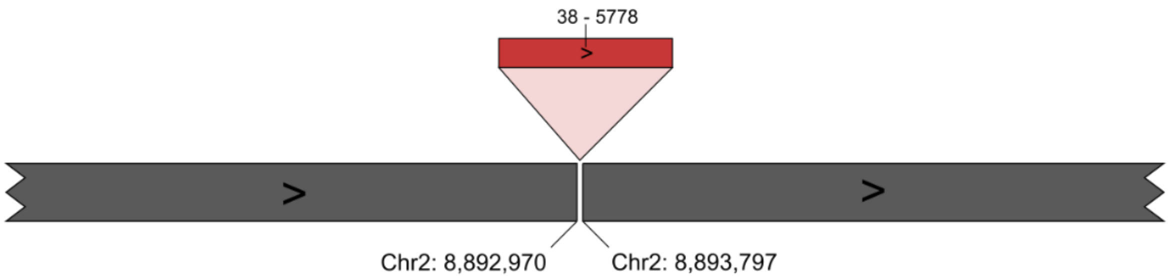

38 - 5778
>
>
>
Chr2: 8,892,970
Chr2: 8,893,797


B

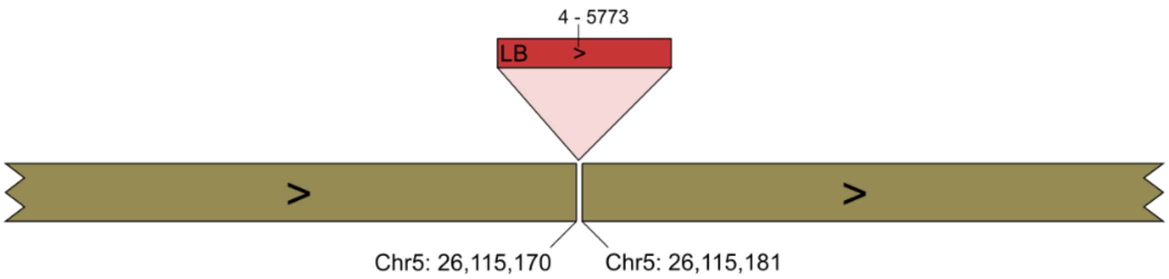

4 - 5773
LB
>
>
>
Chr5: 26,115,170
Chr5: 26,115,181

# GK-410B07

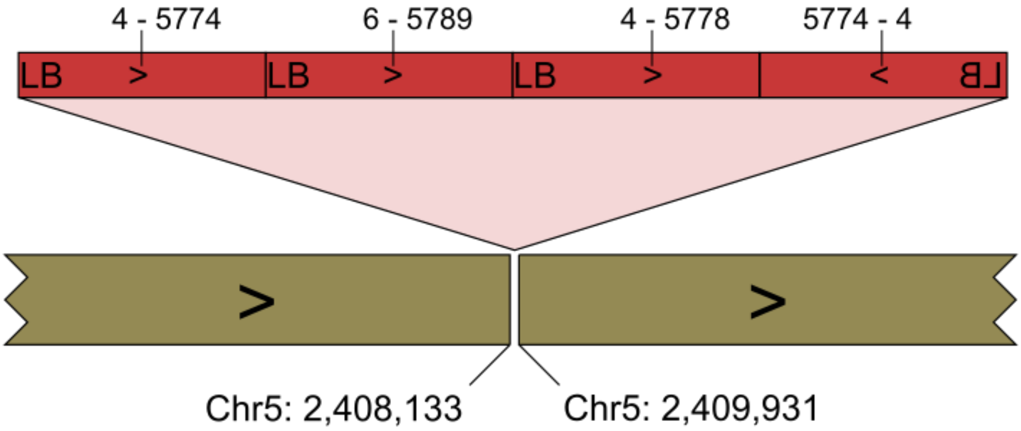

# GK-430F05

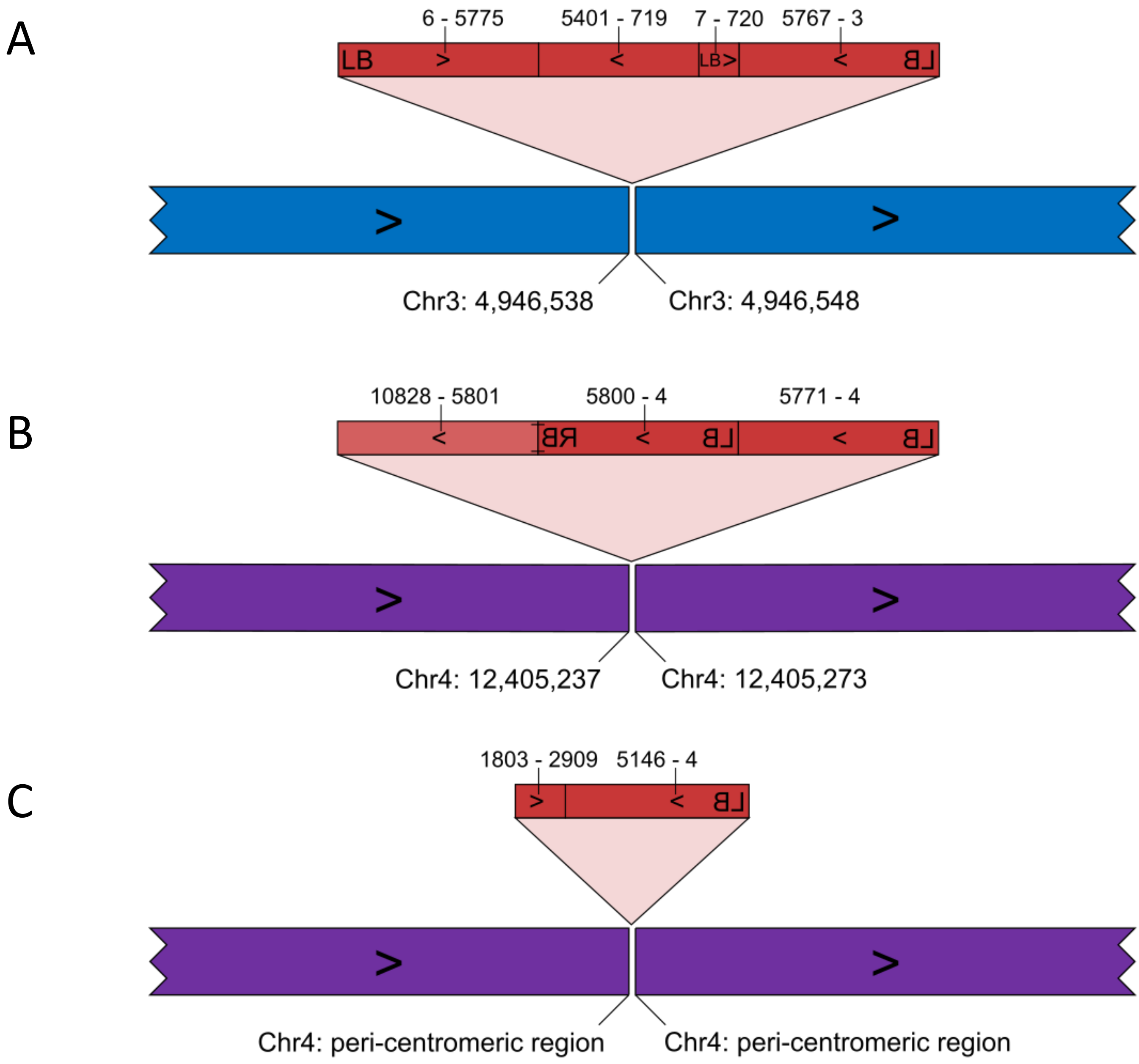
A
6 - 5775
5401 - 719
7 - 720
5767 - 3
LB
LB
Chr3: 4,946,538
Chr3: 4,946,548
B
10828 - 5801
5800 - 4
5771 - 4
Chr4: 12,405,237
Chr4: 12,405,273
C
1803 - 2909
5146 - 4
Chr4: peri-centromeric region
Chr4: peri-centromeric region

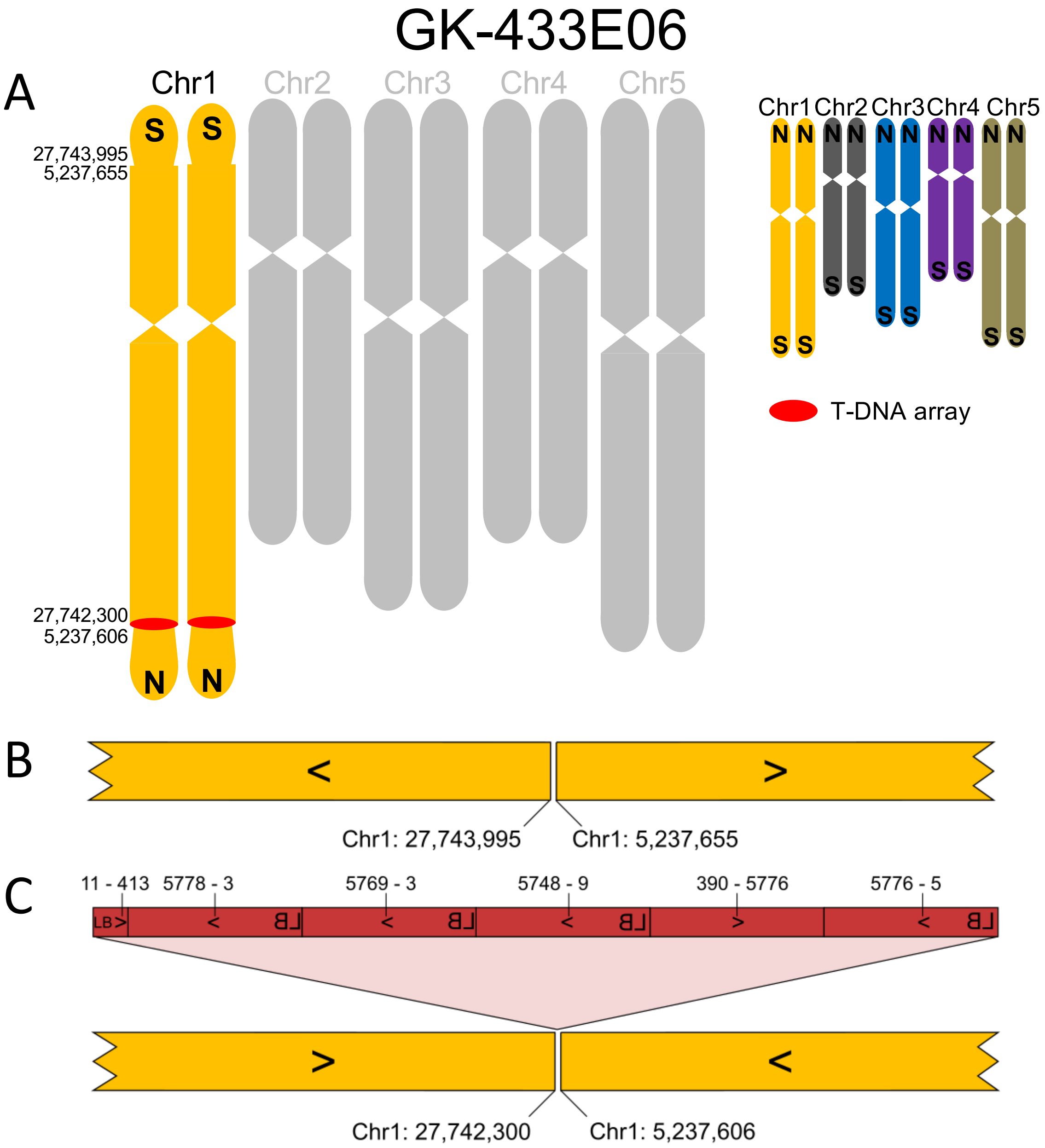
GK-433E06
A
Chr1
Chr2
Chr3
Chr4
Chr5
S
S
27,743,995
5,237,655
27,742,300
5,237,606
N
N
Chr1 Chr2 Chr3 Chr4 Chr5
T-DNA array
B
Chr1: 27,743,995
Chr1: 5,237,655
C
11 - 413
5778 - 3
5769 - 3
5748 - 9
390 - 5776
5776 - 5
LB
Chr1: 27,742,300
Chr1: 5,237,606

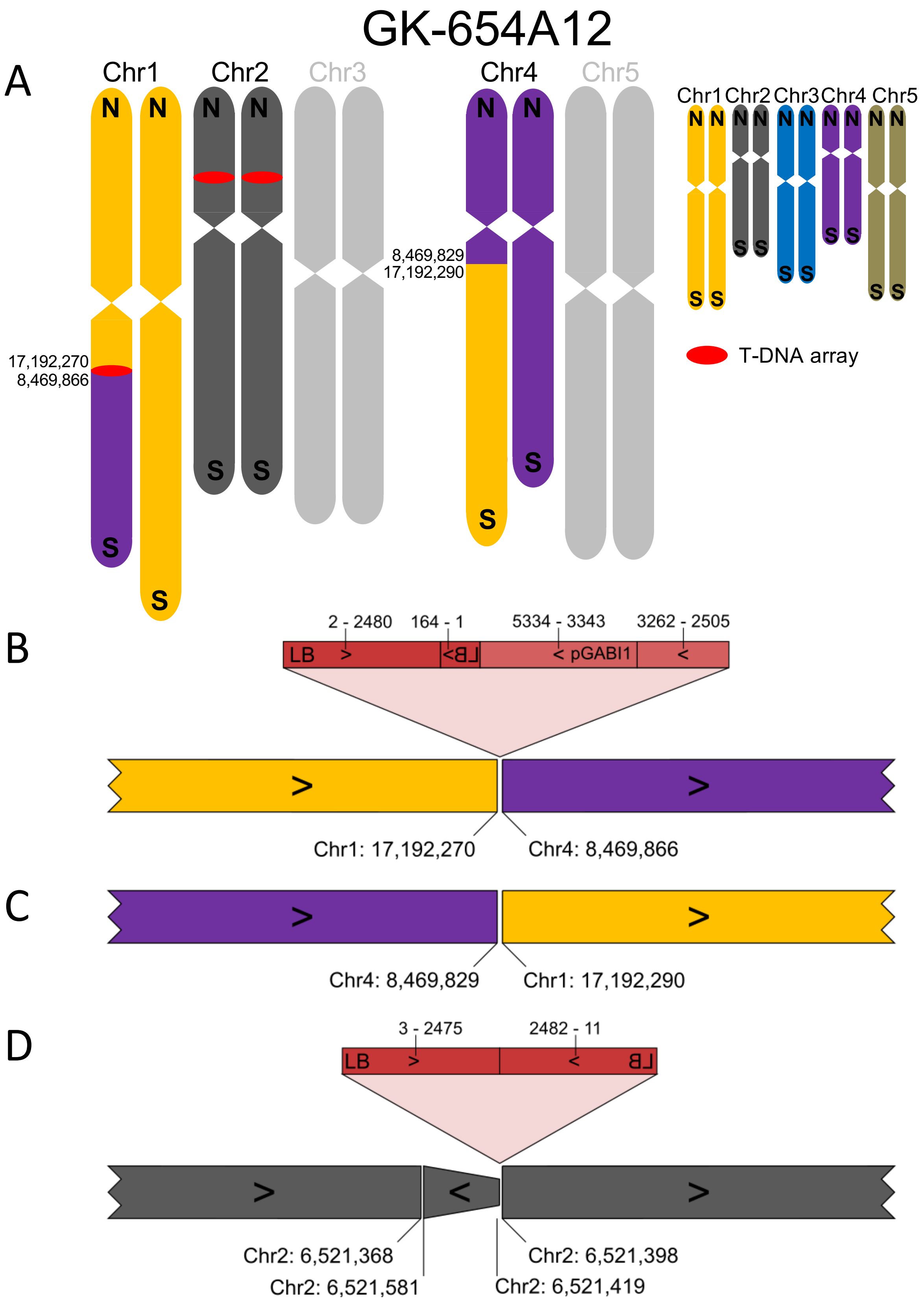
GK-654A12
A
Chr1
Chr2
Chr3
Chr4
Chr5
N
S
17,192,270
8,469,866
8,469,829
17,192,290
T-DNA array
B
2 - 2480
164 - 1
5334 - 3343
3262 - 2505
LB
pGABI1
Chr1: 17,192,270
Chr4: 8,469,866
C
Chr4: 8,469,829
Chr1: 17,192,290
D
3 - 2475
2482 - 11
LB
Chr2: 6,521,368
Chr2: 6,521,581
Chr2: 6,521,398
Chr2: 6,521,419

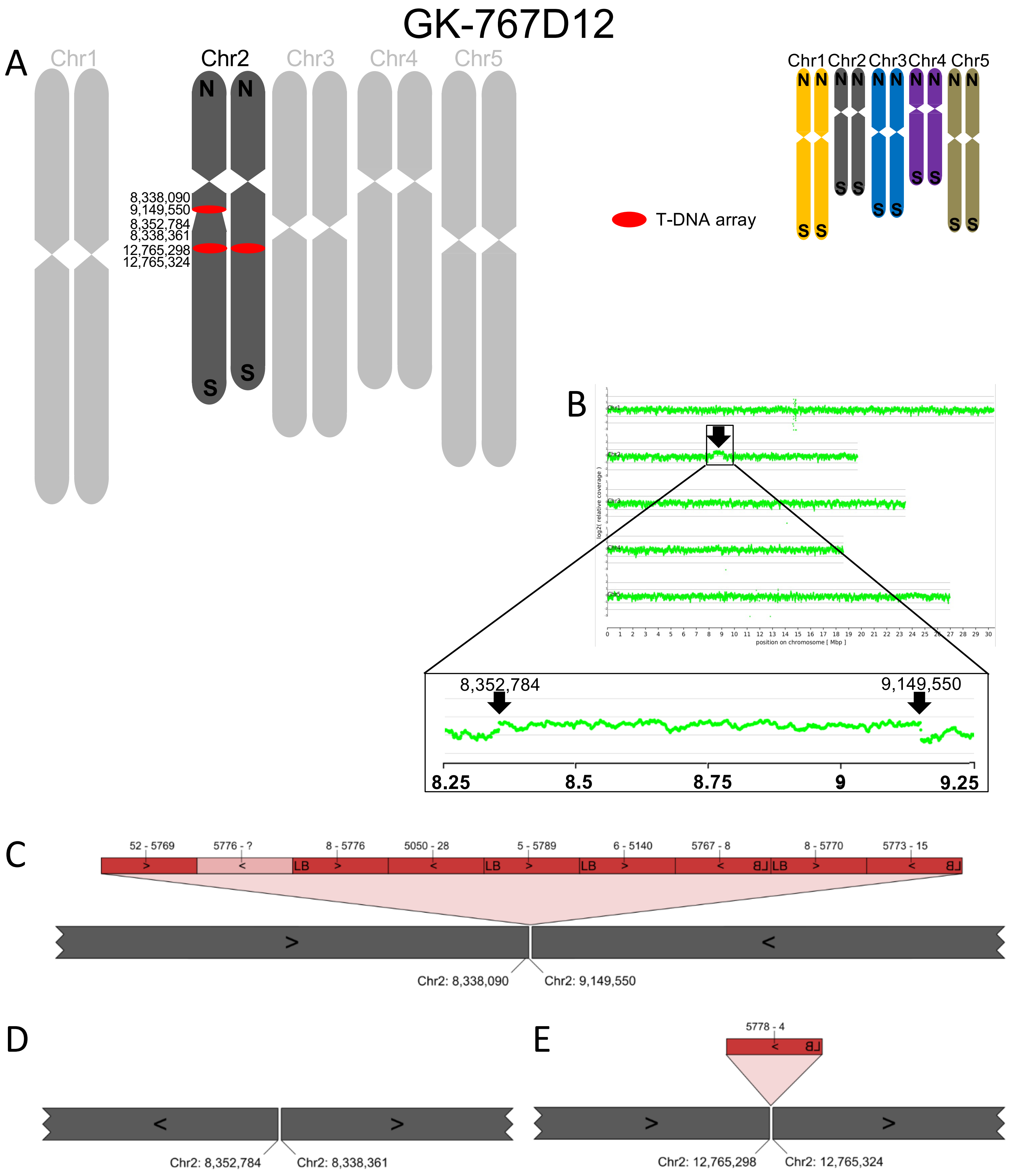
GK-767D12
A
Chr1
Chr2
Chr3
Chr4
Chr5
8,338,090
9,149,550
8,352,784
8,338,361
12,765,298
12,765,324
T-DNA array
B
8,352,784
9,149,550
8.25
8.5
8.75
9
9.25
C
52 - 5769
5776 - ?
8 - 5776
5050 - 28
5 - 5789
6 - 5140
5767 - 8
8 - 5770
5773 - 15
Chr2: 8,338,090
Chr2: 9,149,550
D
Chr2: 8,352,784
Chr2: 8,338,361
E
5778 - 4
Chr2: 12,765,298
Chr2: 12,765,324

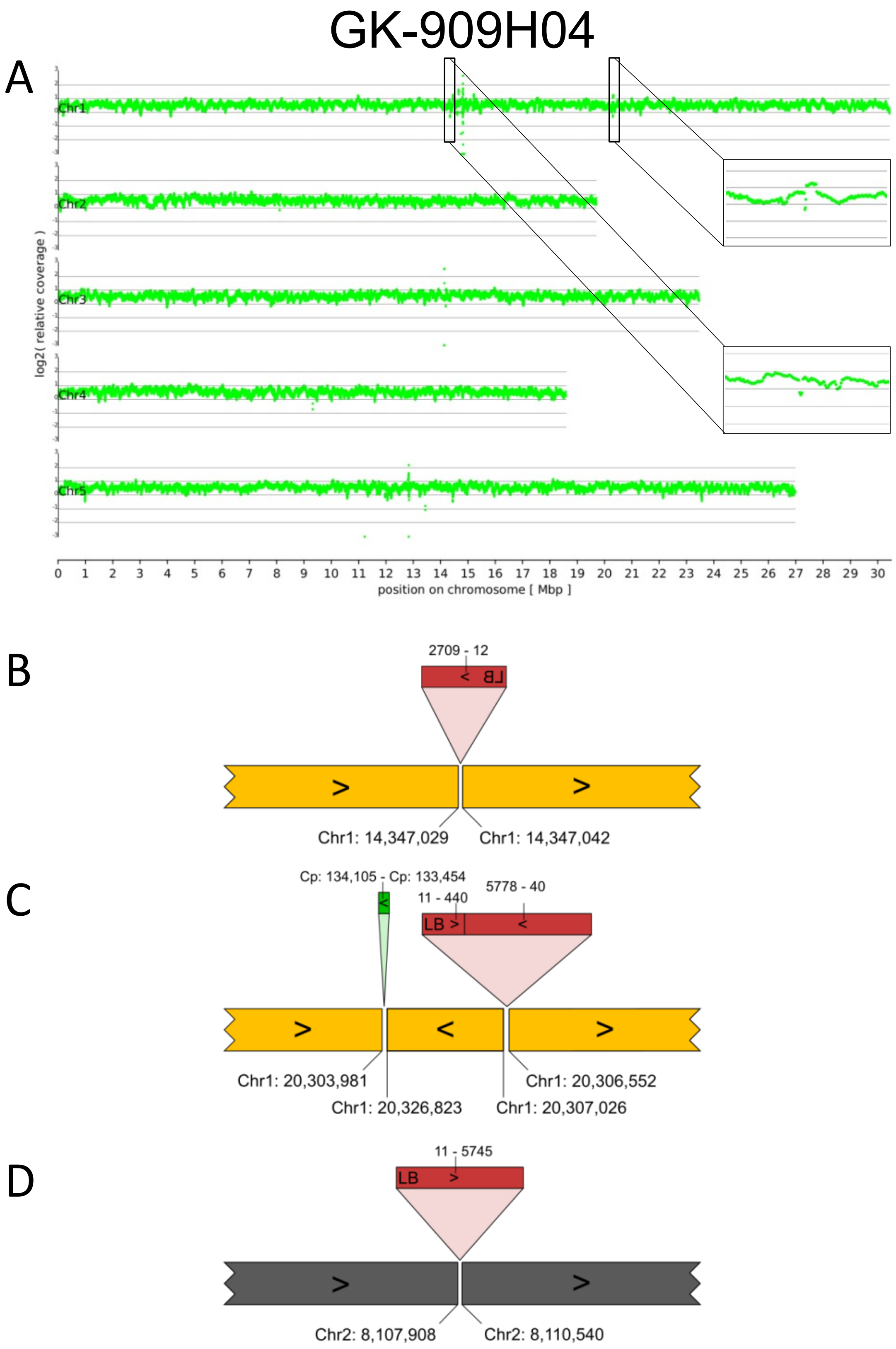
GK-909H04
A
Chr1
Chr2
Chr3
Chr4
Chr5
log2( relative coverage )
0 1 2 3 4 5 6 7 8 9 10 11 12 13 14 15 16 17 18 19 20 21 22 23 24 25 26 27 28 29 30
position on chromosome [ Mbp ]
B
2709 - 12
Chr1: 14,347,029
Chr1: 14,347,042
C
Cp: 134,105 - Cp: 133,454
11 - 440
5778 - 40
LB
Chr1: 20,303,981
Chr1: 20,326,823
Chr1: 20,306,552
Chr1: 20,307,026
D
11 - 5745
LB
Chr2: 8,107,908
Chr2: 8,110,540

# GK-947B06

A

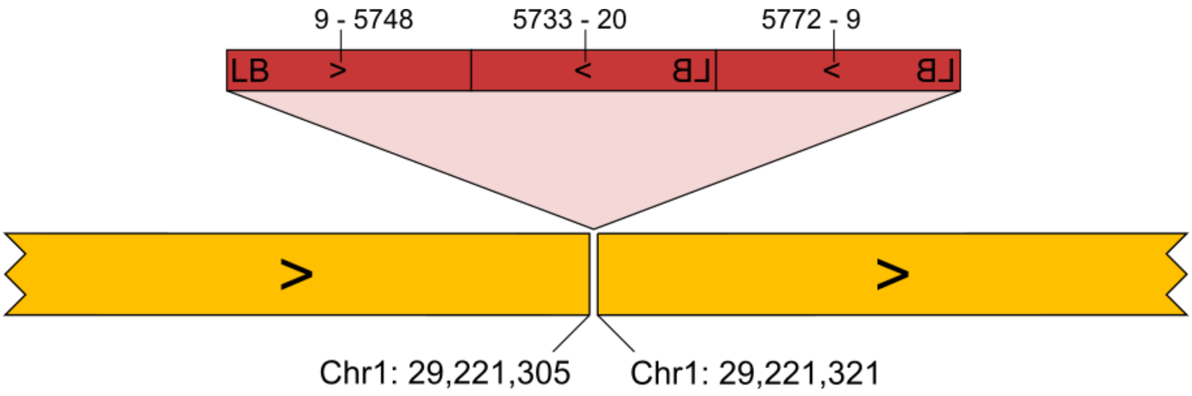
9 - 5748
5733 - 20
5772 - 9
LB
Chr1: 29,221,305
Chr1: 29,221,321

B

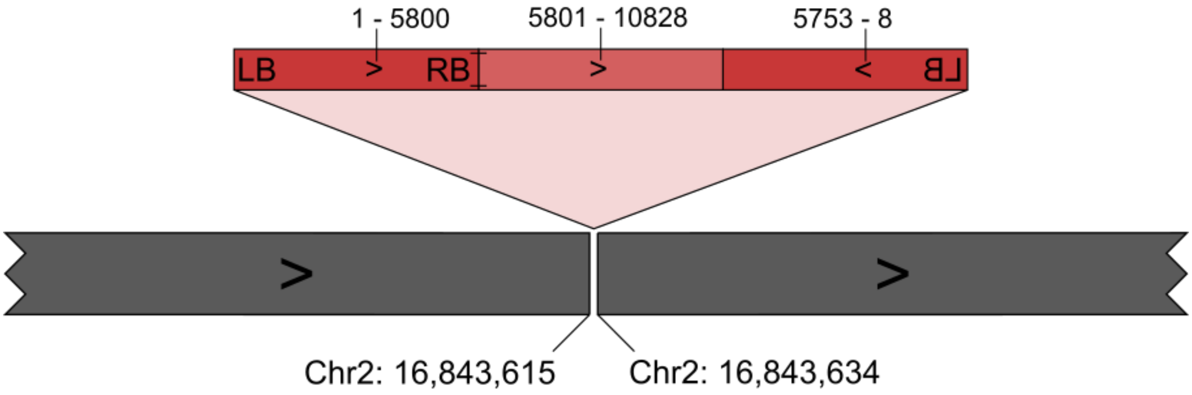
1 - 5800
5801 - 10828
5753 - 8
LB
RB
Chr2: 16,843,615
Chr2: 16,843,634
