## Additional file 9 for "Large scale genomic rearrangements in selected *Arabidopsis thaliana* T-DNA lines are caused by T-DNA insertion mutagenesis"

**Additional file 9: Assembly statistics of Col-0_GK-wt**

**Assembly statistics of the Col-0_GKat-wt assembly (GCA_905067165)**

number of contigs:	35

average contig length [bp]:	3,545,375

minimal contig length [bp]:	106,611

maximal contig length [bp]:	16,157,556

total number of bases:	124,088,133

GC content [%]:	36.43

N25 [bp]:	15,112,119

N50 [bp]:	14,300,866

N75 [bp]:	11,323,513

N90 [bp]:	4,269,132
