## Additional file 11 for "Large scale genomic rearrangements in selected *Arabidopsis thaliana* T-DNA lines are caused by T-DNA insertion mutagenesis"

Additional file 11: Dot plots between TAIR9 and Col-0_GK-wt for potential errors in the reference sequence.

These figures were generated based on a previously described script (Pucker et al. (2019), PLoS-One 14:e0216233). The intensity of blue coloration indicates the sequence similarity of BLAST hits. Structural variants between both genome sequences are revealed by dots that are not located on the expected central diagonal line from lower left to upper right. Titles of individual figures indicate the displayed region of the TAIR9 reference sequence, the AGIs of effected genes, and the IDs of overlapping BACs.

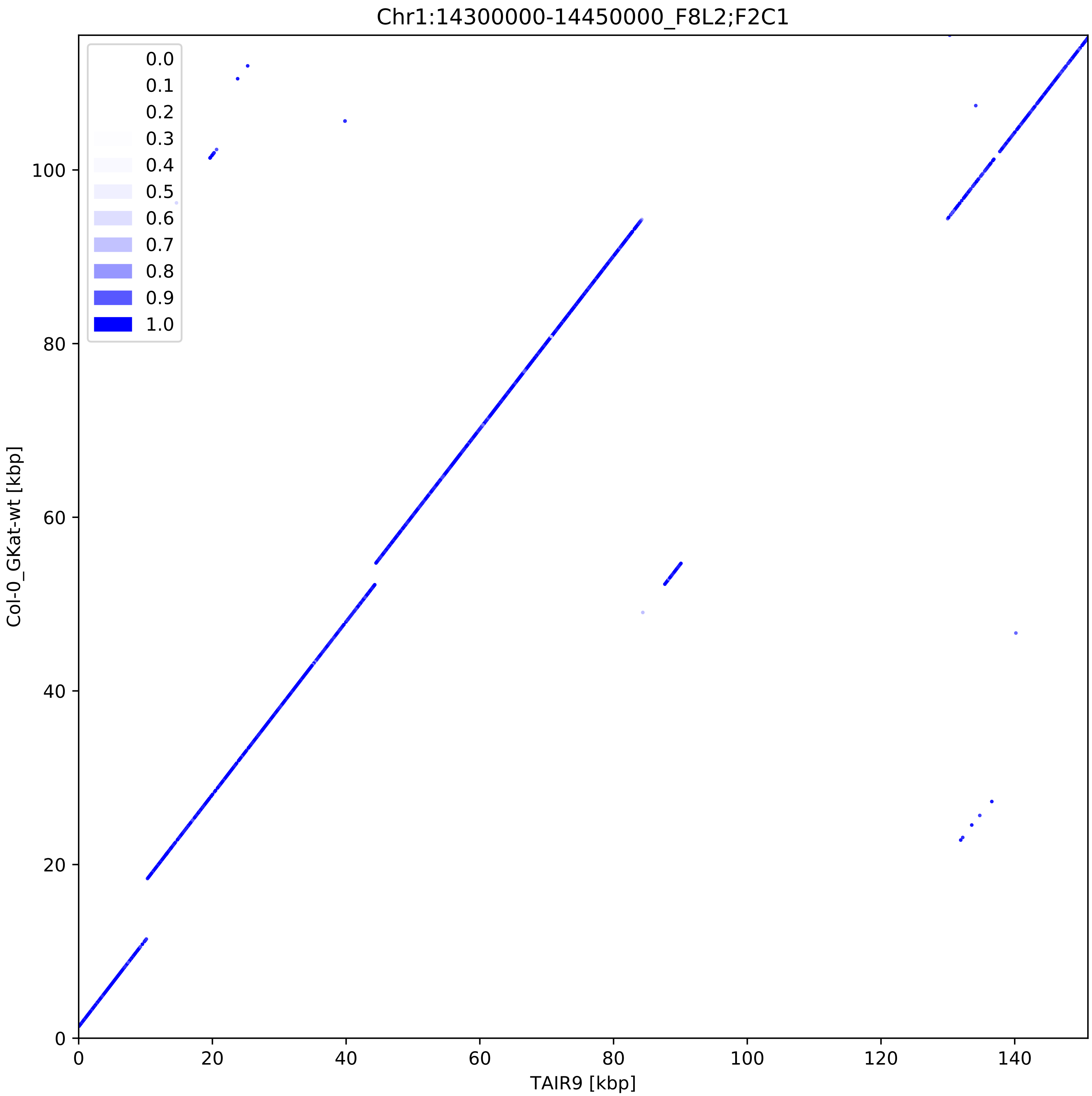

Chr1:14300000-14450000_F8L2;F2C1
0.0
0.1
0.2
0.3
0.4
0.5
0.6
0.7
0.8
0.9
1.0
Col-0_GKat-wt [kbp]
0
20
40
60
80
100
TAIR9 [kbp]
0
20
40
60
80
100
120
140

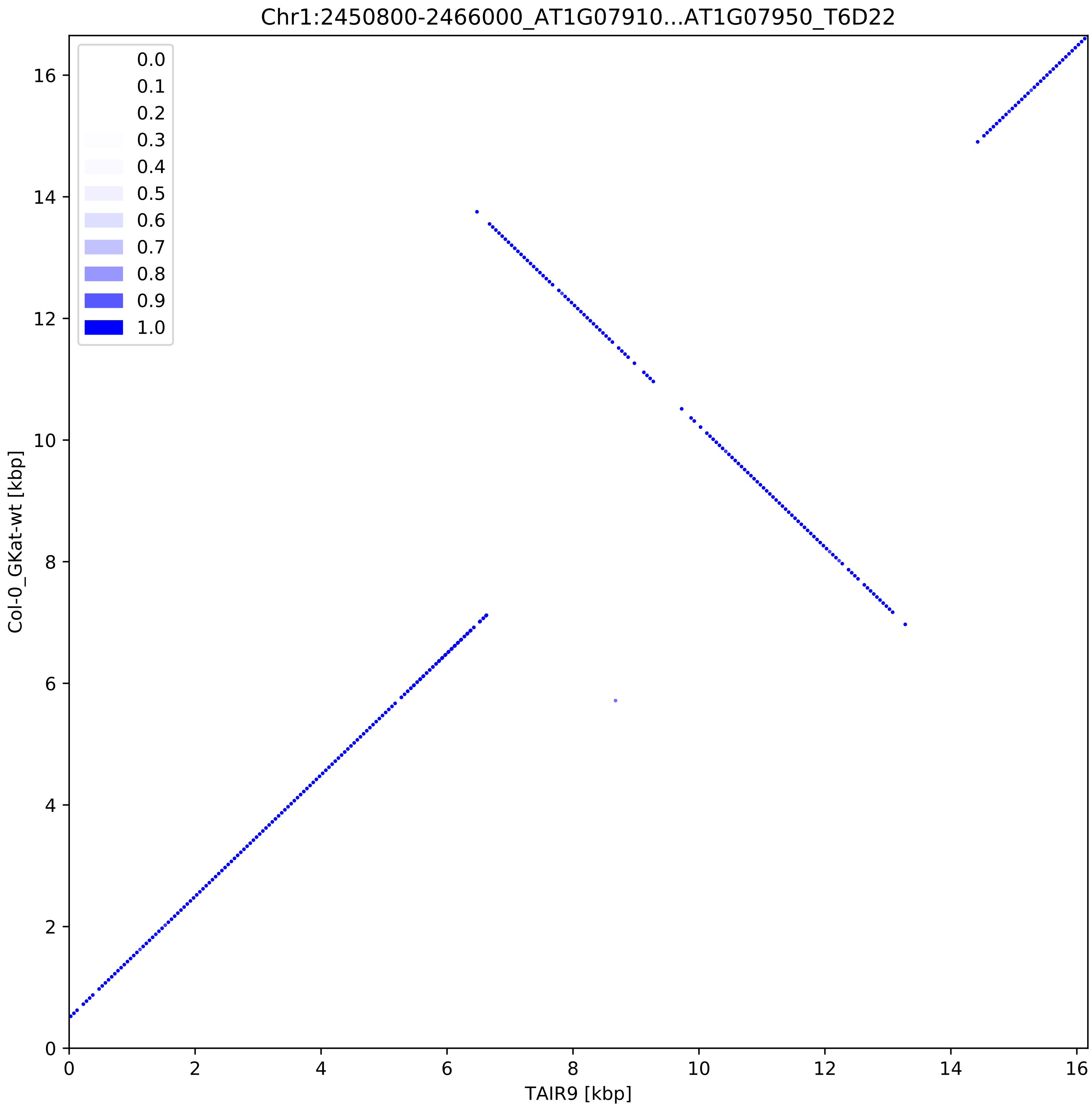
Chr1:2450800-2466000_AT1G07910...AT1G07950_T6D22
0.0
0.1
0.2
0.3
0.4
0.5
0.6
0.7
0.8
0.9
1.0
Col-0_GKat-wt [kbp]
TAIR9 [kbp]
0
2
4
6
8
10
12
14
16

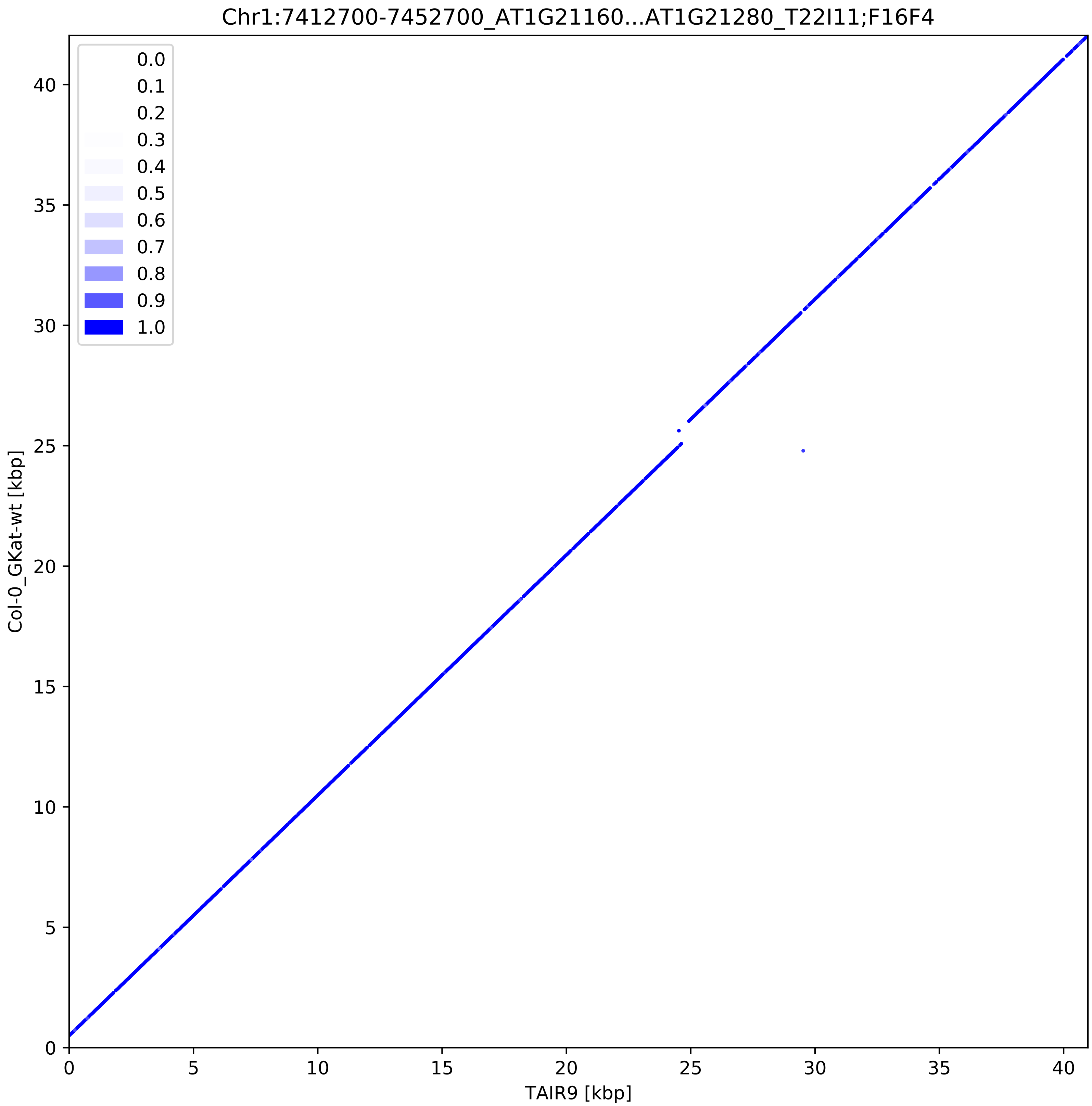
Chr1:7412700-7452700_AT1G21160...AT1G21280_T22I11;F16F4
0.0
0.1
0.2
0.3
0.4
0.5
0.6
0.7
0.8
0.9
1.0
Col-0_GKat-wt [kbp]
TAIR9 [kbp]
0
5
10
15
20
25
30
35
40

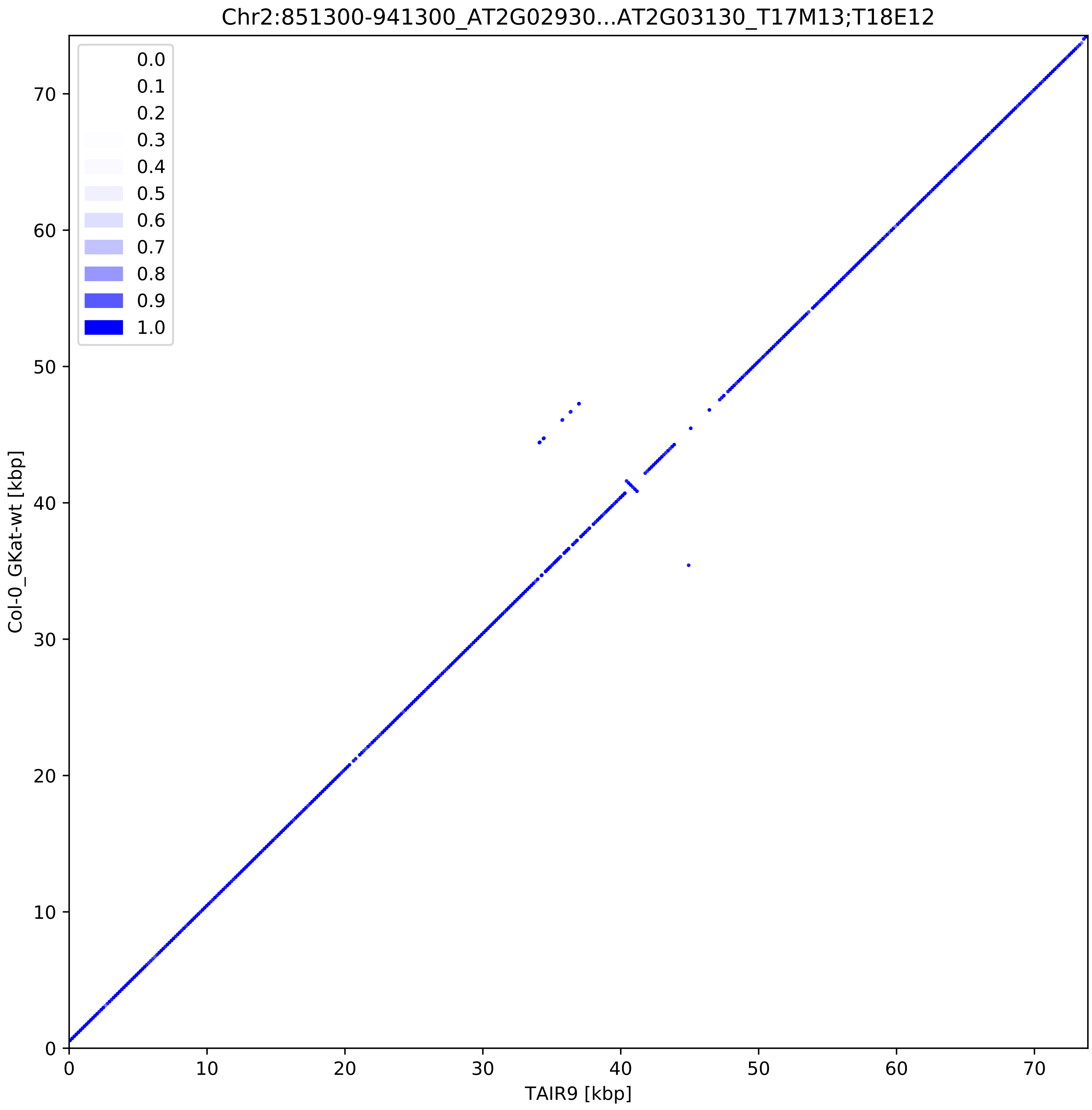
Chr2:851300-941300_AT2G02930...AT2G03130_T17M13;T18E12
0.0
0.1
0.2
0.3
0.4
0.5
0.6
0.7
0.8
0.9
1.0
Col-0_GKat-wt [kbp]
TAIR9 [kbp]
0
10
20
30
40
50
60
70

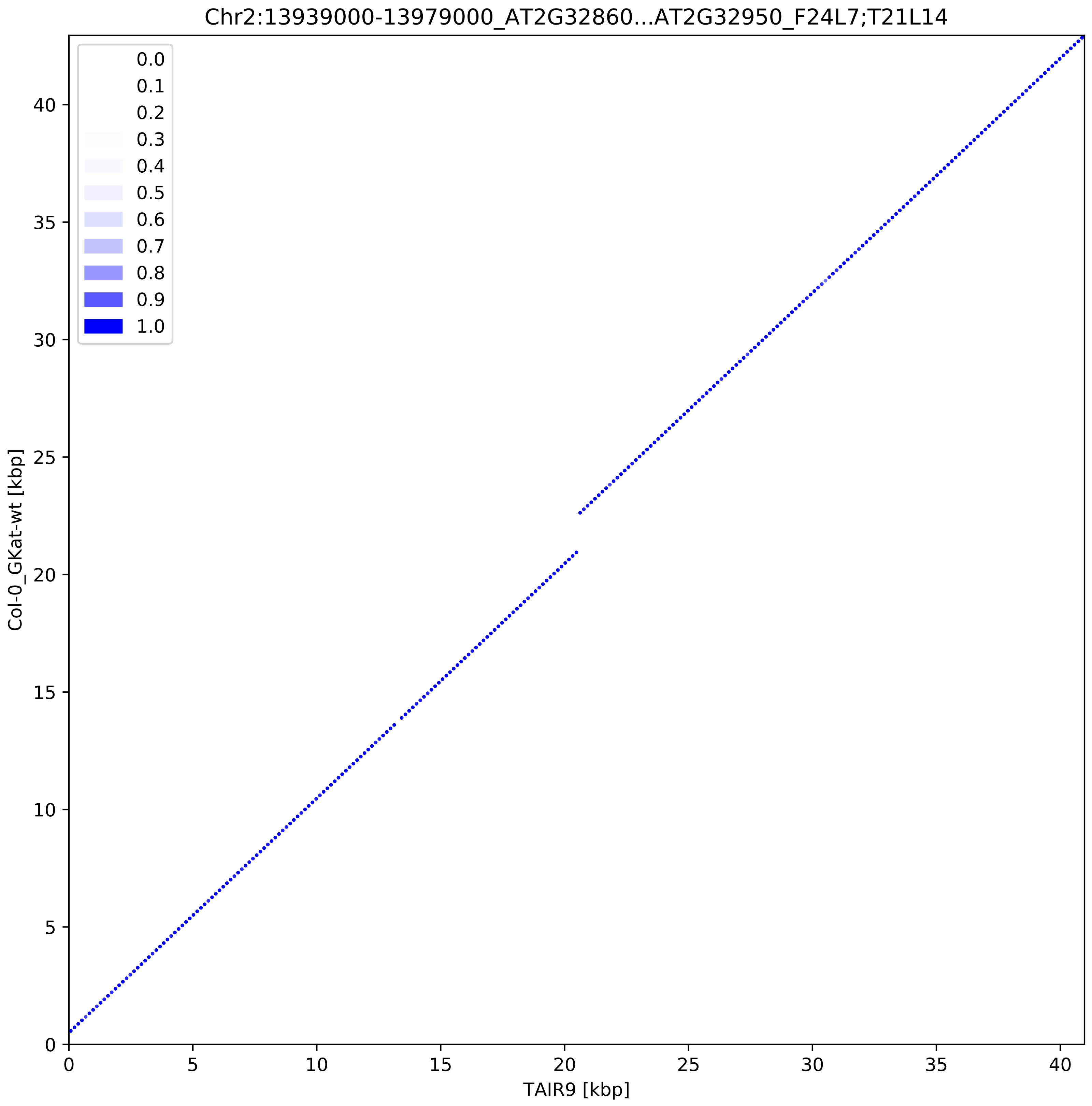
Chr2:13939000-13979000_AT2G32860...AT2G32950_F24L7;T21L14
0.0
0.1
0.2
0.3
0.4
0.5
0.6
0.7
0.8
0.9
1.0
Col-0_GKat-wt [kbp]
TAIR9 [kbp]
0
5
10
15
20
25
30
35
40

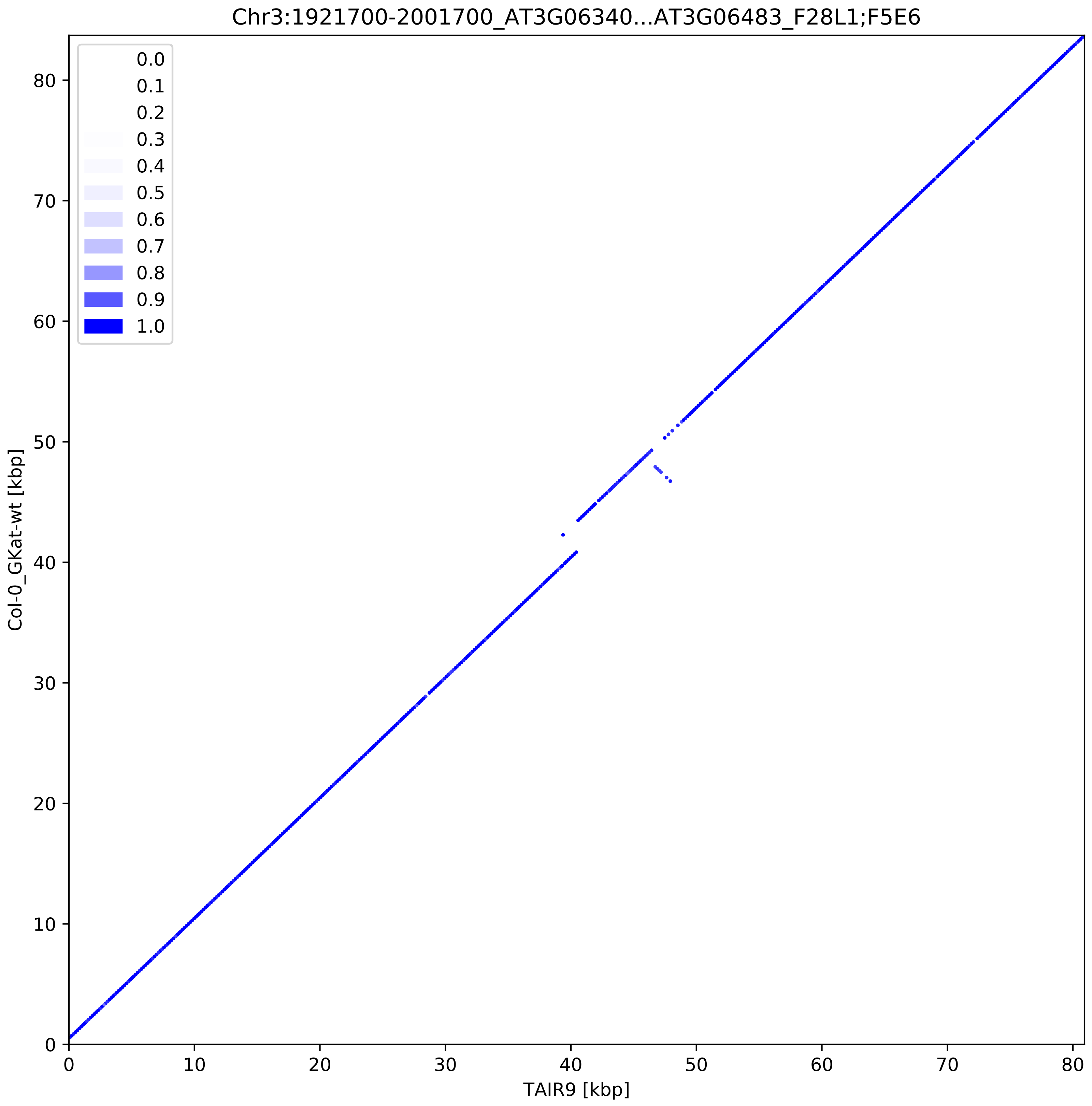
Chr3:1921700-2001700_AT3G06340...AT3G06483_F28L1;F5E6
0.0
0.1
0.2
0.3
0.4
0.5
0.6
0.7
0.8
0.9
1.0
Col-0_GKat-wt [kbp]
TAIR9 [kbp]
0
10
20
30
40
50
60
70
80

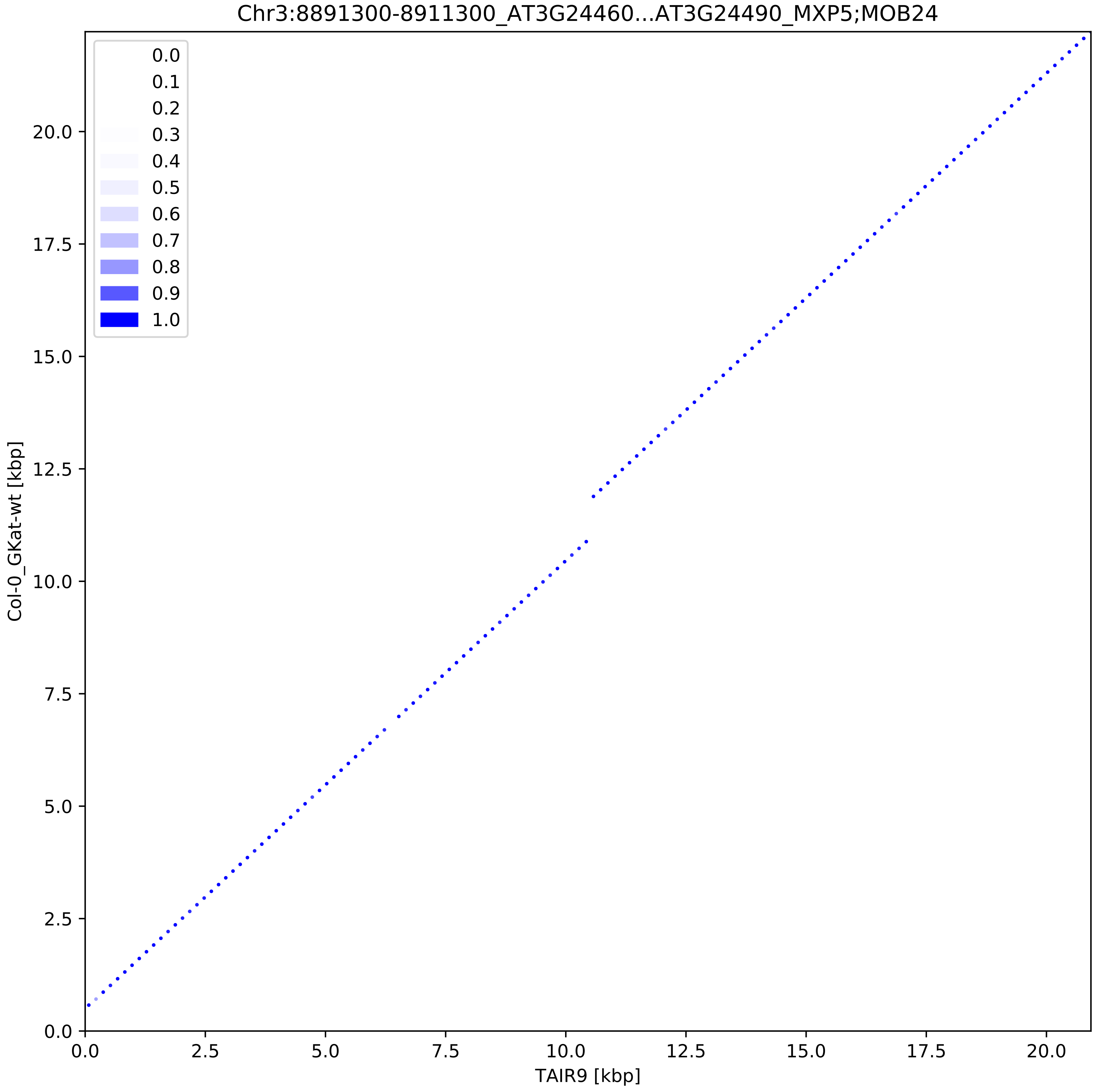

Chr3:8891300-8911300_AT3G24460...AT3G24490_MXP5;MOB24
0.0
0.1
0.2
0.3
0.4
0.5
0.6
0.7
0.8
0.9
1.0
Col-0_GKat-wt [kbp]
TAIR9 [kbp]
0.0
2.5
5.0
7.5
10.0
12.5
15.0
17.5
20.0

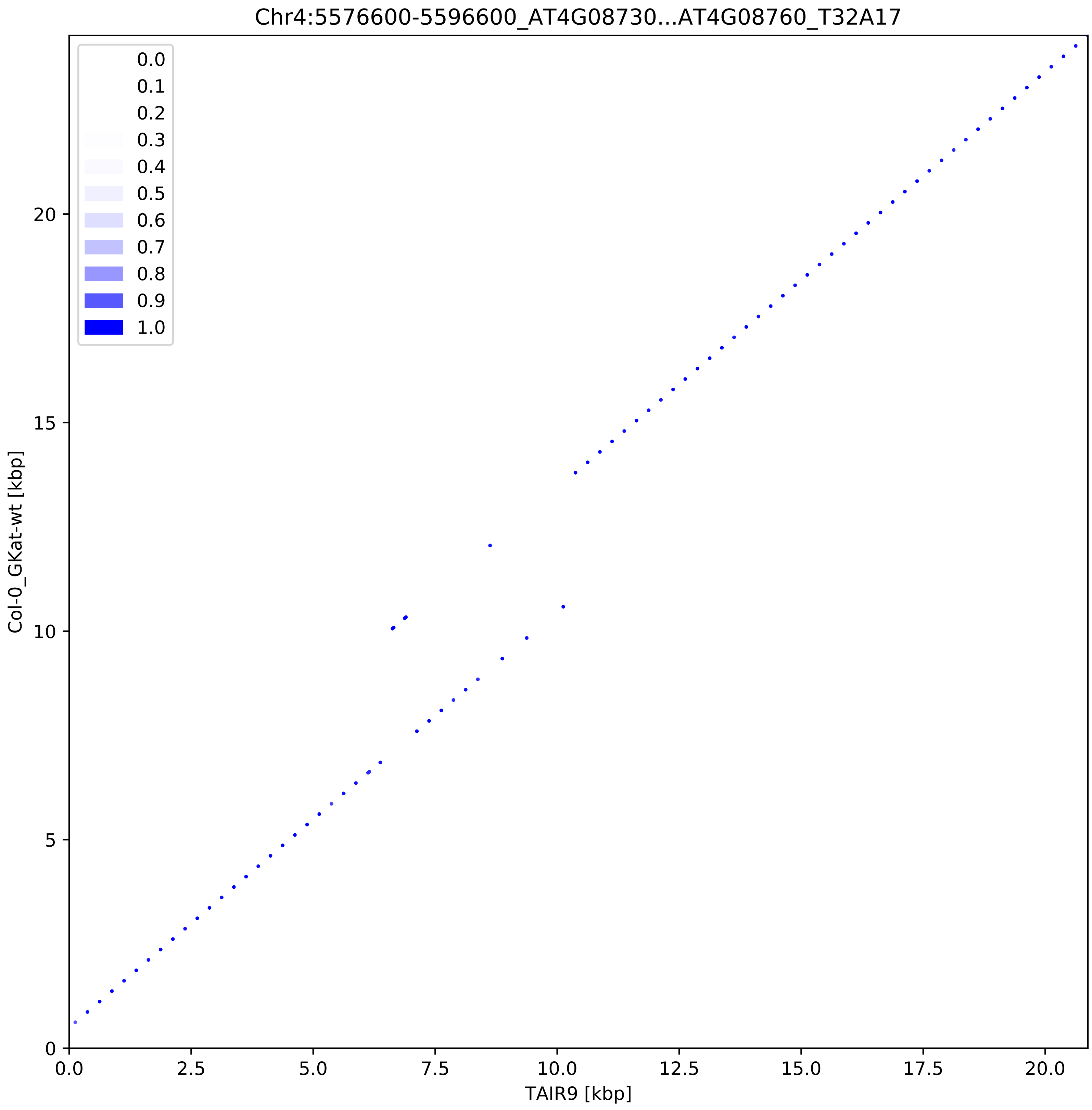
Chr4:5576600-5596600_AT4G08730...AT4G08760_T32A17
0.0
0.1
0.2
0.3
0.4
0.5
0.6
0.7
0.8
0.9
1.0
Col-0_GKat-wt [kbp]
TAIR9 [kbp]
0
5
10
15
20
0.0
2.5
5.0
7.5
10.0
12.5
15.0
17.5
20.0

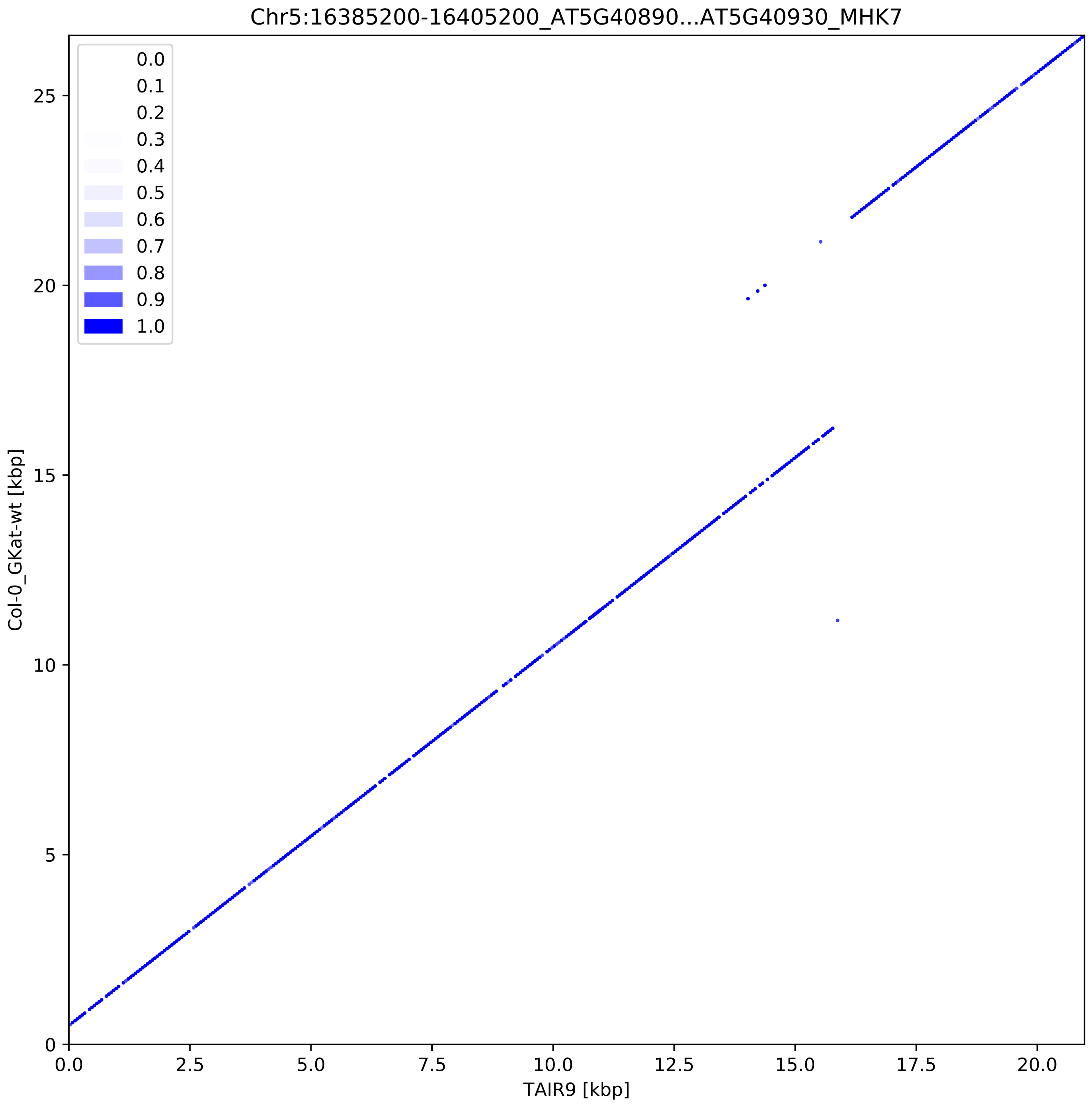

Chr5:16385200-16405200_AT5G40890...AT5G40930_MHK7
0.0
0.1
0.2
0.3
0.4
0.5
0.6
0.7
0.8
0.9
1.0
Col-0_GKat-wt [kbp]
TAIR9 [kbp]
0
5
10
15
20
25
0.0
2.5
5.0
7.5
10.0
12.5
15.0
17.5
20.0

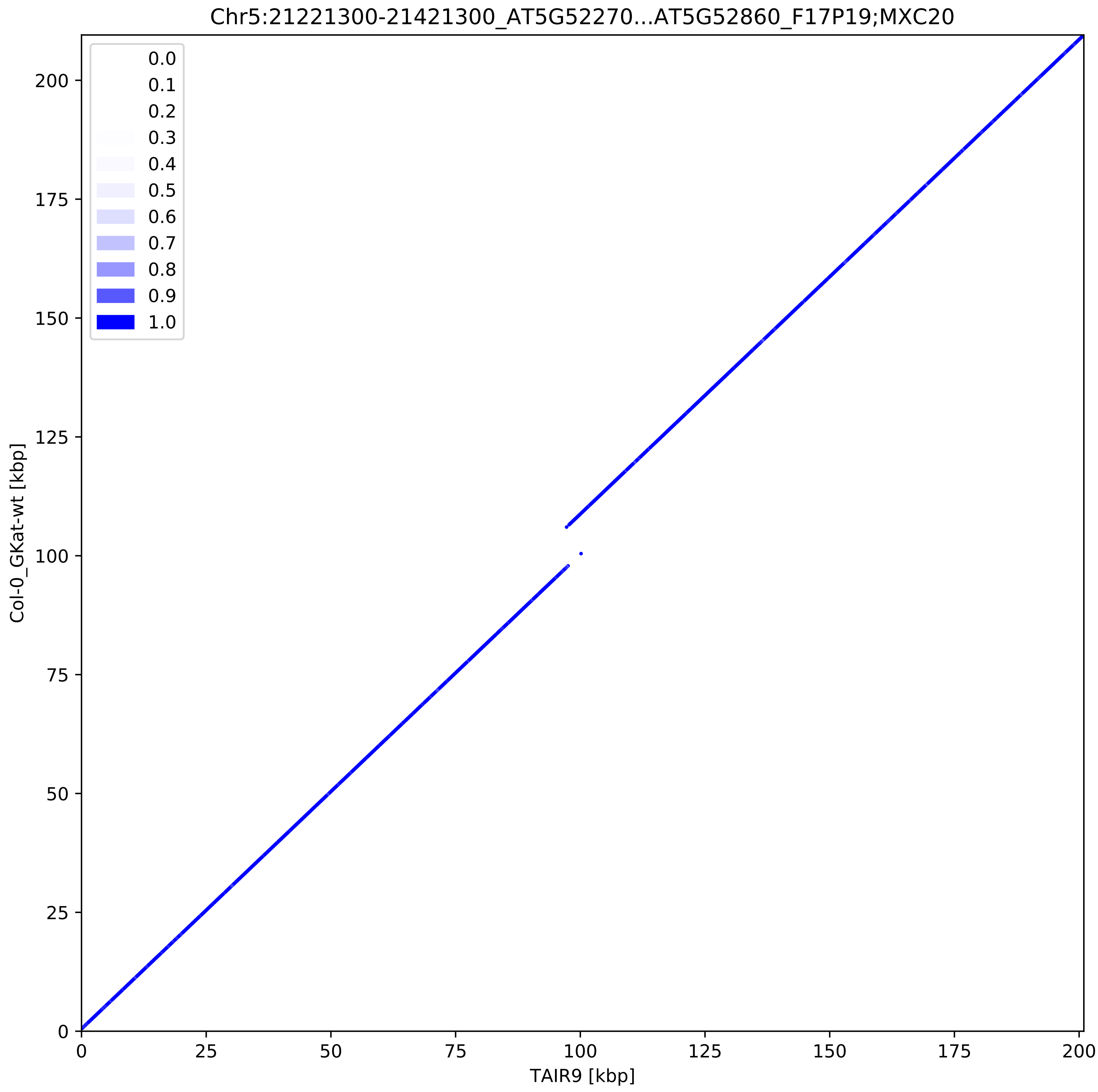
Chr5:21221300-21421300_AT5G52270...AT5G52860_F17P19;MXC20
0.0
0.1
0.2
0.3
0.4
0.5
0.6
0.7
0.8
0.9
1.0
Col-0_GKat-wt [kbp]
TAIR9 [kbp]
0
25
50
75
100
125
150
175
200
