## Additional file 12 for "Large scale genomic rearrangements in selected *Arabidopsis thaliana* T-DNA lines are caused by T-DNA insertion mutagenesis"

**Additional file 12: Protocol for the extraction of genomic DNA from *A. thaliana* for ONT sequencing.**

**Extraction of genomic DNA from *Arabidopsis thaliana* for ONT sequencing**

This protocol is a modification of a CTAB-based DNA extraction protocol (Rosso et al. 2003, PMB 53:247-259). Seedlings, plantlets or young leaves can be used as input material. About 1 g is sufficient.

- preheat 5 ml of CTAB-1 buffer with 300 µL ß-ME in a 50 ml tube to 70-75 °C

- homogenize / grind material using mortar and pestle in liquid N_2_

- add homogenized material to CTAB-1 buffer with ß-ME and mix quickly by inverting

(it is crucial that the powder is completely re-suspended)

- incubate for 30 min at 70-75 °C

(longer is possible if multiple samples are processed in parallel)

- let cool to room temperature (takes 5-10 minutes)

- add 5 ml of dichloromethane and mix gently by inverting the reaction tube

- centrifuge for 30 min at >10,000 x g (room temperature)

- transfer upper phase (4-5 ml) to a new 50 ml tube

- add 10 ml CTAB-2

- incubate for 5 min at room temperature

- centrifuge for 30 min at >10,000 x g (room temperature)

- discard supernatant, keep sediment

- add 1 ml of 1 M NaCl solution and gently dissolve the pellet

- precipitate DNA by adding 1 ml of isopropanol

(this volume needs to match the volume of NaCl solution)

- incubate for 5 min (room temperature)

- centrifuge for 30 min at >10,000 x g (room temperature)

- discard supernatant

- wash once with 2 ml of 70% ethanol

(this should be a fresh solution)

- centrifuge briefly after removal of supernatant and remove remaining drop of ethanol

- dry sediment at room temperature (incubate for about 5 minutes)

- re-dissolve pellet in 100 µl of CTAB-TE (10 mM Tris, pH 8,0; 0,1 mM EDTA pH 8,0) via incubation over night at room temperature

**CTAB buffer 1**

2 % CTAB

100 mM Tris-HCl, pH 8.0

20 mM EDTA

1,4 M NaCl

0.25 % PVP (optional)

**CTAB buffer 2**

1.0 % CTAB

50 mM Tris-HCl, pH 8.0

10 mM EDTA

0.125 % PVP (optional)

**CTAB-TE**

10 mM Tris, pH 8.0

0.1 mM EDTA pH 8.0

7.5 µg/ml RNAse (needs to be completely DNase free)
